# An improved PDE6D inhibitor combines with Sildenafil to synergistically inhibit KRAS mutant cancer cell growth

**DOI:** 10.1101/2023.08.23.554263

**Authors:** Pelin Kaya, Elisabeth Schaffner-Reckinger, Ganesh babu Manoharan, Vladimir Vukic, Alexandros Kiriazis, Mirko Ledda, Maria Burgos, Karolina Pavic, Anthoula Gaigneaux, Enrico Glaab, Daniel Kwaku Abankwa

## Abstract

The trafficking chaperone PDE6D (or PDEƍ) was proposed as a surrogate target for K-Ras, leading to the development of a series of inhibitors that block its prenyl-binding pocket. These inhibitors suffered from low solubility and intracellular potency, preventing their clinical development.

Here we developed a highly soluble PDE6D inhibitor (PDE6Di), Deltaflexin3, which has the currently lowest off-target activity, as we demonstrate in dedicated assays. We further increased the K-Ras focus, by exploiting that PKG2-mediated phosphorylation of Ser181 lowers K-Ras binding to PDE6D. Thus, the combination of Deltaflexin3 with the approved PKG2-activator Sildenafil synergistically inhibits cell- and microtumor growth. However, the overall cancer survival of the high PDE6D/ low PKG2 target population is higher than of the group with the opposite signature. Our results therefore suggest re-examining the interplay between PDE6D and K-Ras in cancer, while recommending the development of PDE6Di that ’plug’, rather than ’stuff’ the hydrophobic pocket of PDE6D.

**Significance:** Combinations of a novel PDE6D inhibitor with Sildenafil synergistically focus the inhibition on K-Ras, however, survival data of the target population suggest an interplay of K-Ras and PDE6D that needs further exploration.

## Introduction

The highly mutated oncogene KRAS is one of the best-established cancer targets. Only recently have two KRAS-G12C inhibitors, sotorasib and adagrasib, been approved for the treatment of lung cancer ^1,2^. While other allele specific-, pan-Ras- and Ras-pathway inhibitors are under intense development ^3,4^, there is still a need to target Ras more profoundly from various angles.

Inhibition of Ras membrane targeting remains a promising strategy for inhibitor development ^5,6^. The trafficking chaperone PDE6D (or PDEƍ) has been proposed as a surrogate drug target in KRAS mutant cancers ^7^. PDE6D possesses a hydrophobic pocket, which can bind to one or even two prenyl-moieties, thus having a cargo spectrum that comprises farnesylated or geranylgeranylated Ras- and Rho-family proteins, as well as Rab proteins ^8,9^. Only proteins that are not in addition palmitoylated in the vicinity of the prenylated cysteine are accepted as cargo, making mono- and dual-palmitoylated N-Ras, K-Ras4A and H-Ras effectively worse cargo in cells than K-Ras4B (hereafter K-Ras) ^10^. Cargo affinity is critically modulated by the four residues upstream of the prenylated cysteine. Structure and sequence comparisons suggest that the two residues upstream of the prenylated cysteine cannot be large amino acids, like Lys, Arg or Glu ^8^. This stretch of four residues also comprises Ser181 at the C-terminus of K-Ras, which can be phosphorylated by PKG2 ^11^. Binding data of PDE6D to K-Ras with a S181E mutation suggest a reduced interaction when K-Ras is phosphorylated on Ser181 ^8^. K-Ras has only micromolar affinity to PDE6D, while another cargo the inositol phosphatase INPP5E, has a low nanomolar affinity ^8,12^. This has important consequences for their subcellular distribution. While K-Ras can be released in the perinuclear area by the allosteric release factor Arl2, which binds to PDE6D when GTP-bound ^13,14^, INPP5E is only dislodged by GTP-Arl3 inside the primary cilium ^12^.

The development of inhibitors that competitively bind to the prenyl-pocket of PDE6D was pioneered by the Waldmann group ^15^. However, their first two generations of PDE6D inhibitors (PDE6Di) Deltarasin and Deltazinone1 appeared to have off-target issues and poor metabolic stability, respectively ^7,16^. In addition, both compounds were ejected by the GTP-Arl2-dependent mechanism, similar to the natural PDE6D cargo. Only their third-generation inhibitors, the Deltasonamides, could withstand GTP-Arl2-mediated ejection, as they were highly optimized for sub-nanomolar affinity. However, these compounds appeared to have low cell penetration ^15^. In an attempt to optimize the pharmacological properties, the chemotype was switched from benzimidazole to pyridazinones, such as Deltazinone ^16^. This led to the development of low nanomolar inhibitors, such as candidate compound **99** that was pharmacokinetically evaluated in mice, without assessment of anti-tumorigenic activity ^17^. Hence, from these pioneering compounds, anti-tumor activity in vivo was only demonstrated with the first-generation compound Deltarasin ^7^. All three compound generations were mostly evaluated in KRAS-mutant pancreatic cancer cell lines, yet both Deltarasin and Deltasonamide were also micromolar active in KRAS mutant and PDE6D-dependent colorectal cancer cell lines ^18^.

Another class of more recent PDE6Di are proteolysis-targeting chimeras (PROTACs). Unlike classical competitive inhibitors they do not have to bind permanently i.e., they can act sub-stoichiometrically ^19^. Proof-of-concept PROTACs from two groups were developed based on previously established competitive PDE6Di, Deltasonamide and Deltazinone ^20,21^. These heterobifunctional compounds bind with their first functional moiety to the prenyl-pocket of PDE6D and with the second they recruit an E3 ubiquitin ligase complex to instruct proteasomal degradation of PDE6D. While the Deltasonamide-derived PROTAC effectively decreased PDE6D levels in pancreatic cancer cells ^20^, the Deltazinone-derived PROTAC was even efficacious in SW480 xenografts in mice ^21^.

Following the pioneering work of the Waldmann group, other PDE6D-pocket competitive inhibitors were investigated, although for several of them clear in vitro or cellular target engagement data are missing. However, the Sheng group developed compounds that bound to PDE6D in vitro with nanomolar affinity. Some suppressed MAPK-output, but again had only micromolar cellular activity ^22,23^. Interestingly, in their most recent work their spiro-cyclic compound **36l** (Kd = 127 nM) showed target engagement in cells, while also demonstrating in vivo efficacy in KRAS mutant primary cell lines ^24^. In another study, the triazole **27** had nanomolar activity in a PDE6D binding assay and robustly inhibited MAPK-output at 10 μM and A549 cell growth at this concentration range ^25^.

Another PDE6Di emerged from a Rac-inhibitor screen, which led to the oxadiazole DW0254 as a submicromolar active compound (Kd = 436 ± 6 nM) ^26^. This compound inhibited downstream signaling of Ras above 20 μM and in vivo activity was observed with pretreatment of transplanted T-cell cancer cells or application of a pump to the graft site, due to poor solubility.

We have previously published novel competitive PDE6Di called Deltaflexins, for which we determined low micromolar affinities in a dedicated surface plasmon resonance assay, that were matched by a similar level of activity in KRAS mutant HCT116 and MDA-MB-231 cancer cells ^27^. Their chemical design features a hexamethylene-amide-backbone, which allowed simple derivatization and compound evolution. Importantly, Deltaflexins demonstrated the expected K-Ras-over H-Ras-selectivity in cells, an important on-target feature.

A number of questions remain unresolved regarding PDE6D as a surrogate target for K-Ras. Current PDE6Di are still at the hit stage and have various problems, such as poor solubility, metabolic instability and off-target issues ^16,17^. This makes the interpretation of phenotypic data and validation of PDE6D as a drug target in vivo difficult ^7^. Together with the broad cargo spectrum of PDE6D, which involves far more prenylated proteins than K-Ras, it is almost impossible to tell in which cancer type PDE6Di should be applied. Hence, clear genetic determinants that could indicate a susceptibility to PDE6D inhibition are lacking.

Here, we established an in silico library of compounds by cross-hybridizing moieties of existing PDE6Di with our previous hexamethylene-amide-backbone ^27^. Aided by computational docking, we derived rationales for the synthesis of 16 novel PDE6Di, that we comprehensively characterized biochemically and in cells for potency and K-Ras- and PDE6D- on-target selectivity. We demonstrate that efficacy and more focused inhibition of K-Ras can be achieved by combining our most selective and highly soluble inhibitor **Deltaflexin3** synergistically with the clinically approved Sildenafil.

## Results

### Computational docking aided design of novel PDE6D inhibitors

We previously demonstrated that PDE6Di can be efficiently generated by using a hexamethylene-amide-backbone ^27^. Using this backbone as a base, we created an in silico library of hybrid compounds, which contained moieties of established PDE6Di, such as Deltarasin, Deltazinone1 and Deltasonamide1 that also served as references in this study (**Figure 1A**) ^7,15,16^.

**Figure 1.**
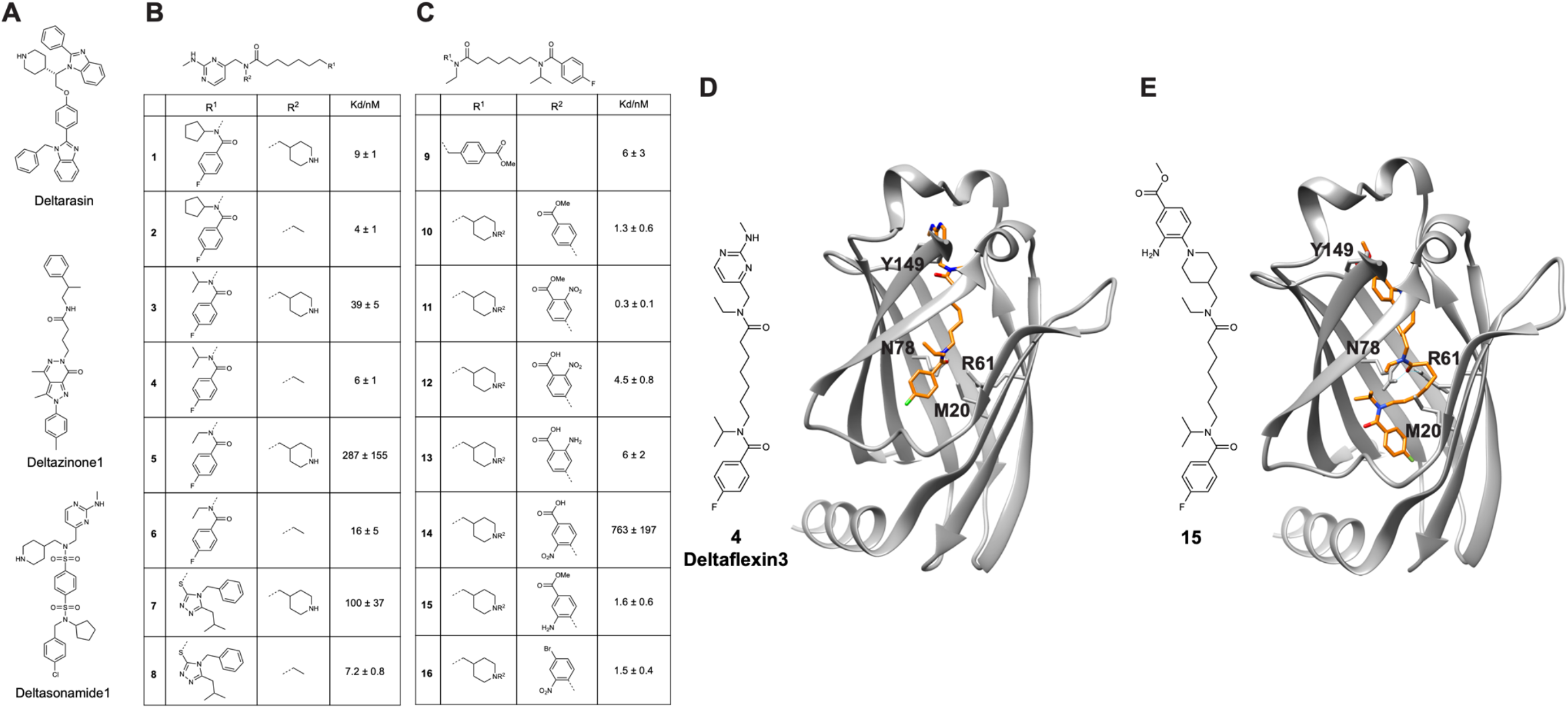
Investigated PDE6D inhibitors (PDE6Di) with affinities and computational docking. (**A**) Structures of employed reference PDE6Di. (**B,C**) Structures of developed first (B) and second (C) round PDE6Di with PDE6D dissociation constants measured using F-Ator in a fluorescence polarization assay; n ≥ 2. (**D,E**) Computational docking of compounds **4** (later named Deltaflexin3; D) and **15** (E) to PDE6D in the open state (PDB ID 4JV8).

Altogether, 313 compounds were thus designed in the first round and computationally docked to PDE6D (PDB ID 4JV8), using Glide docking software ^28^. Compounds selected based on the docking scores, MM-GBSA binding energy and visual inspection were prioritized and provided a rationale for the synthesis of a first round of eight compounds that were biochemically and cell-biologically characterized (**Figure 1B; Data S1 and S2**).

Subsequently, the best performing compound **4** was chosen as a starting point for derivatives that were again first evaluated by in silico docking using SeeSAR. In this second round, compounds were extended to attempt interactions with residues at the entry of the hydrophobic pocket of PDE6D. Based on these computational data a second round of eight candidate compounds was synthesized and characterized like the first-round compounds (**Figure 1C; Data S1 and S2)**.

Computational docking data of two of our compounds **4** and **15** revealed multiple van-der-Waals contacts to residues Met20, Arg61, Gln78, and Tyr149 (**Figure 1D,E**). Hydrogen bonds to these residues were only predicted for **15** with Arg61 and Gln78 (**Figure 1E**). The Arg61 hydrogen bond is shared with the reference inhibitors Deltarasin and Deltazinone1 ^7,16^.

### In vitro affinity and intracellular BRET-assays quantify target engagement and K-Ras-selectivity

All 16 compounds that were prioritized for synthesis first underwent in vitro testing using a previously employed fluorescence polarization assay where the FITC-labelled PDE6D-binder Atorvastatin (F-Ator) was used as a probe ^7^ (**Figure 1B,C; Data S2**). In addition, we determined the affinities of compounds using the FITC-labelled farnesylated peptide derived from the C-terminus of the small GTPase Rheb (F-Rheb) ^14^ (**Data S2**). When using F-Ator as a probe, we recovered affinities in the low nanomolar range for reference compounds, Deltarasin (Kd = 39 ± 15 nM), Deltazinone1 (Kd = 3.8 ± 0.4 nM) and Deltasonamide1 (Kd = 0.11 ± 0.03 nM), similar to previously published values ^7,15,16^. By contrast, affinities determined using F-Rheb were typically only in the sub-micromolar range (**Data S2**). However, both datasets overall correlated and served to rank the in vitro potencies of our 16 compounds and we will in the following refer to the values obtained with F-Ator, unless otherwise stated (**Figure 2A, Figure S 1A**).

**Figure 2.**
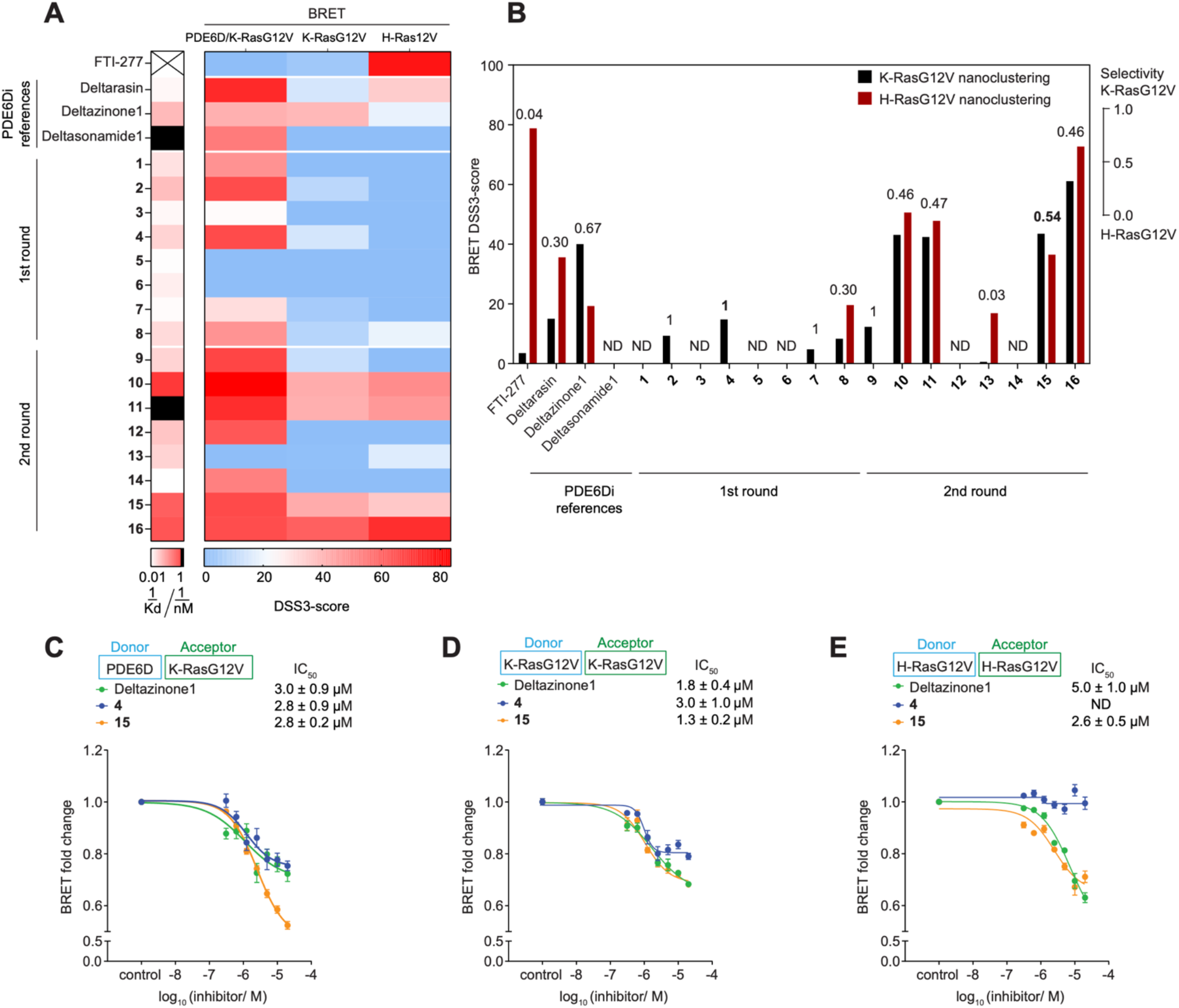
Quantification of on-target activity of PDE6Di in vitro and in cellular BRET-assays. (**A**) Heatmaps of in vitro affinity of compounds determined using F-Ator (first column; n ≥ 2) and DSS3-scores from cellular BRET-experiments. The disruption of the PDE6D/ K-RasG12V complex (second column; n ≥ 2) and of K-RasG12V- and H-RasG12V-membrane anchorage (third and fourth columns, respectively; n ≥ 2) were measured by BRET over a wider concentration range and the area under the curve DSS3-score was determined. (**B**) Quantification of K-RasG12V-selectivity (values above bars) was performed by determining the ratio of K-RasG12V and the sum of K-RasG12V- and H-RasG12V-BRET DSS3-scores from (A). (**C-E**) Dose-dependent change of normalized BRET-signals after treatment with indicated compounds using BRET donor/ acceptor-pairs shown on top; n ≥ 4.

Subsequently three cellular BRET (Bioluminescence Resonance Energy Transfer) assays were applied to profile the disruption of the PDE6D/ K-Ras interaction and loss of functional membrane organization of K-Ras as compared to H-Ras over a wider concentration range in HEK293-EBNA cells (**Figure 2A**). In analogy to our previous FRET-based target engagement assay ^27^, we implemented a BRET-assay with Rluc8-PDE6D and GFP2-K-RasG12V to determine the intracellular potency of compounds to displace K-RasG12V from PDE6D (**Figure 2A; Data S2**).

While intracellular IC50-values were in the micromolar regime (**Data S2**), we generally employed the more robust normalized area under the curve DSS3-score for dose-response data^29^. Overall, DSS3-scores from the PDE6D/ K-RasG12V-BRET correlated with in vitro affinities, and in both datasets, potencies increased markedly from the first to the second round of compounds (**Figure 2A**).

A second set of BRET-assays was likewise built in analogy to previous FRET-assays ^30,31^. We assessed the BRET that emerges between a Rluc8-donor tagged RasG12V and a GFP2-acceptor tagged RasG12V, due to nanoclustering ^32^. This type of assay can sensitively detect perturbations not only of Ras-nanoclustering, but also of any upstream process, such as correct membrane anchorage or lipid modifications ^32,33^ (**Figure S 1B**).

When palmitoylated, prenylated proteins such as dually palmitoylated H-Ras cannot bind to PDE6D, making them effectively worse intracellular cargo ^8,10^. Hence, loss of PDE6D activity such as by siRNA-mediated knockdown, selectively decreases the BRET-signal of K-RasG12V, but not of H-RasG12V (**Figure S 1B-D**). Using these two BRET-assays, we assessed the intracellular K-RasG12V-membrane anchorage disruption and K-RasG12V-selectivity of compounds. This again revealed an increase in potency amongst the second-round compounds (**Figure 2A**). Compound **4** had the best overall K-RasG12V-selectivity and **15** the best selectivity of top second-round compounds (**Figure 2B**) and both compounds compared favorably in all three BRET-assays relative to the most selective reference compound Deltazinone1 (**Figure 2C-E**).

### Assessing the off-target activity of top compounds

Despite clearly inhibiting PDE6D, several compounds did not display exclusive K-RasG12V-selectivity (**Figure 2B**). This may be due to off-target activities, a problem that was already noted for previous PDE6Di by others ^16,17^.

Broad off-target effects are phenotypically determined by comparing the anti-proliferative effect of compounds on cells with and without the target. We therefore compared the cell growth inhibition of MEF cells with a homozygous CRISPR-mediated knockout (KO) of PDE6D to their wild type (WT) counterpart as a measure of PDE6D-selectivity ^34^ (**Figure S 1E**). In line with the BRET-derived K-RasG12V-selectivity data (**Figure 2B; Figure S 1F**), first-round compounds exhibited a higher PDE6D-selectivity than second-round compounds, with **4** showing again the highest overall selectivity (**Figure 3A**).

**Figure 3.**
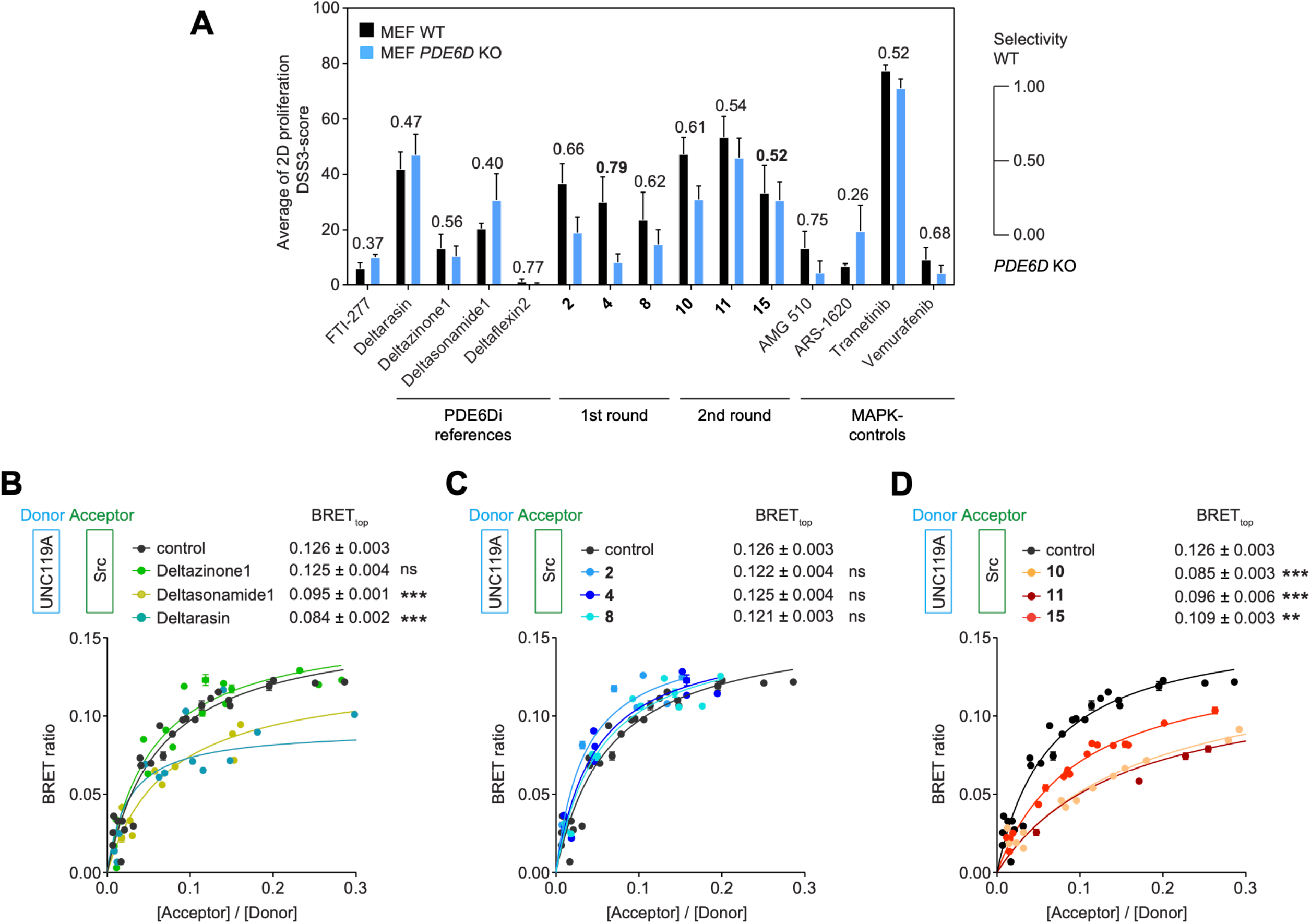
Analysis of PDE6Di off-target activities. (**A**) DSS3-scores of indicated compounds from 2D proliferation assays acquired with WT or *PDE6D*-KO MEFs; n = 4. PDE6D-selectivity was determined as the ratio of the DSS3-scores from WT and the sum of WT and KO MEFs and is indicated above the bars. (**B-D**) BRET-titration curves of UNC119A/ Src complex after treatment with indicated reference PDE6Di (B), top first round (C) or top second round (D) compounds at 5 μM; n ≥ 3. Statistical comparisons of BRETtop values to controls were done using two-tailed Student’s t-test.

UNC119A is a trafficking chaperone of myristoylated proteins and structurally homologous to PDE6D ^12^. Given this relatedness in structure and function, it is a plausible off-target for PDE6Di. We therefore established a BRET-assay to determine the UNC119A-directed off-target activity, by quantifying if the top three compounds from each round disrupted the UNC119A/ Src-complex.

In BRET-titration experiments the characteristic BRET-ratio, BRETtop, that is reached within a defined acceptor-to-donor ratio is a measure for complex stability ^35^. A previously identified inhibitor of UNC119A, Squarunkin A, significantly reduced the BRETtop between UNC119A-Rluc8 and Src-GFP2 (**Figure S 1G**) ^36^. Similarly, treatment with the N-myristoyl-transferase inhibitor IMP-1088 reduced the BRETtop (**Figure S 1G**) ^37^, confirming that our assay can detect myristoyl-pocket dependent disruption of the UNC119A/ Src-interaction.

When testing the reference compounds, we found that surprisingly at 5 μM both Deltarasin and Deltasonamide1, but not Deltazinone1, significantly decreased the UNC119A/ Src-BRET, suggesting off-target binding of the compounds to UNC119A (**Figure 3B**). By contrast, none of our top first-round compounds decreased UNC119A/ Src-BRET (**Figure 3C**), while all our top second-round compounds did, with **15** having the least disruptive activity (**Figure 3D**).

### Inhibition of Ras-signaling and cancer cell proliferation by the top compounds

Next, we continued our selectivity assessment by testing the anti-proliferative activity of the top three compounds from each round on *KRAS*-, *HRAS*- or *BRAF*-mutant cancer cells. In line with in vitro and BRET-data (**Figure 2A**), the anti-proliferative activity was significantly increased in compounds of the second optimization round, with cellular potencies increasing to the low- and sub-micromolar regime (**Figure 4A; Data S2**), but at the expense of selectivity (**Figure 4B**).

**Figure 4:**
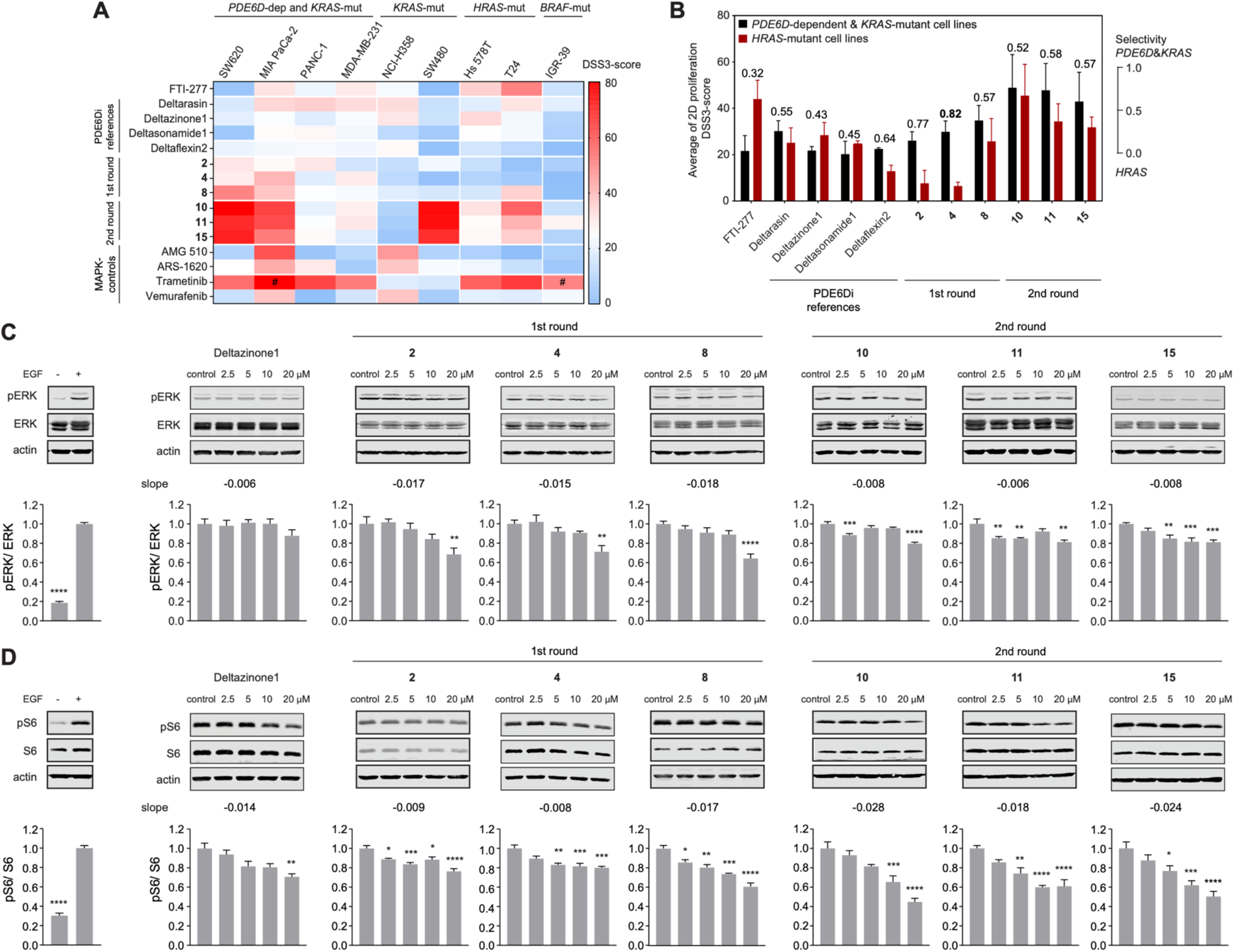
Inhibition of cell proliferation and Ras-signaling by PDE6Di. (**A**) DSS3-scores of indicated compounds from 2D proliferation assays acquired with *PDE6D*-dependent and *KRAS*-mutant, *KRAS*-mutant, *HRAS*-mutant or *BRAF*-mutant cell lines; n ≥ 2; # n = 1. (**B**) Quantification of *PDE6D*-dependent & *KRAS*-mutant-selectivity was performed by determining the ratio of the average of DSS3-scores from *PDE6D*-dependent and *KRAS*-mutant cell lines and the sum of the former and the average DSS3-score of *HRAS*-mutant cell lines from (A); n ≥ 3, except for the condition T24/ compound 8, where n = 2. (**C,D**) Quantified immunoblot data of phosphorylated and total ERK (C; n ≥ 4) or phosphorylated and total S6 (D; n ≥ 3) from *KRAS-G12C* mutated MIA PaCa-2 cells treated with indicated compounds for 4 h before EGF-stimulation; stimulation control data to the far left.

By contrast, **4** displayed the overall highest selectivity for *PDE6D*-dependent and *KRAS*- mutant, as compared to *HRAS*-mutant cancer cell lines (**Figure 4B; Figure S 1H**), consistent with its K-RasG12V-selectivity detected by BRET (**Figure 2B**) and its off-target activity being lowest amongst investigated compounds (**Figure 3**). It therefore surpassed the most selective reference compound, Deltazinone1, ∼6-fold. The highest activity of **4** was seen in MIA PaCa-2 (*KRAS-G12C*-mutant) cells (IC50 = 6 ± 1 μM; **Data S2**), in line with the highest *KRAS*- and *PDE6D*-dependence of this cell line among the tested cell lines (**Figure S 1H**) ^38^.

For compounds that significantly disrupt K-RasG12V-membrane anchorage, it is expected that they also reduce Ras-signaling output. In line with previous data ^7,16^, the reduction in phospho-ERK- (**Figure 4C**) and phospho-S6-levels (**Figure 4D**) downstream of Ras was modest in MIA PaCa-2 cells upon treatment with our top compounds, but better than that seen with the overall best reference compound Deltazinone1.

We subsequently focused our analysis on compound **4**, hereafter named Deltaflexin3, given its overall best performance across all assays and its high water solubility (kinetic solubility, S = 5.68 mM in PBS, pH 7.4, 37 °C).

### PDE6D inhibitor Deltaflexin3 and Sildenafil synergize to inhibit K-Ras activity

The approved drug Sildenafil, which is an inhibitor of cGMP-specific phosphodiesterase type 5 (PDE5), stimulates the PKG2-dependent phosphorylation of Ser181 on the C-terminus of K-Ras ^11^. Given that the phospho-mimetic K-Ras-S181E mutation was shown to reduce the affinity to PDE6D ∼6-fold ^8^, we reasoned that Sildenafil treatment would likewise decrease the affinity.

We therefore sought to increase the anti-tumorigenic activity of Deltaflexin3 by combining it with Sildenafil, which would also focus the inhibitory activity on K-Ras. A more focused inhibition is supported by a survey of >150 small GTPases, which suggests that only 15 other established or predicted PDE6D cargo proteins possess serine or threonine residues in the four residue stretch upstream of the prenylated cysteine that could be affected by Sildenafil in a manner that could impact on PDE6D engagement (**Data S3**).

Using our PDE6D/ K-RasG12V-BRET assay, we found that indeed Sildenafil dose-dependently reduced the BRET-signal consistent with a disruption of the PDE6D/ K-RasG12V-complex (IC50 ∼17 μM) (**Figure 5A**). We then combined Deltaflexin3 with Sildenafil at 10 μM, 20 μM and 30 μM i.e., concentrations that hardly affected the BRET-signal, to test for synergism of these two compounds (**Figure 5A,B**). This analysis revealed a high synergistic activity at ∼20 μM Sildenafil and ∼900 nM Deltaflexin3 (**Figure 5B, right**).

**Figure 5.**
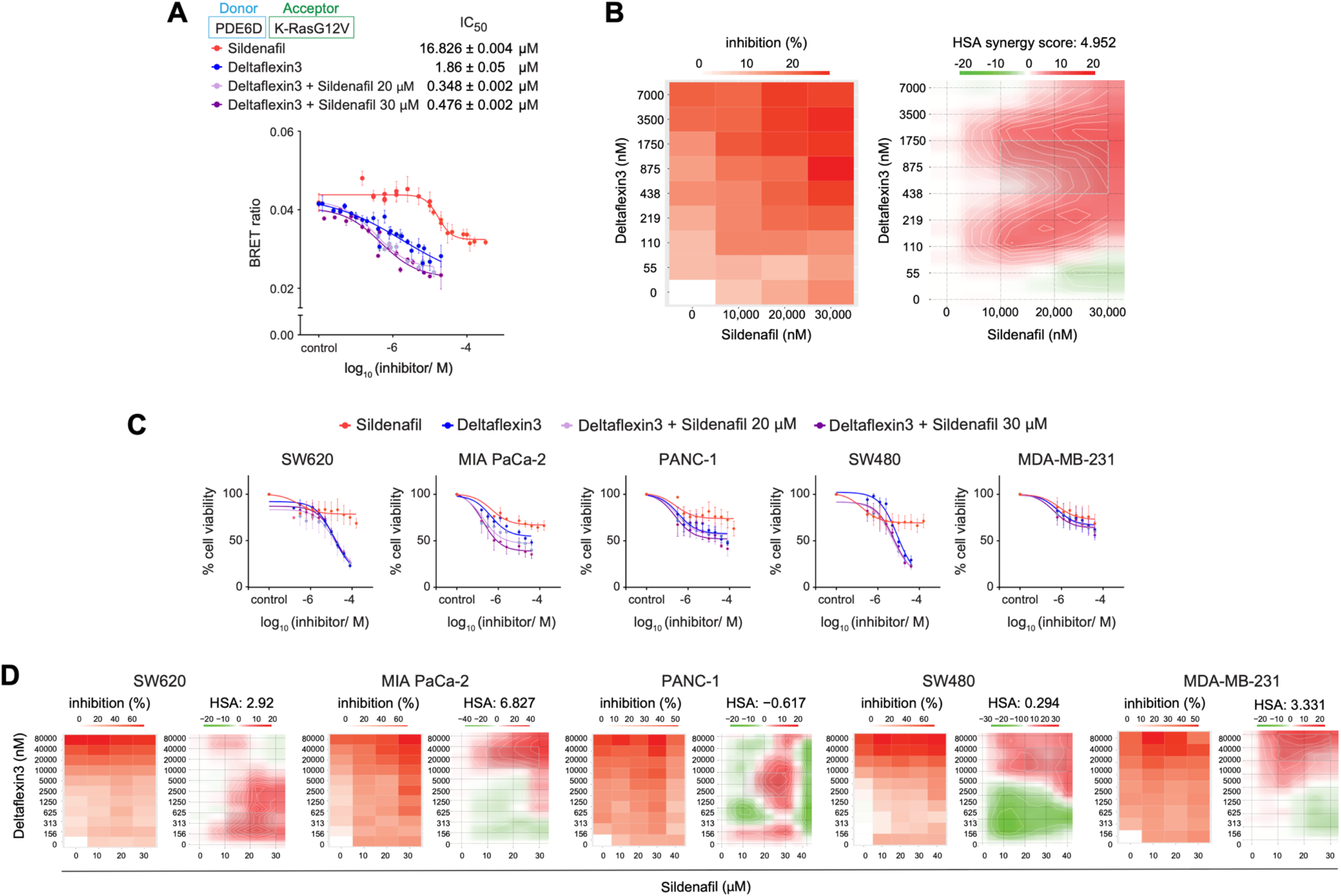
Analysis of Deltaflexin3 (4) and Sildenafil synergism. (**A**) Dose-dependent disruption of PDE6D/ K-Ras complex after treatment with indicated compounds and combinations measured in cellular BRET-assays; n ≥ 3. (**B**) Inhibition (drop in normalized BRET ratio, left) and HSA synergism (right) heatmaps of combinations in (A) and an additional combination with 10 μM Sildenafil; n ≥ 3. Positive HSA synergy scores indicate synergism, while negative scores signify antagonism. (**C**) Compound-dose dependent change of cell proliferation after indicated treatments of *KRAS*-mutant cancer cell lines; n ≥ 2. (**D**) Inhibition and HSA synergism heatmaps for combinatorial Deltaflexin3 and Sildenafil treatment as determined from 2D cell proliferation assays with indicated *KRAS*-mutant cancer cell lines; n ≥ 2.

We therefore continued with a 2D proliferation analysis for synergism in five *KRAS*-mutant and -dependent cancer cell lines with diverse levels of PDE6D- and PKG2-dependencies (**Figure 5C, Figure S 1H**). Amongst the tested cell lines, MIA PaCa-2 showed the highest

HSA synergism score and a clear shift of the inhibition curve to lower concentrations for combinations of the drugs (**Figure 5C,D**). Importantly, high synergism was observed at similar concentrations that were previously identified using the on-target BRET-assay (**Figure 5B,D**).

### Combinations of Deltaflexin3 and Sildenafil efficiently suppress Ras-signaling and microtumor growth

Supported by these proliferation data that suggested a synergism of Deltaflexin3 in combination with Sildenafil, we focused our investigations on MIA PaCa-2 cells.

We first reexamined, whether signaling downstream of Ras was more efficiently inhibited by the combination treatment. Neither Sildenafil at concentrations between 20-30 μM, nor Deltaflexin3 at 10 μM significantly reduced phospho-ERK- (**Figure 6A**) or phospho-S6-levels (**Figure 6B**). Intriguingly, however, the combination of 10 μM Deltaflexin3 and 20 μM Sildenafil significantly reduced phosphorylation levels of both ERK and S6 by ∼ 28 % and ∼ 35 %, respectively.

**Figure 6.**
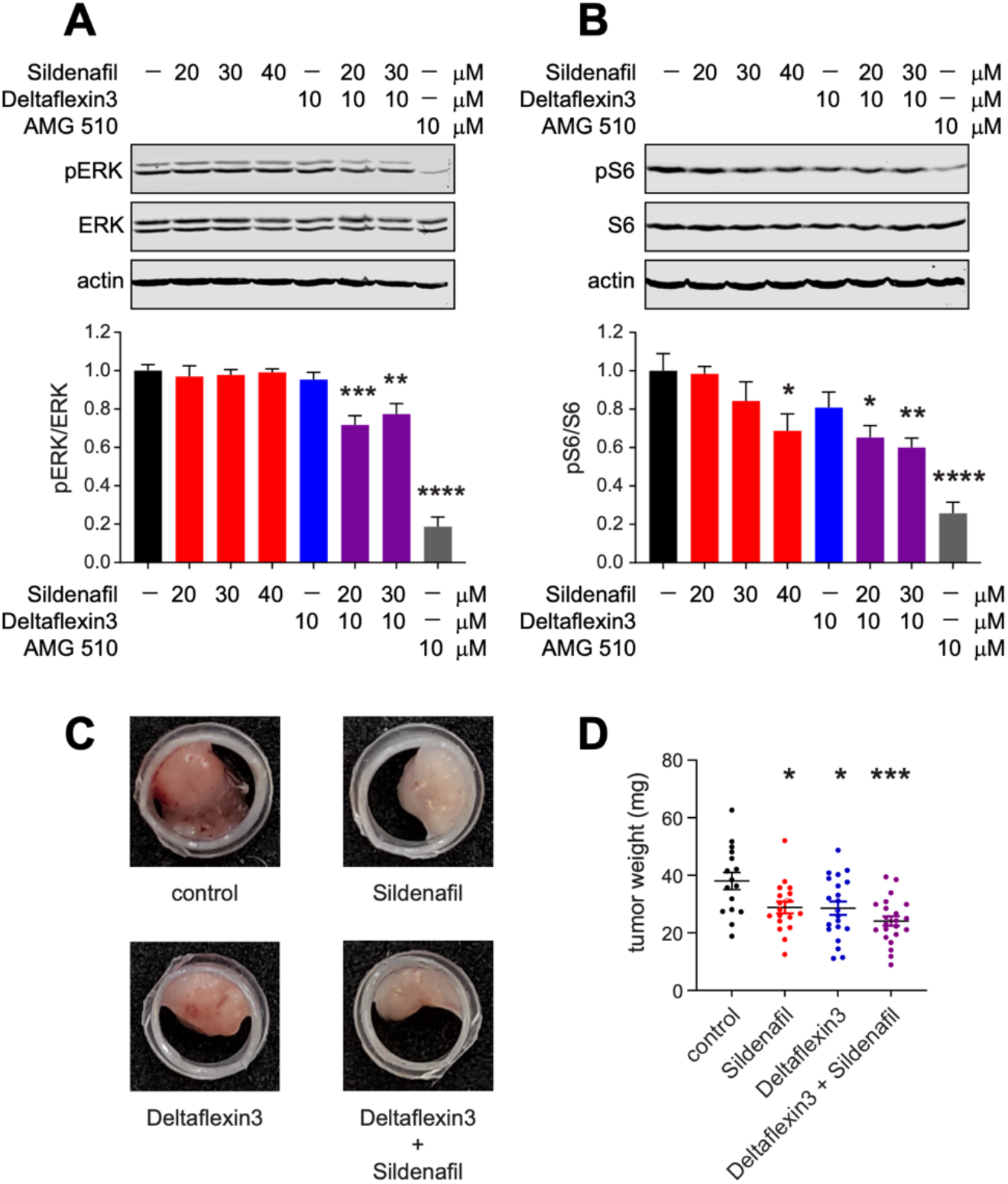
The Deltaflexin3/ Sildenafil combination more potently inhibits Ras-signaling and microtumor growth. (**A,B**) Quantified immunoblot data of phosphorylated and total ERK (A; n ≥ 4) or phosphorylated and total S6 (B; n ≥ 4) from *KRAS-G12C*-mutated MIA PaCa-2 cells treated with indicated compounds for 4 h before EGF-stimulation. (**C**) Representative images of microtumors formed by MIA PaCa-2 cells grown in the CAM assay and treated with inhibitors as indicated. (**D**) Weights of the MIA PaCa-2-derived microtumors (≥ 16 per condition from n = 5) after treatment with 2.5 μM Deltaflexin3 or/ and 30 μM Sildenafil.

Next, we evaluated the anti-tumorigenic activity of Deltaflexin3 in the chorioallantoic membrane (CAM)-assay, where microtumors are raised on the chorioallantoic membrane of fertilized chick eggs ^39,40^. While 10 μM Deltaflexin3 alone significantly reduced MDA-MB-231 cell derived microtumors (**Figure S 1I**), already 2.5 μM Deltaflexin3 were sufficient to achieve a similar reduction in MIA PaCa-2-derived microtumors (**Figure 6C,D**). This is in agreement with the poorer response of MDA-MB-231 to Deltaflexin3 observed in 2D proliferation data (**Figure 4A**). Consistent with the synergistic increase in efficacy observed for the combination of Deltaflexin3 and Sildenafil in BRET-, signaling- and proliferation-assays, MIA PaCa-2-derived microtumor growth was more potently reduced by the combination than by each compound alone (**Figure 6C,D**).

## Discussion

We here developed Deltaflexin3, a nanomolar-active and highly soluble PDE6Di with superior on-target activity as compared to previous reference inhibitors Deltarasin, Deltazinone1 and Deltasonamide1. We show that combinations of Deltaflexin3 with the approved drug Sildenafil synergistically inhibit intracellular binding of K-Ras to PDE6D, and Ras-signaling, proliferation and ex vivo tumor growth of MIA PaCa-2 cells.

Within our dedicated series of 16 compounds, computational docking enabled us to generate several low- and sub-nanomolar binders of PDE6D, which are thus equally potent as previous trailblazer compounds Deltazinone1 and Deltasonamide1. Surprisingly, we measured lower, only submicromolar affinities when employing F-Rheb instead of F-Ator as a probe in our fluorescence polarization-based assay. Interestingly, the submicromolar affinities are more in line with the micromolar activities observed in our BRET- and proliferation-assays (**Data S2**). Previously, we also measured only low micromolar affinities for first generation Deltaflexins and Deltarasin using the F-Rheb probe and in an alternative surface plasmon resonance-based assay that detected the disruption of farnesylated K-Ras binding to PDE6D ^27^. Hence it appears that F-Ator-derived affinities are systematically higher than F-Rheb-derived affinities. The reasons for this are unclear, but it is conceivable that two molecules of F-Ator insert into the hydrophobic pocket of PDE6D, which is large enough to accommodate also dually-geranylgeranylated cargo ^9^. If only one is displaced, the other F-Ator molecule might be able to stabilize the binding of compounds. However, when comparing the F-Rheb derived affinities from our previous compound Deltaflexin2 (Kd[F-Rheb] = 7.17 μM) and Deltaflexin3 (Kd[F-Rheb] = 0.63 μM), a more than 10-fold improvement in affinity becomes apparent.

Another important aspect of our characterization is the dedicated off-target analysis, which has not been done previously. From our BRET-based off-target analysis, it appears that compounds with a PDE6D-affinity below ∼3 nM are more likely to engage UNC119A as an off-target (**Figure 3B-D**). It is plausible that also related UNC119B would be engaged in this way ^41^. Depending on the expression levels of such lipid binding proteins, they may effectively act as sinks for PDE6Di.

In parallel to the UNC119A off-target engagement, water solubility and therefore suitability of compounds for in vivo applications go down. This may not be surprising, as raising compounds with a higher affinity to a highly hydrophobic pocket will render them likewise more hydrophobic. It is possible that this trend then also increases the likelihood of binding to other hydrophobic pockets, such as that of UNC119A.

Importantly, the highest K-RasG12V-selectivity is seen for Deltaflexin3 (**Figure 2B)**, consistent with its lowest off-target effect in both the BRET-based assay looking at UNC119A engagement and its assessment in *PDE6D*-KO MEFs (**Figure 3**). Overall, K-RasG12V-BRET selectivity (**Figure 2B**) and PDE6D-selectivity derived from cell proliferation data of WT and KO-MEFs (**Figure 3A**) show a strong correlation for our compounds, supporting that our assessment selects for least off-target activity (**Figure S 1F**).

PDE6Di development could in the future adopt strategies illustrated in nature. When looking at known cargos of PDE6D, it becomes apparent that their affinity is not modulated within the hydrophobic pocket, but outside of it, at its entry site ^8,9,12^. Contacts with entry site residues are typically not exploited with PDE6Di, albeit our second round of compounds were extended with this goal in mind. Notably for mono-prenylated cargo, it is known that the four residues upstream of the prenylated cysteine significantly modulate the cargo affinity to PDE6D ^8^. While K-Ras has only a moderate micromolar PDE6D-affinity (Kd = 2.3 μM ^8^), the INPP5E-derived peptide has a high, nanomolar affinity (Kd = 3.7 ± 0.2 nM ^12^), and this solely depends on two amino-acids in the four-residue stretch upstream of the farnesylated cysteine ^9^.

The potential of this kind of affinity modulation is essentially illustrated by our Sildenafil data (**Figure 5A**), as Ser181 of K-Ras is part of that four-residue stretch next to the farnesylated cysteine. Therefore, future PDE6Di may rather target that region of the protein, while using a minimal hydrophobic stretch to anchor inside the hydrophobic pocket. We propose that ‘plugging’, rather than ‘stuffing’ the hydrophobic pocket of PDE6D with novel inhibitors may present a way forward.

Inhibitors of Ras membrane anchorage are expected to shut down Ras-signaling output ^5^. For instance, farnesyl-transferase inhibitors that block the enzyme mediating Ras farnesylation are now applied with some success in *HRAS*-mutant head and neck cancers ^42^. While some PDE6Di were shown to dislodge K-Ras more or less from the plasma membrane within 60-90 min ^7,15,16,26^, only in some cases was evidence for a moderate effect on Ras-signaling provided ^16,24,26^. Nevertheless, all of these PDE6Di demonstrated cell killing activity in *KRAS*-mutant pancreatic or colorectal cancer cells, however, these are assays that cannot detect off-target activities.

One explanation for these discrepancies, could be that only a fraction of K-Ras that is trafficked to the plasma membrane does actually depend on PDE6D. We therefore compared the knockdown of PDE6D or that of the alpha-subunit of farnesyl- and geranylgeranyl-transferases with Mevastatin treatment, which would completely block K-Ras membrane anchorage, using our BRET-assay that detects functional K-RasG12V membrane organization (**Figure S 1B**).

These data show that knockdown of the alpha subunit is 49 % as effective as Mevastatin treatment, while PDE6D-knockdown is only 26 % as efficient. This suggest that only between a quarter or a half of functional K-Ras membrane anchorage depends on PDE6D. It is plausible to assume that other trafficking chaperones compensate and salvage K-Ras membrane anchorage thus buffering the loss of PDE6D activity.

It may therefore not be astonishing that both reference PDE6Di Deltazinone1 and our own compounds have such a small effect on phospho-ERK- and phospho-S6-levels (**Figure 4C,D**). Only when combined with Sildenafil could a robust, synergistic ∼28 %-reduction of phospho-ERK- and phospho-S6-levels be observed (**Figure 6A,B**). Indeed, this combination may in general be a way forward for PDE6Di application, as it focuses the inhibitory activity on K-Ras. Apart from K-Ras only 15 other small GTPases can potentially be modulated by both PDE6Di and Sildenafil (**Data S3**).

This synergistic combination also showed promise for the anti-tumorigenic activity of our most selective PDE6Di, Deltaflexin3 (**Figure 6C,D**). However, not all *KRAS*-mutant cancer cell lines respond clearly and synergistically to the Deltaflexin3/ Sildenafil combination (**Figure 5C,D**). MIA PaCa-2 may be particularly responsive, as they have a genetic dependence on both *KRAS* and *PDE6D*, while being not-dependent on *PRKG2* (the gene of PKG2) (**Figure S 1H**). Consequently, this combination could find its application in the treatment of a subset of KRAS-mutant cancers that more often have a high *PDE6D* and a low *PRKG2* expression level, such as colorectal cancer (**Figure S 1J**). However, our analysis of the overall survival of patients with this expression signature across *KRAS*-mutant cancers in the PanCanAtlas dataset shows that they have a significantly better survival than those with the opposite signature (low *PDE6D*/ high *PRKG2*) (**Figure S 1K**). This may indicate a protective effect of the high *PDE6D*/ low *PRKG2* signature, that should not be drug targeted by a PDE6Di/ Sildenafil - combination.

This begs the question as to what specific role PDE6D has for K-Ras trafficking. Given that PDE6D is a major trafficking chaperone of ciliary cargo and that K-Ras has indeed been observed inside the primary cilium ^9^, it is possible that PDE6D inhibition also affects trafficking of K-Ras to this destination. However, the significance of such an inhibition is unclear, given that no function of K-Ras in the cilium is known. Besides, cancer cells are typically not ciliated ^43^, and it would thus not be clear what effect PDE6D inhibition could have in this context.

Another complication of PDE6D as a drug target is its intrinsically broad cargo spectrum ^8,9^. Therefore, its inhibition will not only affect K-Ras and thus *KRAS*-mutant cancer cells, but a host of PDE6D cargos. Finally, the ontogenetic role of PDE6D may be worth considering. Loss of function mutations of PDE6D during development lead to the multisystemic ciliopathy Joubert-Syndrome ^44^. The deletion of PDE6D in mice does not cause gross developmental abnormalities, as mice are fertile and viable ^45^. Some progressive defects in photoreceptor physiology were however observed, as well as an overall reduced body weight. Even though such genetic data do not exactly translate into the effects observed with inhibitors that are typically applied to aged cancer patients, more insight into the PDE6D biology in conjunction with K-Ras seems warranted.

In conclusion, we provide a novel conceptual framework for the future development and application of PDE6Di to be redesigned as ’plugs’ and to be used in combination with PKG2 activators, such as approved Sildenafil. However, we also recommend to better understand the involvement of PDE6D in cancer and the consequences of drug targeting it.

With our novel, potent PDE6D inhibitor Deltaflexin3, which has the highest K-Ras selectivity and lowest off-target activity so far described, we are now providing the currently best tool compound to investigate and further validate the significance of PDE6D (patho)biology.

## Methods

### Cell lines

HEK293-EBNA (HEK) cells were a gift of Prof. Florian M. Wurm, EPFL, Lausanne, Switzerland, and were cultured in Dulbecco’s modified Eagle’s medium (DMEM, #41965-039). WT MEF and MEF PDE6D KO cells (obtained from Prof. Richard A. Kahn, Emory University School of Medicine, Atlanta, USA) were cultured in DMEM. NCI-H358, MDA-MB-231 and IGR-39 were maintained in Roswell Park Memorial Institute medium (RPMI, #52400-025). PANC-1, MIA PaCa-2, Hs 578T and T24 were maintained in DMEM. SW620 and SW480 were maintained in Leibovitz’s L-15 medium (#11415-064). All media were supplemented with 10 % v/v fetal bovine serum (#10270-106), 2 mM L-glutamine (#25030-024) and penicillin 100 U/mL/ streptomycin 100 µg/mL (#15140-122) (complete medium). All cell culture media and reagents were from Gibco, Thermo Fisher Scientific. Cells were grown at 37 °C in a water-saturated, 5 % CO_2_ atmosphere and sub-cultured twice a week. Cell lines SW620 and SW480 were cultured without CO_2_.

### Bacterial strains

Competent *E. coli* DH10B (New England Biolabs, #C3019I), *E. coli* BL21 Star (DE3)pLysS (New England Biolabs, #C2527H) were grown in Luria-Bertani (LB) medium at 37 °C, with appropriate antibiotics unless otherwise mentioned.

### Expression constructs

All expression constructs were produced by multi-site Gateway cloning technology as described ^46^. Briefly, entry clones with compatible LR recombination sites, encoding the CMV promoter, Rluc8 or GFP2 tag and a gene of interest. The location of the tag in the expression constructs is indicated by its position in the construct name, i.e., a tag at the N-terminus of the protein of interest is written before the name of the protein. Genes were obtained either from the Ras-Initiative (K-Ras4BG12V, H-RasG12V both from the RAS mutant clone collection, kit #1000000089 and PDE6D #R702-E30) or by custom synthesis from GeneCust (Src, UNC119A). The cDNAs encoding human c-Src kinase and human UNC119A inserted in the pDONR221 vector were obtained from GeneCust. The three entry clones of promotor, tag and gene of interest were then inserted into pDest-305 or pDest-312 as a destination vector using Gateway LR Clonase II enzyme mix (#11791020, Thermo Fisher Scientific). The reaction mix was transformed into ccdB sensitive *E. coli* strain DH10B (# C3019I, New England Biolabs) and positive clones were selected in the presence of ampicillin. The His6-MBP-Tev-PDEd construct for PDE6D protein production was obtained from the Ras-Initiative (#R702-X31-566).

### In silico docking of compounds

The synthetic rationale for first round compounds was based on computational docking. Three-dimensional coordinates for the molecular structure and sequence of the open and closed conformations of the PDE6D protein (PDB ID: 4JV8 and 1KSH, respectively) were retrieved from the RCSB protein data bank ^7^. The 3D structures of all docked compounds were constructed using Maestro software in the Schrödinger software (Schrödinger Release 2019-2; Maestro, Schrödinger, LLC: New York, NY, USA, 2019). The geometry optimization of docked compounds was performed using the OPLS3 force field ^47^. Powell conjugated gradient algorithm method was applied with a convergence criterion of 0.01 kcal/ (mol Å) and maximum iterations of 1,000.

Molecular docking simulations were performed by using the program Glide ^28^. Flexible compound, extra precision mode and the Epik state penalties were included in the protocol. The MM-GBSA method with VSGB 2.0 solvation model was used to calculate compound binding affinities ^48^. For MM-GBSA calculations, residues within a distance of 8.0 Å from the compound were assigned as flexible.

Computational evaluations to derive second round compounds was slightly different. While using the same protein data as for first round compounds, the putative binding pocket of PDE6D was re-inferred using the software SeeSAR v10.3 (“SeeSAR” 2020) with default parameters and prior domain knowledge to select and refine the most relevant pocket. Compound chemical formulas, defined as SMILES strings, were converted to 3D structures using OpenBabel v2.3.2 with default parameters ^49^. Compounds were docked to PDE6D (PDB ID 4JV8) using SeeSAR v10.3 and the optimal docking pose was manually selected by ranking poses according to their predicted binding affinity and filtering compounds to ensure acceptable lipophilic compound efficiency, limited torsions of the compound backbone and minimal intra- and inter-molecular clashes of the resulting protein-ligand complex.

### Expression and purification of PDE6D

Recombinant PDE6D protein was produced according to a published protocol that was adapted^8^. Briefly, *E. coli* BL21 Star™ (DE3)pLysS strain (#C602003, Thermo Fisher Scientific) was transformed with pDest-His6-MBP-PDE6D and grown at 37 °C in LB medium supplemented with ampicillin at 1:1,000 dilution from 100 mg/ mL stock. When OD reached 0.6, protein expression was induced by adding isopropyl β-D-1-thiogalactopyranoside (IPTG, #437145X, VWR) at 16 °C overnight. Next, the 4 L cultures were pelleted by centrifugation, the pellets were rinsed with PBS and stored at -20 °C until purification.

Purification was conducted using ÄKTA pure chromatography system (Cytiva). All buffers were degassed by placing for 5 min in ultrasonic bath. The cells were lysed by sonication on ice in a buffer composed of 50 mM Tris-HCl, pH 7.5, 150 mM NaCl, 1 mM β-mercaptoethanol, 0.5 mg/ ml lysozyme (#89833, Thermo Fisher Scientific) and protease inhibitor cocktail (#A32955, Pierce). For sonication, a Bioblock Scientific ultrasonic processor instrument (Elmasonic S 40 H, Elma) was used. Lysates were cleared by centrifugation at 18,000 g for 20 min at 4 °C. Cleared supernatant was loaded onto a prepacked HisTrapHP column (#17-5248-02, Cytiva) equilibrated in a binding buffer, which had the same composition as lysis buffer, but without lysozyme and containing 35 mM imidazole. After washing with 20 column volumes, the bound material was eluted by isocratic elution using 100 % of eluting buffer (50 mM Tris-HCl, pH 7.5, 150 mM NaCl, 1 mM β-mercaptoethanol, 500 mM imidazole). The eluted fractions were analyzed by resolving on 4-20 % SDS-PAGE (#4561094 or #4651093 BioRAD) and stained with Roti-Blue quick (#4829-2, Carl ROTH). Fractions were concentrated on AmiconUltra centrifugal filters (molecular weight cut-off, MWCO of 30 kDa, Merck Millipore) by centrifuging at 7,500 g and pulled for dialysis into buffer containing 50 mM Tris-HCl, pH 7.5, 150 mM NaCl, 3 mM DTE, using D-Tube dialyzer with molecular weight cut-off (MWCO) 12-14 kDa, overnight at 4 °C. Next, samples were centrifuged for 15 min at 4,000 g and 4 °C and then loaded onto a size exclusion chromatography column (HiLoad 16/ 600 Superdex 75 pg, with 120 mL column volume, #28989333, Cytiva) at a flow rate of 1 mL/ min, with elution with two column volumes. Fractions were analyzed as above, then concentrated to a volume of about 500 µL. In the next step, protein tags were removed by tobaccoetchvirus (TEV) protease (#T4455, Sigma-Aldrich) (1:25 w/w, TEV/ fusion protein) during overnight dialysis. This step was repeated twice, with 50 % and 70 % approximate cleavage efficiencies. The cleaved mixture was loaded onto HisTrapHP column and the non-bound (tag-free) PDE6D was collected. The collected PDE6D fractions were concentrated using MICROSEP Advance (MWCO 10 kDa, # 88527, Pierce) by centrifugation at 7,500 g and 4 °C. The sample was finally dialyzed overnight in a buffer composed of 20 mM HEPES, pH 7.4, 150 mM NaCl, 5 mM MgCl_2_ and 1 mM TCEP. The PDE6D final concentration of 245.3 µM was determined by Bradford assay. Final purification yield from 4 L starting bacterial culture was 890 µg of PDE6D.

### Fluorescence polarization assay

The IC50 and Kd of compounds to purified PDE6D were determined in a displacement assay using fluorescein-labelled Atorvastatin (F-Ator) or fluorescein-labelled farnesylated Rheb (F-Rheb) peptide as probes ^7,14^. F-Ator was used at 5 nM concentration with 5 nM of PDE6D and F-Rheb peptide was used at 0.5 µM concentration with 2 µM PDED. Assays were carried out in black low volume round bottom 384-well plates (#4514, Corning) with a reaction volume of 20 µL for F-Ator- and 10 µL for F-Rheb-based experiments. Compounds were three-fold diluted in assay buffer (DPBS no Ca^2+^/Mg^2+^; #14190-094, Gibco) with 0.05 % CHAPS (#1479, Carl Roth) for F-Ator based experiments or in a freshly prepared buffer composed of 30 mM Tris, 150 mM NaCl and 3 mM dithiothreitol for F-Rheb based experiments, as described previously ^27,50^. The fluorescence polarization signals were read on the CLARIOstar plate reader (BMG Labtech GmbH) with λ_ex_ = 482 ± 8 nm and λ_em_ = 530 ± 20 nm at 25 °C. The blank corrected milli Polarization value (mP or P × 1,000) calculated from the MARS (BMG Labtech) program was plotted against the logarithmic concentration of inhibitors. The data were fitted into log inhibitor vs. response 4-parametric equation of Prism (GraphPad) to obtain the IC50 values. The IC50 values were converted into Kd using the modified Cheng-Prusoff equation, 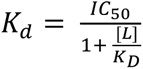, where Kd is the dissociation constant between PDE6D and inhibitor, [L] is the ligand or probe concentration used in the assay and K_D_ is the dissociation constant between the PDE6D and the ligand or fluorescent probe ^27^.The reported K_D_ values were 7.1 ± 4 nM for F-Ator to PDE6D ^7^ and from 0.15 µM ^14^ to 0.45 µM ^12^ for F-Rheb to PDE6D. The mean of the F-Rheb K_D_ value of 0.3 µM was used for the calculations. Note that the concentration of PDE6D is not part of the equation.

### Bioluminescence Resonance Energy Transfer (BRET) assay

BRET assays were essentially performed as described by us previously ^35,51,52^. Briefly, 150,000 to 200,000 HEK293-EBNA cells were plated in 1 mL complete DMEM per well of 12-well cell culture plates (#665180, Greiner bio-one, Merck KGaA). After 24 h, donor and acceptor plasmids were transfected into cells using 3 μL of jetPRIME transfection reagent (#114–75, Polyplus) following the manufacturer’s instructions.

For BRET donor saturation titration experiments, the concentration of donor plasmid (50 ng) was kept constant, and the concentration of acceptor plasmid was increased from 0 to 1,000 ng. The empty pcDNA3.1 plasmid was used to top-up the total DNA load per well to 1,050 ng.

After determination of the optimal acceptor to donor plasmid ratio from titration experiments (A/D plasmid ratio 20:1 for GFP2-K-RasG12V/ Rluc8-PDE6D, 5:1 for GFP2-K-RasG12V/ Rluc8-K-RasG12V, 3:1 for GFP2-HRasG12V/ Rluc8-HRasG12V and 20:1 for UNC119A-Rluc8/ Src-GFP2), compound dose-response experiments were performed. 24 h after transfection, cells were treated for another 24 h with DMSO 0.1 % v/v as vehicle control or with compounds at 5 to 8 different concentrations ranging from 20 µM to 0.15 µM, prepared as 2-fold dilution series in complete medium.

To study the effect of siRNA-mediated knockdown, cells were plated and after 24 h co-transfected with 50 nM siRNA and 500 ng plasmid DNA per well (same A/D plasmid ratio as described above) using 4 μl Lipofectamine 2000 (#11668019, Thermo Fisher Scientific) in Opti-MEM medium (#31985062, Gibco).

BRET-measurements were performed on a CLARIOstar plate reader at 25 °C after 48 h as described ^35,51,52^. Technical quadruplicates were measured using specific channels for the luminophores (GFP2-acceptor signal, RFU, at λ_ex_ = 405 ± 10 nm and at λ_em_ = 515 ± 10 nm; after 10 μM coelenterazine 400a (#C-320, Gold Biotechnology) addition, simultaneous recording of Rluc8-signals for donor signal, RLU, λ_em_ = 410 ± 40 nm and for the BRET-signal at λ = 515 ± 15 nm). The BRET ratio was calculated as before ^35,51,52^.

For BRET donor saturation titration experiments, the BRET ratio was plotted against the relative expression. The relative expression of acceptor to donor ([Acceptor]/[Donor]) was determined as the ratio between RFU and RLU. All independent repeat experiments were plotted at once using these normalized data i.e., BRET ratio against relative expression. The data were fitted into one phase association equation of Prism 9 (GraphPad) and the top asymptote Ymax-value was taken as the BRETtop. It represents the maximal BRET ratio reached within the defined [Acceptor]/[Donor] ratio. Statistical analysis between the BRETtop values was performed using the student’s t-test.

### 2D cell proliferation assay

Cancer cells were seeded at a density of 1,000 cells/ 100 μL complete medium into 96-well cell culture plates (#655180, Greiner bio-one, Merck KGaA). After 24 h, control and test compounds were added to the cells with DMSO (0.1 % v/v) as a vehicle control. Compound activities were analyzed from 9-point dose-response curves, with compounds prepared as 2-fold dilution series ranging from 40 µM to 0.15 µM (PDE6Di and FTI-277) or from 20 µM to 0.02 µM for MAPK-control compounds. Following incubation for 72 h with the compounds, the cell viability was assessed using the alamarBlue reagent (#DAL1100, Thermo Fisher Scientific) according to the manufacturer’s instructions. After addition of alamarBlue reagent at a 10 % v/v final volume, cells were incubated for 2 to 4 h at 37 °C. Then, the fluorescence intensity was read at λ_ex_ = 530 ± 10 nm and λ_em_ = 590 ± 10 nm at 25 °C using a CLARIOstar plate reader. The obtained raw fluorescence intensity data were normalized to vehicle control (100 % viability) and plotted against the compound concentration.

### Drug sensitivity score analysis (DSS3)

As described before ^51^, a drug sensitivity score (DSS) analysis was performed in order to quantify the drug sensitivity with a more robust parameter than the IC50 or EC50 values. DSS values are normalized area under the curve (AUC) measures of dose-response inhibition data, where the DSS3-score takes drug-responses better into account that are achieved across a broad concentration range ^29^. Drug response data from BRET assays or 2D cell proliferation assays were prepared according to the example file on the Breeze website (https://breeze.fimm.fi/), uploaded and analyzed ^53^. The output file included DSS3 scores as well as several other drug sensitivity measures such as IC50 and AUC.

### Synergy analysis of drug combinations

The synergistic potential of compounds was analyzed essentially as described before ^52^. For PDE6D/ K-RasG12V BRET-experiments, full dose-response analyses of Deltaflexin3 (between 7 µM to 0.014 µM) or Sildenafil (between 320 µM to 1.8 µM) alone or for Deltaflexin3 in combination with Sildenafil maintained at a fixed concentration of either 10, 20, 30 μM were performed. For 2D proliferation experiments, full dose-response analyses of Deltaflexin3 (between 80 µM to 0.156 µM) or Sildenafil (between 160 µM to 0.312 µM) alone or for Deltaflexin3 in combination with Sildenafil maintained at a fixed concentration of either 10, 20, 30 or for some 40 μM were performed. Comparison between the drug response profiles of the combinations and the profiles of each single agent was then carried out using the web-application SynergyFinder ^54^(https://synergyfinder.fimm.fi). We employed the HSA model, which considers that the expected drug combination effect corresponds to the maximum of the single agent responses at the corresponding concentrations. The resulting HSA synergy score S_HSA_ is defined as follows

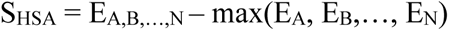

with E_A,B,…,N_ being the combination effect between N drugs and E_A_, E_B_,…, E_N_ being the single agent responses at the corresponding concentrations.

### ATARiS gene dependence score

Gene dependence scores of selected genes of interest for cancer cell lines used in this study were obtained from the drive data portal (https://oncologynibr.shinyapps.io/drive/). The DRIVE project has provided the dependence data of 7,837 genes for 398 cancer cell lines, as determined by large-scale RNAi screening in cell viability assays ^38^. A double gradient heatmap for the extracted gene dependence scores was then generated using GraphPad Prism software.

### Immunoblotting

Following a 16 h serum starvation, MIA PaCa-2 cells were treated with 0.1 % v/v DMSO vehicle control or with compounds at 37 °C for 4 h and then stimulated with 200 ng/mL human epidermal growth factor (hEGF, #E9644, Sigma) at 37 °C for 10 min. *In situ* cell lysis was performed in ice-cold lysis buffer (50 mM Tris-HCl pH 7.5, 150 mM NaCl, 0.1 % v/v SDS, 5 mM EDTA, 1 % v/v Nonidet P-40, 1 % v/v Triton X-100, 1 % v/v sodium-deoxycholate, 1 mM Na_3_VO_4_, 10 mM NaF, 100 µM leupeptin and 100 µM E64D protease inhibitor) supplemented with a cocktail of protease inhibitors (#A32955, Pierce) and a cocktail of phosphatase inhibitors (PhosSTOP, #4906845001, Roche Diagnostics GmbH). After lysate clarification, the total protein concentration was determined by Bradford assay using the Quick Start Bradford 1x Dye reagent (#5000205, Bio-Rad) and BSA (#23209, Thermo Fisher Scientific) as a standard. Proteins (50 µg per lane) were resolved by SDS-PAGE in a 10 % v/v homemade polyacrylamide gel under reducing conditions and transferred to a nitrocellulose membrane by semi-dry transfer (kit #1704272, Bio-Rad). Membranes were saturated in phosphate-buffered saline (PBS) containing 2 % w/v bovine serum albumin (#A6588, AppliChem GmbH) and 0.2 % Tween for 1 h at room temperature, then incubated with primary antibodies overnight at 4 °C. For phospho-ERK and phospho-S6 detection, a combination of mouse anti-phospho-ERK and rabbit anti-ERK or a combination of rabbit anti-phospho-S6 and mouse anti-S6 antibodies were used, respectively (see Key Resources). Incubation with secondary antibodies was performed for 1 h at room temperature. Each antibody incubation was followed by at least three wash steps in PBS supplemented with 0.2 % v/v Tween 20. Signal intensities were quantified using the Odyssey Infrared Image System (LI-COR Biosciences). The ratio between the intensities obtained for phosphorylated protein versus total protein was calculated and then normalized to the sum of all the ratios calculated for one blot to make blots comparable by accounting for technical day-to-day variability. For representative purposes, data were scaled to the controls present on each blot and are represented as the mean ± SEM of at least three independent biological repeats. The slope of the dose-response data was determined from fitting a line using GraphPad Prism. For each blot, either β-actin or GAPDH levels were determined as a loading control.

### Chorioallantoic membrane (CAM) assay

Fertilized chicken eggs were obtained from VALO BioMedia GmbH (Osterholz-Scharmbeck, Germany) and, on day 1, the development of the embryos was started by incubating the eggs at 37 °C in a > 60 % humidified egg hatcher incubator (MG200/300, Fiem). A small hole was made with the help of an 18 Gauge needle (#305196, Becton Dickinson) into the narrower end of each egg on day 3 and was kept covered with parafilm to avoid contamination. On day 8, 2 × 10^6^ MDA-MB-231 cells, or 3.5 × 10^6^ MIA PaCa-2 cells were resuspended in 10 µL cell culture medium without FBS and mixed 1:1 with Matrigel (#356234, Corning). This mix was then deposited in sterilized 5 mm diameter plastic rings cut from PCR tubes (#683201, Greiner bio-one, Merck KGaA) on the surface of a chicken embryo chorioallantoic membrane. After 1 day, the growing tumors were treated with a volume identical to the deposited cell suspension of 0.2 % v/v vehicle control or test compounds 2× concentrated in medium without FBS ^31,39^. Treatment was performed daily and after 5 days of treatment the microtumors were harvested at day 14. Then the tumor weight was determined using a balance (E12140, Ohaus).

### Survival analysis

All data were retrieved from TCGA Pan-Cancer Atlas (https://dev.xenabrowser.net/datapages/?cohort=TCGA%20Pan-Cancer%20) (PANCAN). The 647 cancer samples with non-silent KRAS mutation were selected. We used the list of non-silent somatic mutations as defined in Xena (https://ucsc-xena.gitbook.io/project/overview-of-features/visual-spreadsheet/mutation-columns).

Expression data was retrieved for *PDE6D* and *PRKG2* genes data in “batch effects normalized mRNA data” units, and samples were split in 4 groups according to high or low expression of each gene, setting the limit at median expression value for each gene. The difference between the two curves was tested using Kaplan Meyer estimation. Data analyses were performed in R version 4.2.1 ^55^. Survival analyses and plots were done using survival v.3.4 ^56^ and survminer v 0.4 ^57^ libraries.

### Quantification and Statistical Analysis

For statistical analysis and plot preparation, GraphPad Prism (version 9.5.1 for Windows, GraphPad Software, USA, www.graphpad.com) was used. The sample size n represents the number of independent biological repeats and is indicated in the respective figure legends. All graphs show mean values ± SEM across all technical and biological repeats. We determined statistical differences to control samples by employing one-way ANOVA with Tukey’s multiple comparison test, unless otherwise mentioned in the legends. A p value of < 0.05 is considered statistically significant. Statistical significance levels are annotated in the plots as * = p < 0.05; ** = p < 0.01; *** = p < 0.001; **** = p < 0.0001.

## Data availability

This study did not report standardized datatypes. All unique/ stable reagents generated in this study are available from the corresponding author with a completed materials transfer agreement.

## Supporting information

DataS1_CompoundSynthesis

DataS2

DataS3

## Acknowledgements

We thank Professor Richard A. Kahn (Emory University School of Medicine, Atlanta, USA) for providing the PDE6D KO cell line. We are grateful to Dr. Eyad K. Fansa (Max Planck Institut, Dortmund, Germany) for giving us FITC-labelled farnesylated Rheb peptide.

DKA received a Grant4Targets grant (Ref. 2019-08-2426) from Bayer AG.

This work was supported by grants from the Luxembourg National Research Fund (FNR): AFR individual grant 13589879 to PK and PoC20/15269106-inhibitPDE-RASv2 to DKA.

## Author Contributions

PK characterized compounds by BRET, in proliferation experiments, extracted ATARiS information, performed synergy experiments and analyzed these data.

ESR and MB performed immunoblot experiments and analyses and ESR carried out the CAM assay and analyzed WB and CAM assay results.

GM collected FP data and evaluated them. AG did gene expression and survival analyses.

VV and AK generated the in silico library and performed computational docking experiments of first round compounds.

ML and EG performed computational docking experiments of second round compounds. PK and ESR helped to prepare the manuscript.

DKA initiated the study, supervised the project, designed compounds, and wrote the manuscript.

## Supplemental Information

**Supplementary Figure S1:** Data supplementing information in the main figures.

**Data S1: Compound Synthesis.** Chemical synthesis routes and compound analytics.

**Data S2: Activity Data Summary.** Collects data from plots by figure and shows in the first tab a table collecting all activity data per compound.

**Data S3: Survey of potential PDE6D cargo amongst all small GTPases.**

Based on the four residues upstream of the prenylated cysteine, we identified those small GTPases that are putative PDE6D cargo and contain serine or threonine residues in that stretch that could be targeted by Sildenafil-stimulated PKG2 phosphorylation.

**Table S1:**
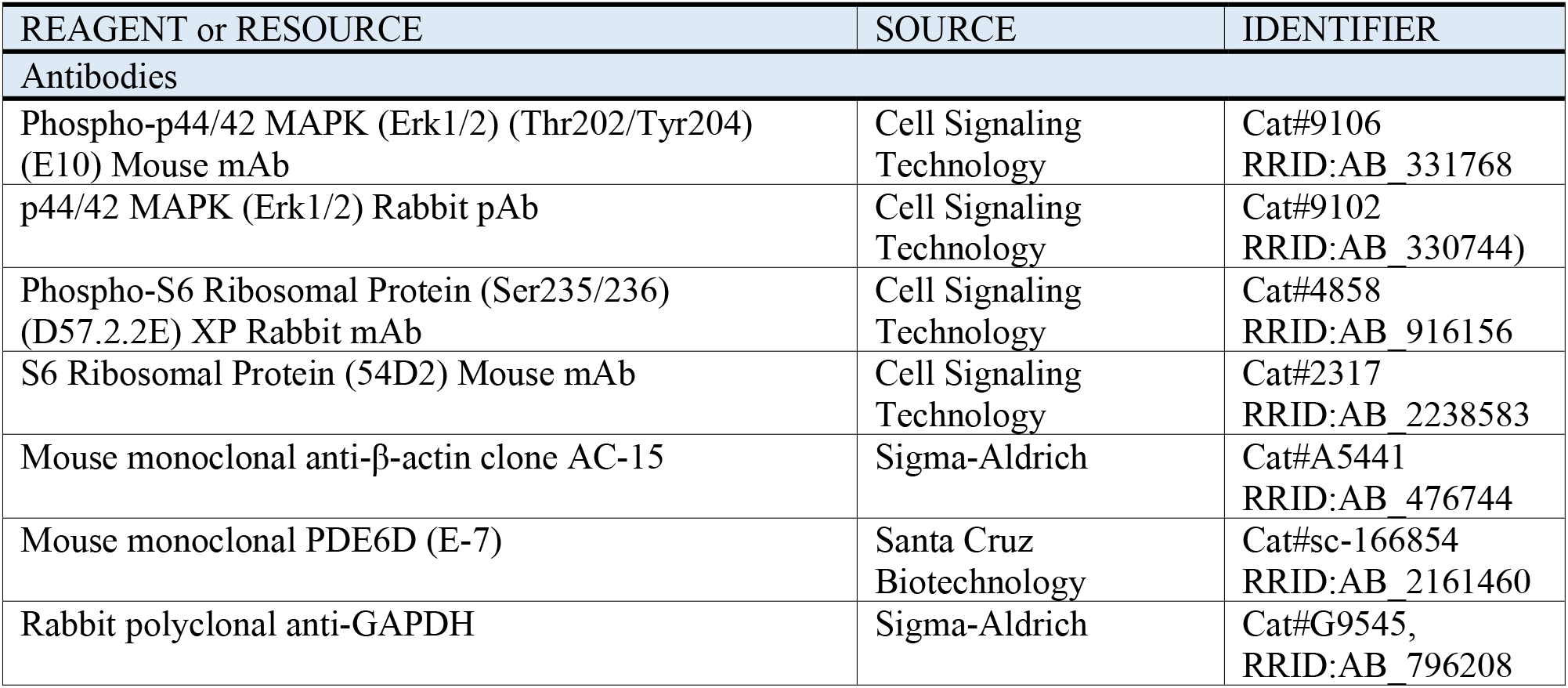
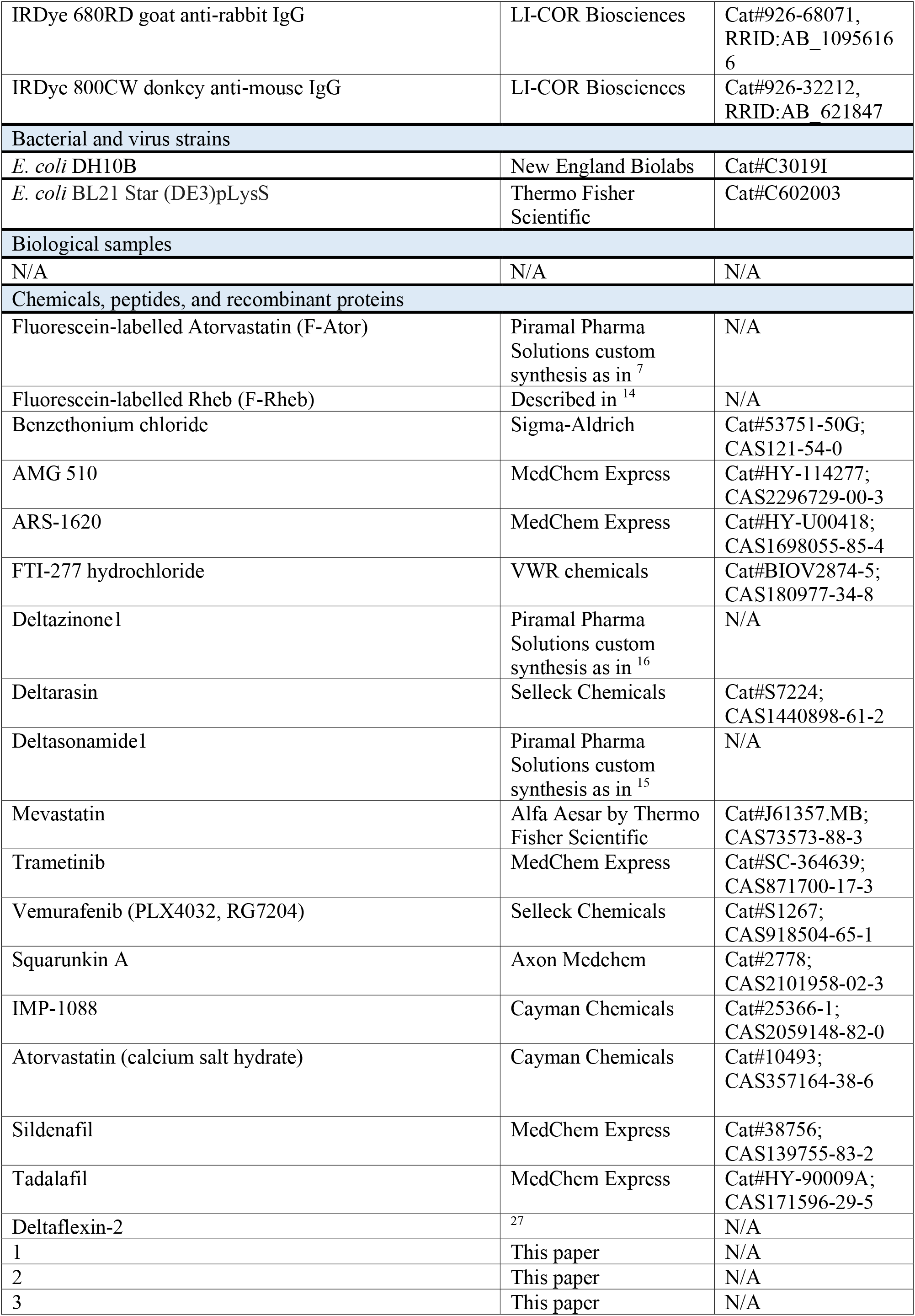
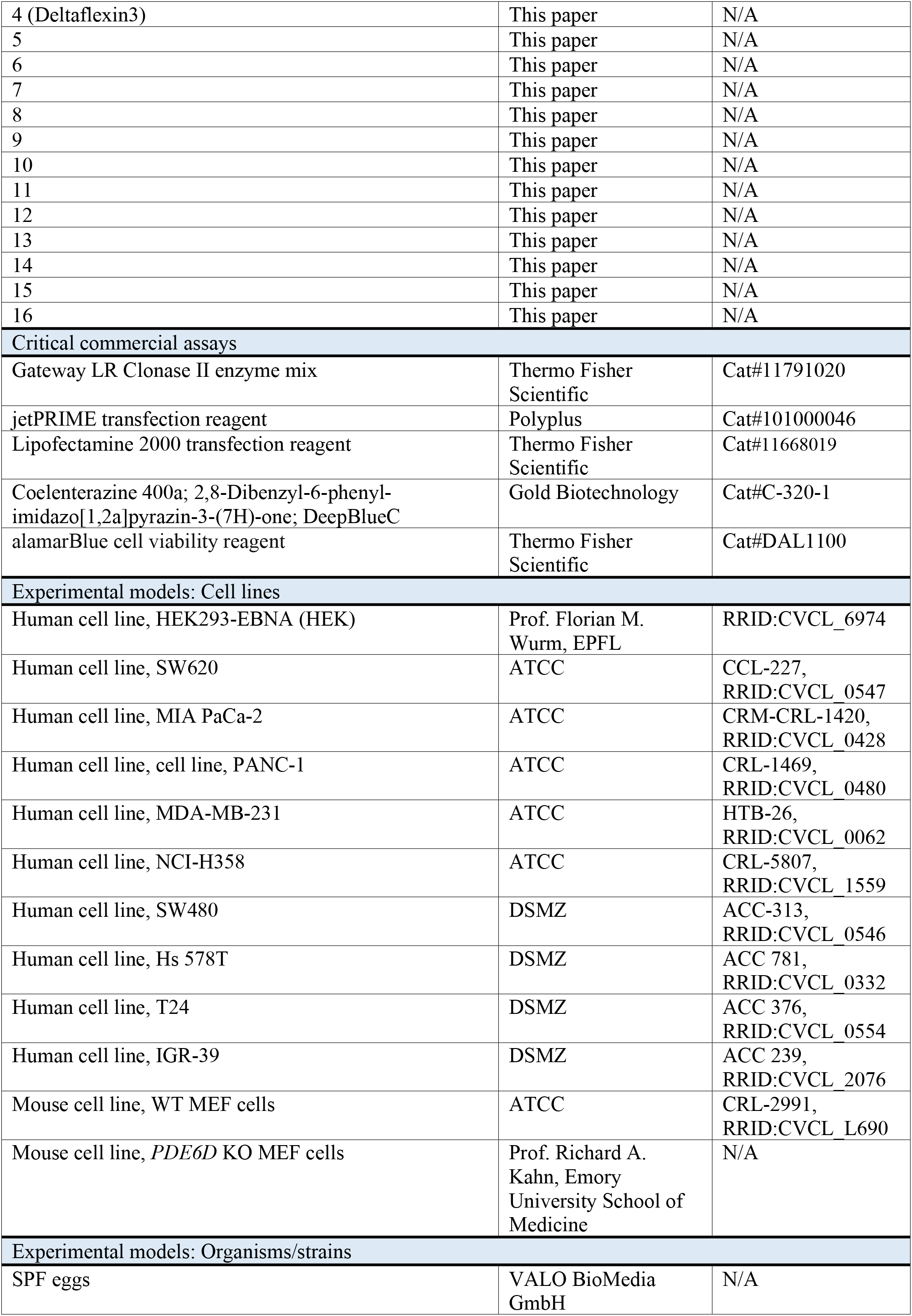
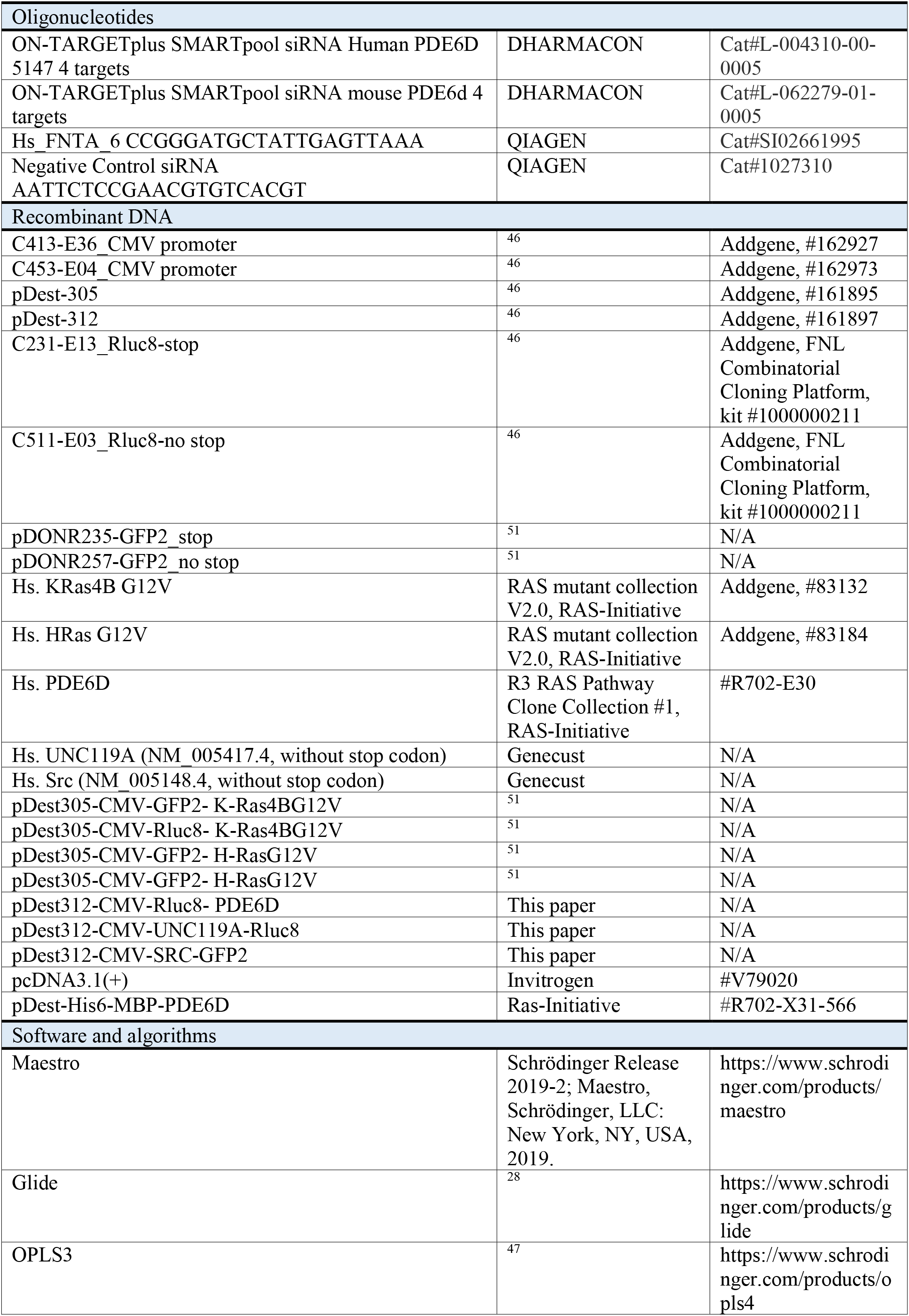
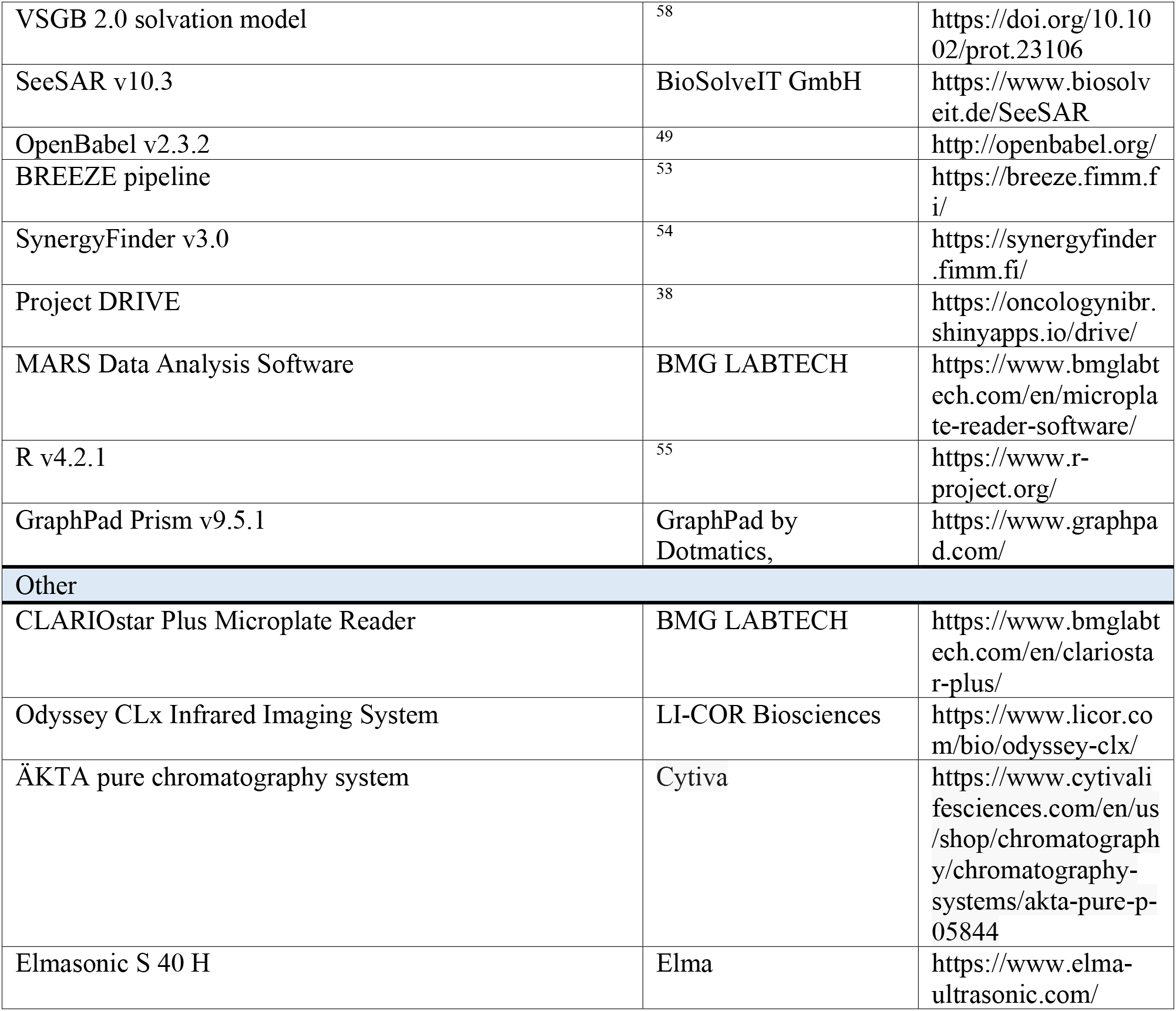
Materials employed in the study.

