## Supplementary material for "An improved PDE6D inhibitor combines with Sildenafil to synergistically inhibit KRAS mutant cancer cell growth": DataS1_CompoundSynthesis

### Supplementary Data S1 related to Compound synthesis and analytical data in the Methods section - chemical compound synthesis, of:

### An improved PDE6D inhibitor combines with Sildenafil to synergistically inhibit KRAS mutant cancer cell growth

#### Syntheses

##### General Information

Altogether 16 different heptylbenzamides and heptanamides were synthesized (Scheme S1) as described below.

Intermediates in schemes are labelled with S as prefix and consecutively numbered within each scheme or shared between consecutive schemes describing related syntheses. Those intermediates used in several syntheses across schemes have in addition the prefix 'Int.-' ; corresponding designations are unique.

Unless otherwise noted, reactions were performed under an argon atmosphere in glassware that was flame-dried and equipped with a magnetic stirring bar. Plastic syringes were used to transfer air- and moisture-sensitive reagents. Solvent was freshly distilled/ degassed prior to use unless otherwise noted. Analytical TLC was performed with silica gel GF254 plates. All compounds were judged pure by TLC analysis (single spot/ two solvent systems) using a UV lamp or spray reagents for detection purposes. For column chromatography, a 200-300 mesh silica gel was employed. Organic solutions were concentrated under reduced pressure using a rotary evaporator. Room temperature (RT) is 23 - 25°C. The reaction temperatures refer to internal reaction temperatures.

Deuterated solvents were purchased from Cambridge Isotope Laboratories. $^{1}$H-NMR spectra were recorded on Bruker AVANCE III 400, Chemical shifts (δ) were reported in ppm relative to the residual solvent signal (DMSO-$d_6$ δ = 2.50 ppm for $^{1}$H-NMR). Chemical shifts (ppm) were recorded with tetramethylsilane (TMS) as the internal reference standard. Multiplicities are given as s (singlet), d (doublet), t (triplet), dd (doublet of doublets), td (triplet of doublets) or m (multiplet). The coupling constants (J) are given in Hertz (Hz). Mass spectra were obtained using electro spray ionization (ESI-TOF). LCMS mass spectra were acquired at high resolution on a Single quadrupole mass spectrometer typically coupled to a Waters Acquity UPLC H-Class with PDA.

*LCMS-Method-C3:* Acquity BEH C18; particle size: 1.7 µm; column size: 2.1 x 50 mm; Eluent A: 2mM ammonium acetate followed by 0.1%formic acid in water; Eluent B: 0.1% formic acid in acetonitrile; Gradient: 98:2 at 0.01 min to 0.30 min at Flow rate: 0.550ml/min, 50:50 at 0.60 min at Flow rate: 0.550ml/min, 25:75 at 1.10 min at Flow rate: 0.550 ml/min, 00:100 at 2.0 min up to 2.70 min at Flow rate: 0.600 ml/min, 98:2 at 2.71 min up to 3.00 min at Flow rate: 0.550 ml/min . Column temperature: Ambient.

*LCMS-Method-J2:* Acquity BEH C18; particle size: 1.7 µm; column size: 2.1 x 50 mm; Eluent A: 2mM ammonium acetate followed by 0.1%formic acid in water; Eluent B: 0.1% formic acid in acetonitrile; Gradient: 98:02 at 0.01 min up to 0.50 min at Flow rate: 0.450ml/min, 30:70 at 3.00 min, 05:95 at 4.0 min up to 5.50 min at Flow rate: 0.500ml/min, 98:02 at 5.51 min up to 6.00 min at Flow rate: 0.450 ml/min; Column temperature: Ambient.

*PDS_HPLC_METHOD-35*: YMC Triart C18; particle size: 5 µm; column size: 4.6 x 150 mm; Eluent A: 0.1%formic acid in water; Eluent B: 0.1% formic acid in acetonitrile; Gradient: 90:10 at 0.01 min, 10:90 at 5.00 min, 5:95 at 6.00 min up to 10.00 min, 90:10 at 10.01 min up to 14.00 min at Flow rate: 1.0 ml/min; Column temperature: 25°C

*PDS_HPLC_METHOD-59*: X-Bridge C18; particle size: 3.5 µm; column size: 4.6 x 100 mm; Eluent A: 0.1%formic acid in water; Eluent B: 0.1% formic acid in acetonitrile; Gradient: 90:10 at 0.01 min, 10:90 at 4.00 min, 5:95 at 5.00 min up to 7.50 min, 90:10 at 7.51 min up to 10.00 min at Flow rate: 1.0 ml/min; Column temperature: 25°C

Commercial reagents were purchased from Sigma-Aldrich, TCI, Combi-Blocks, Alfa, Spectrochem or GLR, and used as received. Other commercially available reagents and solvents were used without further purification.

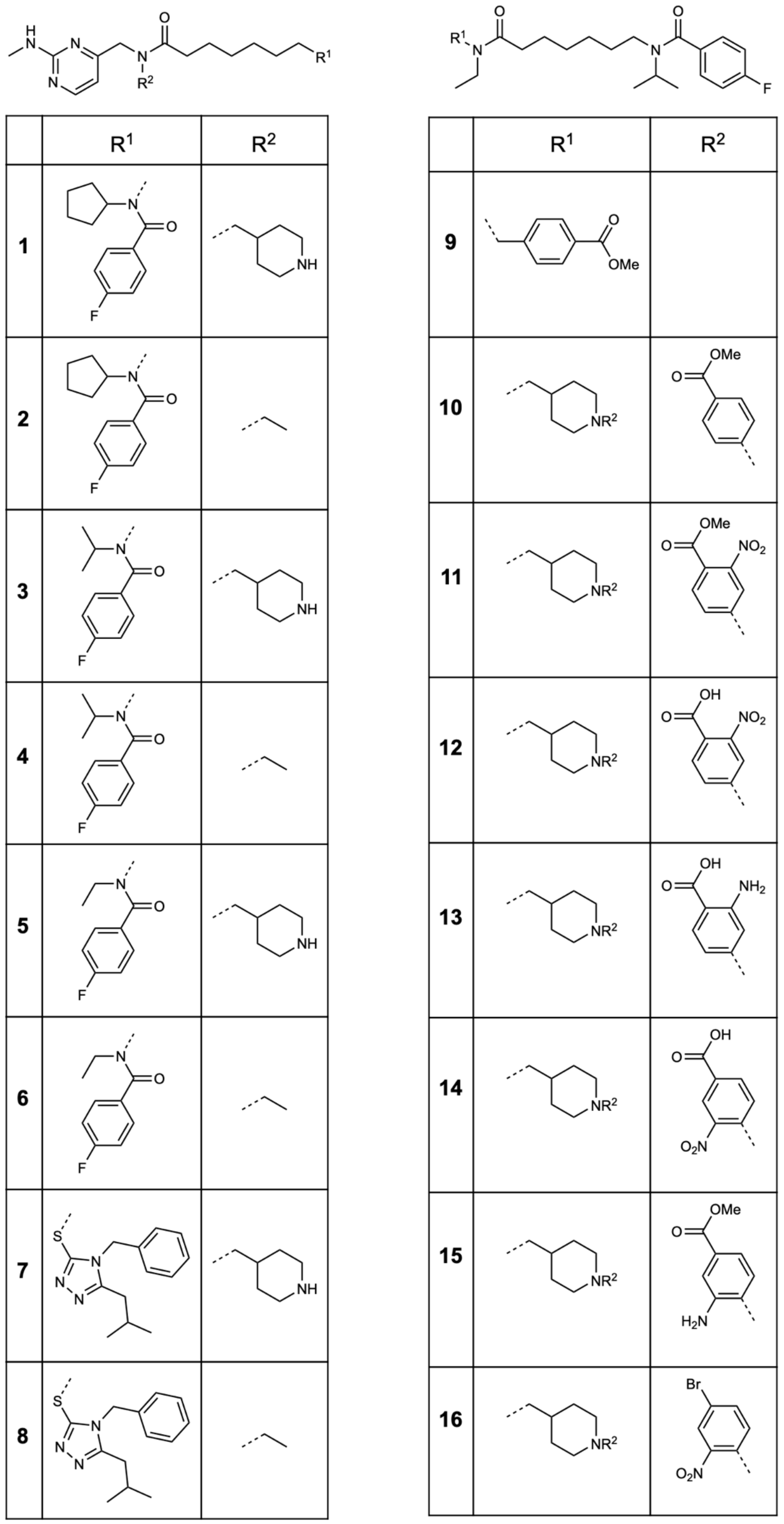

**Scheme S1: Compounds synthesized in this study**

#### Synthesis of (1) and (2) described in Schemes S2 and S3

**Synthesis of N-cyclopentyl-4-fluoro-N-(7-(((2-(methylamino)pyrimidin-4-yl)methyl)(piperidin-4-ylmethyl)amino)-7-oxoheptyl)benzamide (1)**

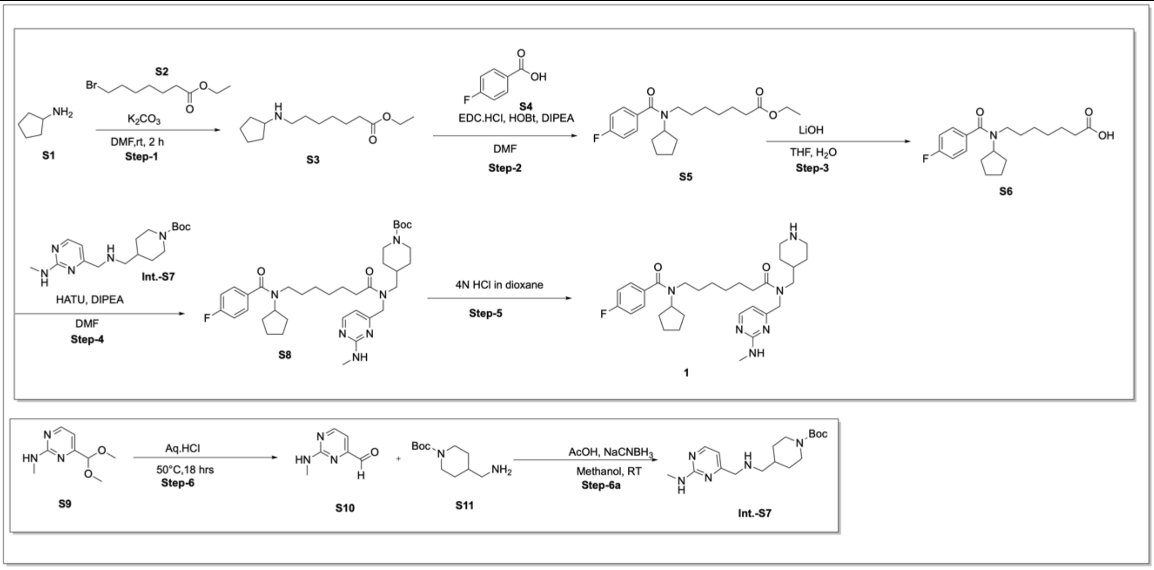

**Scheme S2**

**Step-1: Synthesis of compound (Ethyl 7-(cyclopentylamino)heptanoate) (S3)**

To a solution of cyclopentanamine (**S1**) (1.0 g, 11.7 mmol, 1.0 eq) in DMF (10 ml) was added $K_2CO_3$ (3.24g, 22.5 mmol,2.0 eq) and resulting mixture was stirred at RT 1 h. After 1 h Ethyl 7-bromoheptanoate (**S2**) (3.31g, 14.10 mmol, 1.2 eq) was added dropwise at RT and resulting mixture was stirred at RT for 3 h. After completion of reaction, the reaction mixture was poured into water, extracted with ethyl acetate (3×100ml). The combined organic layer was dried over $Na_2SO_4$ and concentrated under vacuum. The crude product was purified by column chromatography on silica gel (60-120 mesh size) using Dichloromethane:Methanol (2-5%) as eluent. The desired product isolated as solid (1.5 g, 53 %). LCMS: 100%, ESI-MS 242.2 m/z [M+H]+. $^1$H NMR (400 MHz, Chloroform-d) δ 4.16-4.10(q, 2H), 3.49 (s, 1H), 3.31-3.24(m, 1H), 2.81-2.77(t, 2H), 2.31-2.27 (t, 2H), 2.03-2.00 (m, 2H), 1.83-1.70 (m, 6H), 1.65-1.57 (m ,4H), 1.37-1.36(m ,4H), 1.30-1.24 (m, 3H).

**Step-2: Synthesis of compound (Ethyl 7-(N- cyclopentylfluorobenzamido)heptanoate) (S5)**

To a solution of Ethyl 7-(cyclopentylamino)heptanoate (**S3**) (0.750 g, 4.6 mmol, 1.0 eq) in DMF (10 ml) was added EDC.HCl (2.0 g,10.7 mmol, 2.0 eq), DIPEA (2.7 ml, 16.0 mmol, 3.0 eq), HOBT(0.144 g, 1.07 mmol,0.2 eq) and 4-Fluorobenzoic acid (**S4**) (1.5 g, 6.42 mmol, 1.2 eq) at RT. The reaction mixture was stirred at RT for 16 h. After completion of reaction, the reaction mixture was poured into water, extracted with ethyl acetate (3×100 ml). The combined organic layer was dried over $Na_2SO_4$ and concentrated under vacuum. The crude product was purified by column chromatography on silica gel (60-120 mesh size) using Dichloromethane:Methanol (3-5%) as eluent. The desired product isolated as solid (1.4 g, 62%). LCMS: 100%, ESI-MS 364.36 m/z [M+H]+. $^1$H NMR (400 MHz, DMSO-d6) δ 7.41-7.68(t, 2H), 7.28-7.24 (t, 2H), 4.08-4.03(m, 2H), 3.19(s, 2H), 2.27 (s, 2H), 1.64-1.41 (m, 12H), 1.25-1.17 (m, 8H).

**Step-3: Synthesis of compound (7-(N-cyclopentyl-4-fluorobenzamido)heptanoic acid) (S6)**

To a solution of Ethyl 7-(N-cyclopentyl-4-fluorobenzamido)heptanoate (**S5**) (1.4 g, 3.8 mmol, 1.0 eq) in THF:$H_2O$ (1:1) (14 ml), LiOH (0.320 g,7.7 mmol, 2.0 eq) was added at RT. The mixture was stirred at RT for 3 h. After completion of the reaction, the reaction mixture was diluted in water (50 ml), extracted with ethyl acetate (1×50 ml). The aqueous layer was acidified with 2N HCl until pH~2, It was extracted with ethyl acetate (3×100 ml). The combined organic layer was dried over $Na_2SO_4$ and concentrated under vacuum. The crude product was directly used in the next step without further purification (1.0 g, 90%). LCMS: 100%, ESI-MS 336.34 m/z [M+H]+.

**Step-4: Synthesis of compound Tert-butyl 4-((7-(N-cyclopentyl-4-fluorobenzamido)-N-((2-(methylamino)pyrimidin-4-yl)methyl)heptanamido)methyl)piperidine-1-carboxylate (S8)**

To a solution of 7-(N-cyclopentyl-4-fluorobenzamido)heptanoic acid (**S6**) (0.10 g, 0.28 mmol, 1.0 eq) in DMF (1 ml) was added HATU (0.168 g, 0.4 mmol, 1.5 eq), DIPEA (0.14 ml, 0.8 mmol, 3.0 eq) followed by addition of tert-butyl 4-((((2-(methylamino)pyrimidin-4-yl)methyl)amino)methyl)piperidine-1-carboxylate (**Int.-S7**) (0.10 g, 0.28 mmol, 1.0 eq) at RT. The reaction mixture was stirred at RT for 16 h. After completion of reaction, the reaction mixture was poured into water, extracted with ethyl acetate (3×50 ml). The combined organic layer was dried over $Na_2SO_4$ and concentrated under vacuum. The resulting crude product was directly used in the next step (0.230 g, crude). LCMS: 47.70%, ESI-MS 653.7 m/z [M+H]+.

**Step-5: Synthesis of compound N-cyclopentyl-4-fluoro-N-(7-(((2-(methylamino)pyrimidin-4-yl)methyl)(piperidin-4-ylmethyl)amino)-7-oxoheptyl)benzamide (1)**

To a solution of tert-butyl 4-((7-(N-cyclopentyl-4-fluorobenzamido)-N-((2-(methylamino)pyrimidin-4-yl)methyl)heptanamido)methyl)piperidine-1-carboxylate (**S8**) (0.230 g, 0.3 mmol, 1.0 eq) in DCM (10 ml) was added 4N HCl in dioxane (2.5 ml) drop wise at RT. The reaction was stirred at RT for 2 h. After completion of the reaction the reaction mixture was evaporated under vacuum and it was then purified by preparative HPLC purification using the following condition: X-Bridge C18 (250*50) mm, 5 µm; eluent A: 5 mM Ammonium bicarbonate + 0.1% $NH_3$ in water; eluent B: Acetonitrile. Gradient: 80:20 at 0.01 min at flow rate: 80 ml/min, 55:45 at 18.00 min, 55:45 at 24.00 min, 00:100 at 24.01 min up to 4.0 min at flow rate: 80 ml/min; column temperature: ambient. The desired product was isolated as white solid (35 mg, 20%). LCMS: 100%, ESI-MS 553.7 m/z [M+H]+. $^1$H NMR (400 MHz, DMSO-d6, high temperature 350.5 K) δ 8.22-8.17 (d, 1 H), 7.39 (dd, J = 8.4, 5.5 Hz, 2H), 7.23 (t, J = 8.7 Hz, 2H),6.76-6.66 (d, 1H),6.396 (d,1H), 4.38 (s, 2H), 4.04 (m,1H), 3.21 (s, 4H), 3.08 (s, 4H), 2.93 (s, 2H), 2.83 (d, J = 4.9 Hz, 3H), 2.40 (d, J = 12.4 Hz, 2H), 2.28 (s, 1H), 1.75 – 1.65 (m, 6H), 1.50 (d, J = 29.2,7H), 1.26 (d, J = 15.7 Hz, 4H), 1.05 (s, 2H).

**Step-6: Synthesis of compound 2-(methylamino)pyrimidine-4-carbaldehyde (S10)**

4-(dimethoxymethyl)-N-methylpyrimidin-2-amine (**S9**) (2.0 g, 11.05 mmol, 1.0 eq) was taken in 3N HCl (20 ml). The reaction mixture was stirred at 50°C for 18 h. The progressed of the reaction was monitored by TLC. After completion of the reaction, the reaction mixture was neutralized by $NaHCO_3$ solution until pH~ 8. The product was extracted with ethyl acetate (2×200 ml). The combined organic layer was dried over $Na_2SO_4$ and concentrated under vacuum. The resulting crude product was directly used in the next step without further purification. The desired product was isolated as yellow gum (1.25 g-crude).

**Step-6a: Synthesis of compound tert-butyl 4-((((2-(methylamino)pyrimidin-4-yl)methyl)amino)methyl)piperidine-1-carboxylate (Int.-S7)**

To a solution of 2-(methylamino)pyrimidine-4-carbaldehyde (**S10**) (1.0 g, 7.26 mmol, 1.0 eq) in Methanol (10 ml) was added Acetic acid (0.05 g, 0.729 mmol, 0.1 eq) and tert-butyl 4-(aminomethyl)piperidine-1-carboxylate (**S11**) (1.75g, 8.17 mmol, 1.5 eq) at RT. The reaction mixture was stirred at RT for 3-4 h. Sodium cyanoborohydride (0.68 g, 10.93 mmol, 1.5 eq) was added to the reaction mixture at RT portion wise. After completion of the reaction, the reaction mixture was quenched in ice water (100 ml), the resulting mixture was extracted with ethyl acetate (3x100 ml). The combined organic layer was washed with brine (200 ml). It was dried over $Na_2SO_4$ and concentrated under vacuum. The crude product was purified by column chromatography on silica gel (60-120 mesh size) using Dichloromethane:Methanol (2-5%) as eluent. The desired product was isolated as semi-solid (0.65 g, 65%). LCMS: 91.2%, ESI-MS 336.36 m/z [M+H]+.

**Synthesis of N-cyclopentyl-N-(7-(ethyl((2-(methylamino)pyrimidin-4-yl)methyl)amino)-7-oxoheptyl)-4-fluorobenzamide (2) described in Scheme S3**

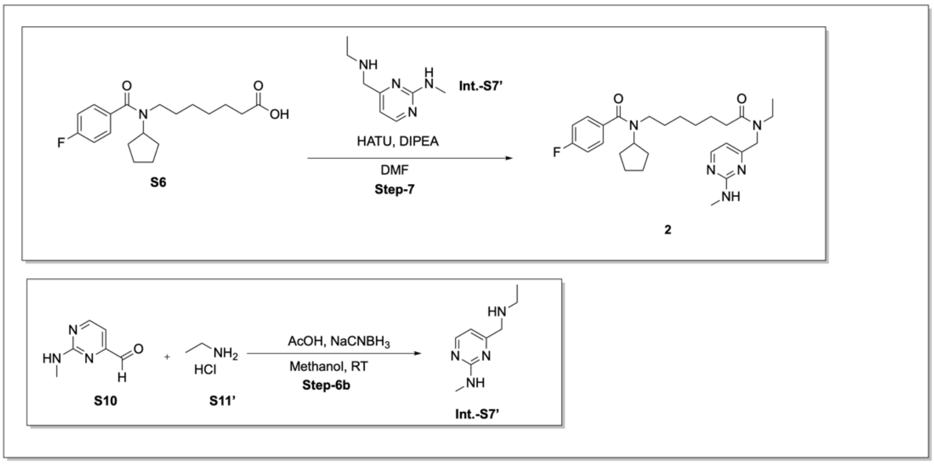

**Scheme S3**

**Step-7: Synthesis of compound N-cyclopentyl-N-(7-(ethyl((2-(methylamino)pyrimidin-4-yl)methyl)amino)-7-oxoheptyl)-4-fluorobenzamide (2)**

To a solution of 7-(N-cyclopentyl-4-fluorobenzamido)heptanoic acid (**S6**) (0.15 g, 0.44 mmol, 1.0 eq) in DMF (1 ml) was added HATU (0.168 g, 0.4 mmol, 1.5 eq), DIPEA (0.14 ml, 0.8 mmol, 3.0 eq) followed by addition of 4-((ethylamino)methyl)-N-methylpyrimidin-2-amine (**Int.-S7'**) (0.11 g, 0.67 mmol, 1.5 eq) at RT. The reaction mixture was stirred at RT for 16 h. After completion of the reaction, the reaction mixture was poured into water, extracted with ethyl acetate (3×50 ml). The combined organic layer was dried over $Na_2SO_4$ and concentrated under vacuum. The resulting crude product was purified by preparative HPLC purification using 5 mM ABC + 0.1% $NH_3$ in water:Acetonitrile as eluent. The desired product was isolated as white solid (10 mg, 10%). LCMS: 100%, ESI-MS 484.28 m/z [M+H]+. $^1$H NMR (400 MHz, DMSO-d6-High temp.350.5 K) δ 8.19 (s, 1H), 7.39 (dd, J = 8.4, 5.5 Hz, 2H), 7.23 (t, J = 8.9 Hz, 2H), 6.70 (s, 1H), 6.37 (s, 1H), 4.37 (s, 2H), 4.04-4.02 (m,1H), 3.29 (s,2H), 3.21 (s, 2H), 2.83 (d, J = 4.9 Hz, 3H), 2.35-2.34 (m, 2H), 2.27 (s, 1H), 1.85 – 1.60 (m, 6H), 1.60 – 1.39 (m, 6H), 1.28-1.13 (m, 4H), 1.09 (d, J = 30.8 Hz, 2H).

**Step-6b: Synthesis of compound 4-((ethylamino)methyl)-N-methylpyrimidin-2-amine (Int.-S7')**

To a solution of 2-(methylamino)pyrimidine-4-carbaldehyde (**S10**) (1.0 g, 7.26 mmol, 1.0 eq) in Methanol (10 ml) was added Acetic acid (0.05 g, 0.729 mmol, 0.1 eq) and Ethylamine hydrochloride (**S11'**) (0.36 g, 8.19 mmol, 1.5 eq) at RT. Reaction mixture was stirred at RT for 3-4 h. Sodium cyanoborohydride (0.68 g,10.93 mmol,1.5 eq) was added in reaction mixture at RT portion wise. After completion of reaction, the reaction mixture was quenched in ice water (100 ml), the resulting mixture was extracted with ethyl acetate (3 x 100 ml). The combined organic layer was washed with brine (200 ml), it was then dried over $Na_2SO_4$ and concentrated under vacuum. The crude product was purified by column chromatography on silica gel (60-120 mesh size) using Dichloromethane:Methanol (2-5%) as eluent. The desired product was isolated as semi-solid (0.75 g, 68%). LCMS: 72.26%, ESI-MS 167.0 m/z [M+H]+.

#### Synthesis of (3) and (4) described in Schemes S4 and S5

**Synthesis of 4-fluoro-N-isopropyl-N-(7-(((2-(methylamino)pyrimidin-4-yl)methyl)(piperidin-4-ylmethyl)amino)-7-oxoheptyl)benzamide (3)**

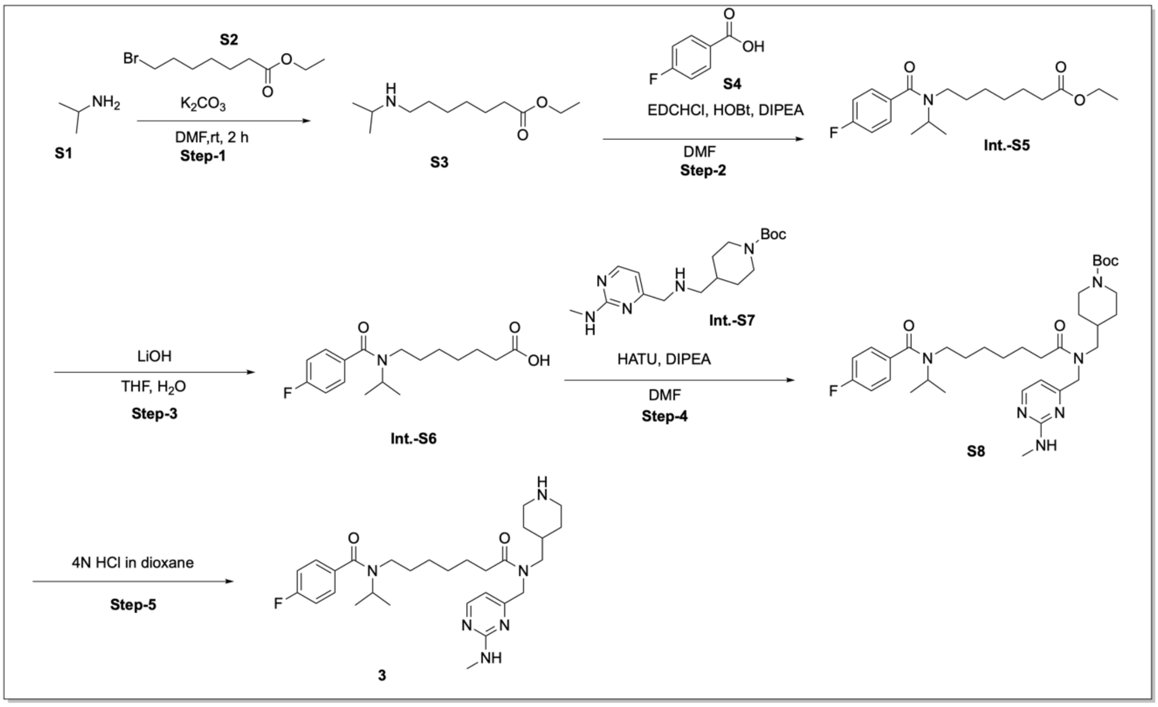

**Scheme S4**

**Step-1: Synthesis of compound Ethyl 7-(isopropylamino)heptanoate (S3)**

To a solution of Propan-2-amine (**S1**) (1.0 g, 11.7mmol, 1.0 eq) in DMF was added $K_2CO_3$ (3.2 g, 22.5 mmol, 2.0 eq) and the resulting mixture was stirred at RT for 1 h. After 1h Ethyl 7-bromoheptanoate (**S2**) (1.86 g, 7.85 mmol, 0.668 eq) was added dropwise and the resulting mixture was stirred at RT for 2.5 h. After completion of the reaction, the reaction mixture was poured into water, extracted with ethyl acetate (3×100 ml). The combined organic layer was dried over $Na_2SO_4$ and concentrated under vacuum. The crude product was purified by column chromatography on silica gel (60-120 mesh size) using Dichloromethane:Methanol (2-5%) as eluent. The desired product was isolated as solid (1.2 g, 33%). LCMS: 100%, ESI-MS 216.44 m/z [M+H]+.

**Step-2: Synthesis of compound Ethyl 7-(4-fluoro-N-isopropylbenzamido)heptanoate (Int.-S5)**

To a solution of Ethyl 7-(isopropylamino)heptanoate (**S3**) (0.650 g, 4.6 mmol, 1.0 eq) in DMF was added EDC.HCl (1.7g, 9.2 mmol, 2.0 eq), DIPEA (2.4 ml, 13.9 mmol, 3.0 eq), HOBT (0.125 g, 0.92 mmol, 0.2 eq) and 4-fluorobenzoic acid (**S4**) (0.990 g, 4.6 mmol, 1.0 eq). The mixture was stirred at RT for 16 h. After completion of the reaction, the reaction mixture was poured into water, extracted with ethyl acetate (3×100ml). The combined organic layer was dried over $Na_2SO_4$ and concentrated under vacuum. The crude product was purified by column chromatography on silica gel (60-120 mesh size) using Dichloromethane:Methanol (2-5%) as eluent. The desired product was isolated as solid (1.15 g, 73%). LCMS: 99%, ESI-MS 338.36 m/z [M+H]+.

**Step-3: Synthesis of compound 7-(4-fluoro-N-isopropylbenzamido)heptanoic acid (Int.-S6)**

To a solution of Ethyl 7-(4-fluoro-N-isopropylbenzamido)heptanoate (**Int.-S5**) (1.15 g, 3.4 mmol, 1.0 eq) in THF:$H_2O$ (1:1) (12 ml), LiOH (0.286 g, 6.8 mmol, 2.0 eq) was added. The mixture was stirred at RT for 3 h. After completion of the reaction, the reaction mixture was diluted in water (50 ml), extracted with ethyl acetate (1×50 ml). The aqueous layer was acidified with 2N HCl until pH~2. It was then extracted with ethyl acetate (3×100 ml). The combined organic layer was dried over $Na_2SO_4$ and concentrated under vacuum. The crude product was directly used in the next step without further purification (0.85g, 90%). LCMS: 100%, ESI-MS 310.30 m/z [M+H]+.

**Step-4: Synthesis of compound Tert-butyl 4-((7-(4-fluoro-N-isopropylbenzamido)-N-((2-(methylamino)pyrimidin-4-yl)methyl)heptanamido)methyl)piperidine-1-carboxylate (S8)**

To a solution of 7-(4-fluoro-N-isopropylbenzamido)heptanoic acid (**Int.-S6**) (0.10 g, 0.32 mmol, 1.0 eq) in DMF (1 ml) was added HATU (0.181 g, 0.2 mmol, 1.5 eq), DIPEA (0.14 ml,0.8 mmol, 3.0 eq) followed by addition of tert-butyl 4-((((2-(methylamino)pyrimidin-4-yl)methyl)amino)methyl)piperidine-1-carboxylate (**Int.-S7 in Scheme S2**) (0.108 g, 0.32 mmol, 1.0 eq) at RT. The reaction mixture was stirred at RT for 16 h. After completion of the reaction, the reaction mixture was poured into water, extracted with ethyl acetate (3×50 ml). The combined organic layer was dried over $Na_2SO_4$ and concentrated under vacuum. The resulting crude product was directly used in the next step (0.310 g-crude). LCMS: 73.63%, ESI-MS 627.67 m/z [M+H]+.

**Step-5: Synthesis of compound 4-Fluoro-N-isopropyl-N-(7-(((2-(methylamino)pyrimidin-4-yl)methyl)(piperidin-4-ylmethyl)amino)-7-oxoheptyl)benzamide (3)**

To a solution of tert-butyl tert-butyl 4-((7-(4-fluoro-N-isopropylbenzamido)-N-((2-(methylamino)pyrimidin-4-yl)methyl)heptanamido)methyl)piperidine-1-carboxylate (**S8**) (0.230 g, 0.3 mmol, 1.0 eq) in DCM (10 ml) was added 4N HCl in dioxane (2.5 ml) drop wise at RT. The reaction was stirred at RT for 2 h. After completion of the reaction, the reaction mixture was evaporated under vacuum and it was then purified by preparative HPLC purification using 5 mM ABC + 0.1% $NH_3$ in water:Acetonitrile as eluent. The desired product was isolated as white solid (25 mg, 25%). LCMS: 100%, ESI-MS 527.68 m/z [M+H]+. $^1$H NMR (400 MHz, DMSO-d6, high temperature 350.5 K) δ 8.23-8.17(d, 1H),7.38 (dd, J = 8.3, 5.4 Hz, 2H), 7.23 (dd, J = 10.0, 7.4 Hz, 2H), 6.77-6.66 (d, 1H),6.40 (s, 1H), 4.39 (s, 2H), 4.01 (s, 1H), 3.21 (s, 4H), 3.08 (s, 4H), 2.93 (s, 2H), 2.82 (d, J = 4.8 Hz, 3H), 2.40 (d, J = 13.6 Hz, 2H), 2.28 (s, 1H), 1.70-1.69 (m,1 H), 1.53 (s, 5H), 1.26 (d, J = 19.9 Hz, 3H), 1.17 (d, J = 6.7 Hz, 6H), 1.05 (s, 2H).

**Synthesis of N-(7-(ethyl((2-(methylamino)pyrimidin-4-yl)methyl)amino)-7-oxoheptyl)-4-fluoro-N-isopropylbenzamide (4) described in Scheme S5**

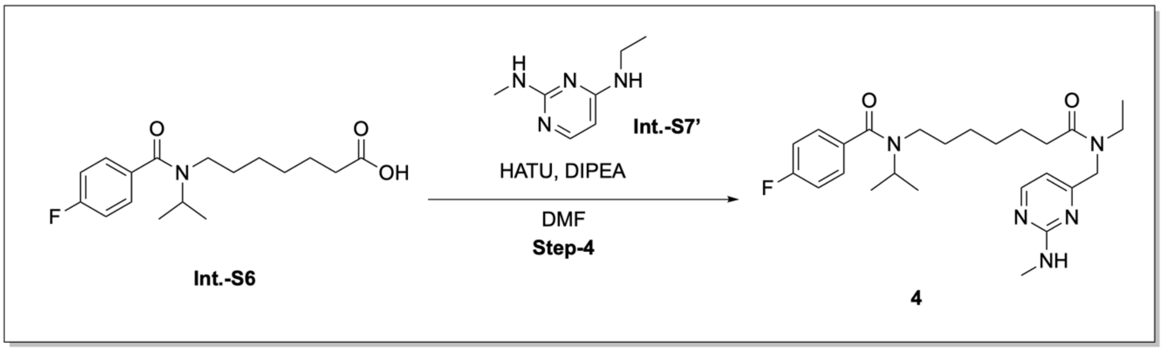

**Scheme S5**

**Step-7: Synthesis of compound N-(7-(ethyl((2-(methylamino)pyrimidin-4-yl)methyl)amino)-7-oxoheptyl)-4-fluoro-N-isopropylbenzamide (4)**

To a solution of 7-(4-fluoro-N-isopropylbenzamido)heptanoic acid (**Int.-S6 in Scheme S4**) (0.100 g, 0.32 mmol, 1.0 eq) in DMF (1 ml) was added HATU (0.168 g, 0.4 mmol, 1.5 eq), DIPEA (0.14 ml, 0.8 mmol, 3.0 eq) followed by addition of 4-((ethylamino)methyl)-N-methylpyrimidin-2-amine (**Int.-S7' in Scheme S3**) (0.11 g, 0.67 mmol, 1.5 eq) at RT. The reaction mixture was stirred at RT for 16 h. After completion of the reaction, the reaction mixture was poured into water, extracted with ethyl acetate (3×50 ml). The combined organic layer was dried over $Na_2SO_4$ and concentrated under vacuum. The resulting crude product was purified by preparative HPLC purification using 5 mM ABC + 0.1% $NH_3$ in water:Acetonitrile as eluent. The desired product was isolated as white solid (10 mg, 11%). LCMS: 100%, ESI-MS 458.57 m/z $[M+H]^+$. $^{1}$H NMR (400 MHz, DMSO-d6, high temperature 350.5 K) δ 8.19 (s, 1H), 7.38 (dd, J = 8.4, 5.5 Hz, 2H), 7.23 (t, J = 8.7 Hz, 2H), 6.70 (s, 1H), 6.37 (s, 1H), 4.37 (s, 2H), 4.01 (s, 1H), 3.39 (s, 2H), 3.20 (s, 2H), 2.83 (d, J = 4.9 Hz, 3H), 2.46-2.34 (m, 2H), 2.27 (s, 1H), 1.54 (s, 4H), 1.28 (s, 4H), 1.17 (d, J = 6.7 Hz, 6H), 1.05-0.96 (m, 2H).

#### Synthesis of (5) and (6) described in Schemes S6 and S7

**Synthesis of N-ethyl-4-fluoro-N-(7-(((2-(methylamino)pyrimidin-4-yl)methyl)(piperidin-4-ylmethyl)amino)-7-oxoheptyl)benzamide (5)**

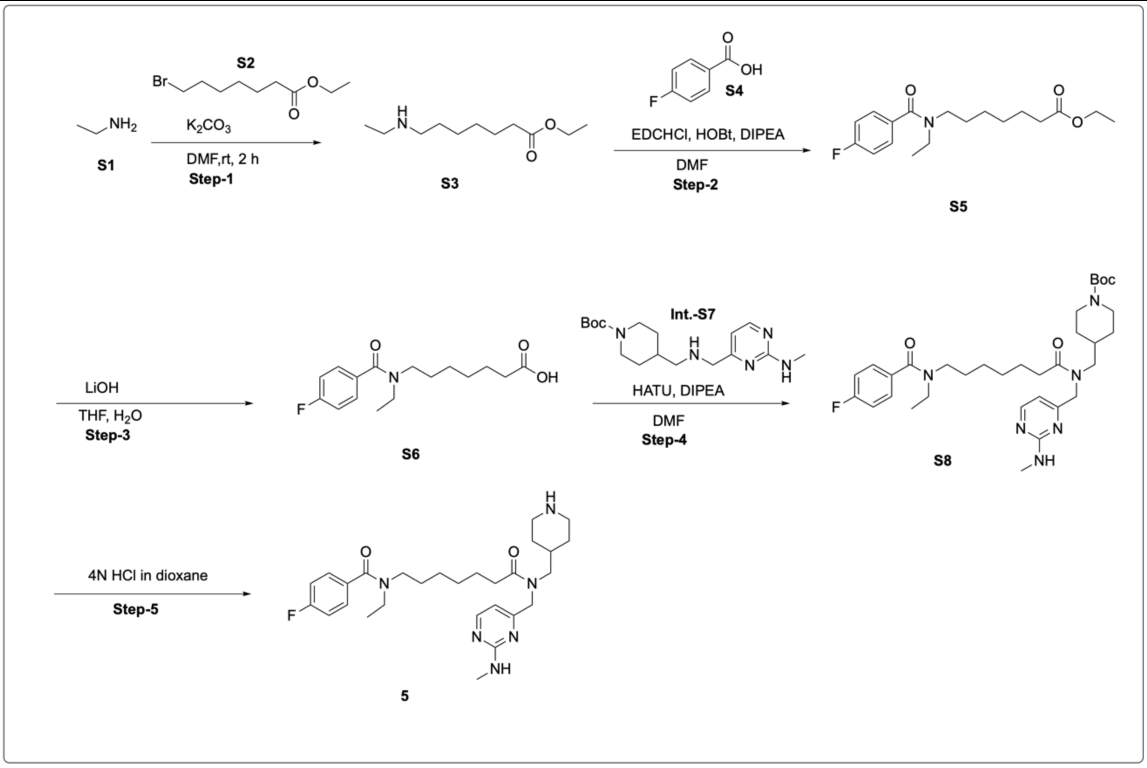

**Scheme S6**

**Step-1: Synthesis of compound Ethyl 7-(ethylamino)heptanoate (S3)**

To a solution of Ethanamine (**S1**) (3.0 g, 66 mmol, 1.0 eq) in DMF (20 ml) was added $K_2CO_3$ (18.2 g, 132 mmol, 2.0eq) and resulting mixture was stirred at RT 1 h. After 1 h Ethyl 7-bromoheptanoate (**S2**) (7.2 g, 4.44 mmol, 0.668 eq) was added dropwise and the resulting mixture was stirred at RT for 2.5 h. After completion of the reaction, the reaction mixture was poured into water, extracted with ethyl acetate (3×100ml). The combined organic layer was dried over $Na_2SO_4$ and concentrated under vacuum.

The crude product was purified by column chromatography on silica gel (60-120 mesh size) using Dichloromethane:Methanol (5-7%) as eluent. The desired product was isolated as solid (3.5 g, 35%). $^1$H NMR (400 MHz, DMSO-d6) δ 4.07-4.02 (m, 2H), 2.54-2.39 (m, 4H), 2.32-2.35 (m, 2H), 1.53-1.50 (m, 2H), 1.38-1.35 (m, 2H), 1.272 (S, 4H),1.20-1.16 (m, 3H), 1.017-0.982 (m, 3H).

**Step-2: Synthesis of compound Ethyl 7-(N-ethyl-4-fluorobenzamido)heptanoate (S5)**

To a solution of Ethyl 7-(ethylamino)heptanoate (**S3**) (1.50 g, 7.46 mmol, 1.0 eq) in DMF (15 ml) was added EDC.HCl (1.46 g, 10.44 mmol, 2.0 eq), DIPEA (2.26 g, 22.38 mmol, 3.0 eq), HOBT (0.20 g, 1.49 mmol, 0.2 eq) and 4-Fluorobenzoic acid (**S4**) (1.46 g, 10.44 mmol, 1.4 eq). The mixture was stirred at RT for 16 h. After completion of the reaction, the reaction mixture was poured into water, extracted with ethyl acetate (3×100 ml). The combined organic layer was dried over $Na_2SO_4$ and concentrated under vacuum. The crude product was purified by column chromatography on silica gel (60-120 mesh size) using Dichloromethane:Methanol (2-5%) as eluent. The desired product was isolated as solid (1.5 g, 62%). LCMS: 95%, ESI-MS 324.34 m/z [M+H]+.

**Step-3: Synthesis of compound 7-(N-ethyl-4-fluorobenzamido)heptanoic acid (S6)**

To a solution of Ethyl 7-(N-ethyl-4-fluorobenzamido)heptanoate (**S5**) (1.5 g, 4.62 mmol, 1.0 eq) in THF:$H_2O$ (1:1) (20 ml), LiOH (0.45 g, 9.2 mmol, 2.0 eq) was added. The mixture was stirred at RT for 3 h. After completion of the reaction, the reaction mixture was diluted in water (50 ml), extracted with ethyl acetate (1×50 ml). The aqueous layer was acidified with 2N HCl until pH~2, It was then extracted with ethyl acetate (3×100 ml). The combined organic layer was dried over $Na_2SO_4$ and concentrated under vacuum. The crude product was directly used in the next step without further purification (1.2 g, 87%). LCMS: 100%, ESI-MS 296.34 m/z [M+H]+.

**Step-4: Synthesis of compound Tert-butyl 4-((7-(N-ethyl-4-fluorobenzamido)-N-((2-(methylamino)pyrimidin-4-yl)methyl)heptanamido)methyl)piperidine-1-carboxylate (S8)**

To a solution of 7-(N-ethyl-4-fluorobenzamido)heptanoic acid (**S6**) (0.200 g, 0.6 mmol, 1.0 eq) in DMF (5 ml) was added HATU (0.386 g, 1.0 mmol, 1.5 eq), DIPEA (0.14 ml, 0.8 mmol, 3.0 eq) followed by addition of tert-butyl 4-((((2-(methylamino)pyrimidin-4-yl)methyl)amino)methyl)piperidine-1-carboxylate (**Int.-S7**) (0.340 g, 1.01 mmol, 1.5 eq). The reaction mixture was stirred at RT for 16 h. After completion of the reaction, the reaction mixture was poured into water, extracted with ethyl acetate (3×50 ml). The combined organic layer was dried over $Na_2SO_4$ and concentrated under vacuum. The resulting crude product was directly used in the next step (0.22 g, crude). LCMS: 84.20%, ESI-MS 613.39 m/z [M+H]+.

**Step-5: Synthesis of compound N-ethyl-4-fluoro-N-(7-(((2-(methylamino)pyrimidin-4-yl)methyl)(piperidin-4-ylmethyl)amino)-7-oxoheptyl)benzamide (5)**

To a solution of tert-butyl 4-((7-(N-ethyl-4-fluorobenzamido)-N-((2-(methylamino)pyrimidin-4-yl)methyl)heptanamido)methyl)piperidine-1-carboxylate (**S8**) (0.22 g, 0.30 mmol, 1.0 eq) in DCM (10 ml) was added 4N HCl in dioxane (2.5 ml) drop wise at RT. The reaction was stirred at RT for 2 h. After completion of the reaction, the reaction mixture was evaporated under vacuum and it was purified by preparative HPLC using 5 mM ABC + 0.1% $NH_3$ in water:Acetonitrile as eluent. The desired product was isolated as white solid (40 mg, 25%). LCMS: 100%, ESI-MS 513.18 m/z [M+H]+. $^1$H NMR (400 MHz, DMSO-d6-High temp.350.5 K) δ 8.22 (s, 1H), 7.40 (t, J = 6.9 Hz, 2H), 7.23 (t, J = 8.8 Hz, 2H), 6.37 (s, 1H), 6.39 (s, 1H), 4.38 (s, 2H), 3.32 (m, 4H), 3.22 (d, J = 7.1 Hz, 2H), 3.078 (s, 4H), 2.94 (d, J = 12.1 Hz, 2H), 2.82 (d, J = 4.8 Hz, 3H), 2.45 (d, J = 12.9 Hz, 2H), 2.27 (s, 1H), 1.70 (m, 1H), 1.53 (s, 5H), 1.23 (s, 3H), 1.10 (t, J = 7.1 Hz, 3H), 1.08 (m, 1H).

**Step-7: Synthesis of compound (6)**

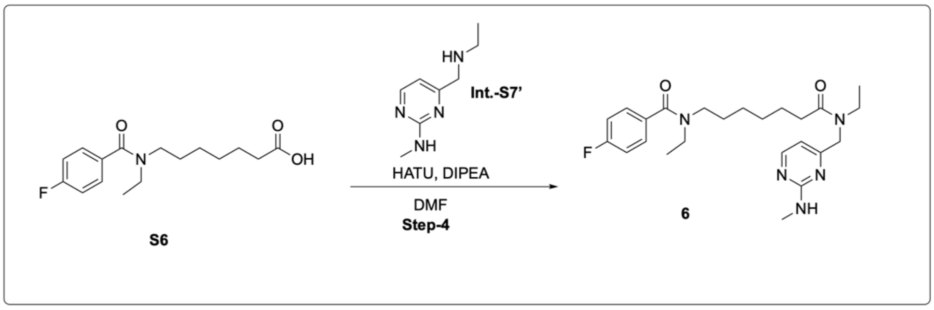

**Scheme S7**

To a solution of 7-(N-ethyl-4-fluorobenzamido)heptanoic acid (**S6**) (0.100 g, 0.32 mmol, 1.0 eq) in DMF (1 ml) was added HATU (0.168 g, 0.4 mmol, 1.5 eq), DIPEA (0.14 ml, 0.8 mmol, 3.0 eq) followed by addition of 4-((ethylamino)methyl)-N-methylpyrimidin-2-amine (**Int.-S7' in Scheme S3**) (0.11 g, 0.67 mmol, 1.5 eq) at RT. The reaction mixture was stirred at RT for 16 h. After completion of the reaction, the reaction mixture was poured into water, extracted with ethyl acetate (3×50 ml). The combined organic layer was dried over $Na_2SO_4$ and concentrated under vacuum. The resulting crude product was purified by preparative HPLC using 5 mM ABC + 0.1% NH3 in water:Acetonitrile as eluent. The desired product was isolated as white solid (10 mg, 11%). LCMS: 100%, ESI-MS 444.02 m/z [M+H]+. 1H NMR (400 MHz, DMSO-d6-High temp.350.5 K) δ 8.20 (s, 1H), 7.40 (t, J = 6.9 Hz, 2H), 7.23 (t, J = 8.7 Hz, 2H), 6.69 (s, 1H), 6.37 (s, 1H), 4.37 (s, 2H), 3.39-3.30 (m, 5H), 2.83 (d, J = 4.9 Hz, 3H), 2.27 (s, 1H), 1.54 (s, 4H), 1.28 (s, 7H), 1.11 (t, J = 7.0 Hz, 5H).

#### Synthesis of (7) and (8) described in Schemes S8 and S9

**Synthesis of 7-((4-benzyl-5-isobutyl-4H-1,2,4-triazol-3-yl)thio)-N-((2-(methylamino)pyrimidin-4-yl)methyl)-N-(piperidin-4-ylmethyl)heptanamide (7)**

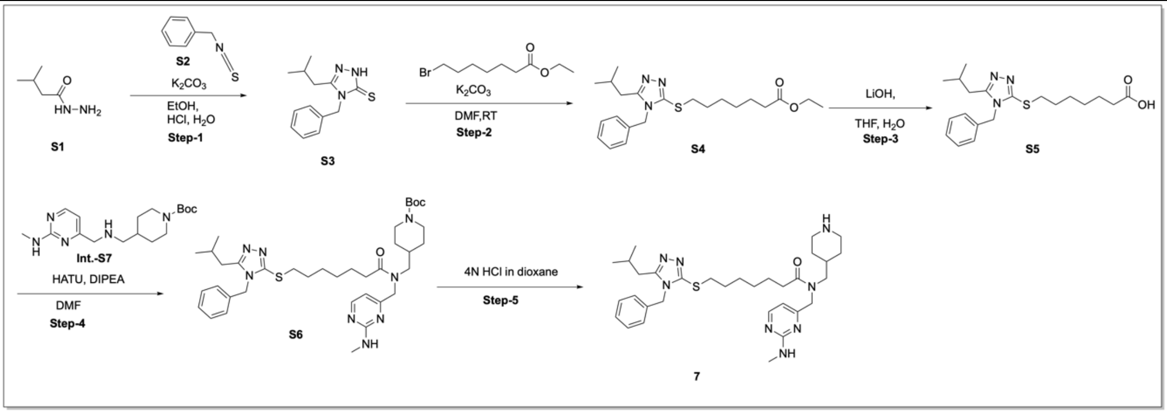

**Scheme S8**

**Step-1: Synthesis of compound 4-Benzyl-5-isobutyl-2,4-dihydro-3H-1,2,4-triazole-3-thione (S3)**

Isopentylhydrazine (**S1**) (0.600 g, 5.1 mmol, 1.0 eq) and (Isothiocyanatomethyl)benzene (S2) (0.770 g, 5.1 mmol, 1.0 eq) were taken in ethanol (10 ml) and refluxed for 2 h. The reaction mixture was cooled to RT and $K_2CO_3$ (0.650 g, 5.1 mmol, 1.0 eq) was added in one portion. The reaction mixture was then refluxed for another 1 h. After completion of the reaction, the reaction mixture was poured into water and neutralized by 1N HCl until pH~7. The resulting solid was filtered and washed with water (50 ml). The desired product was isolated as solid (0.735 g, 50%). LCMS: 100%, ESI-MS 248.1 m/z [M+H]+.

$^1$H NMR (400 MHz, DMSO-d6) δ, 7.31-7.39 (m,3H), 7.27 (d, J =4Hz ,2H), 5.30 (s, 2H), 2.369 (d, J =7.2 Hz, 2H), 1.80-2.02 (m,1H), 0.753 (d, J =6.4 Hz, 6H).

**Step-2: Synthesis of compound Ethyl 7-((4-benzyl-5-isobutyl-4H-1,2,4-triazol-3-yl)thio)heptanoate (S4)**

To a solution of 4-Benzyl-5-isobutyl-2,4-dihydro-3H-1,2,4-triazole-3-thione (**S3**) (0.300 g, 1.20 mmol, 1.0 eq) and Ethyl 7-bromoheptanoate (0.287g,1.20 mmol,1.0 eq) in DMF (10 ml) was added $K_2CO_3$ (0.225 g, 1.20 mmol, 1.0 eq) at RT. The reaction mixture was stirred at 50°C for 16 h. After completion of reaction, the reaction mixture was poured into water and extracted with ethyl acetate (3×100 ml). The combined organic layer was dried over $Na_2SO_4$ and concentrated under vacuum to give crude product. The desired product was isolated as solid (0.40 g, 84.00%). LCMS: 97.20%, ESI-MS 404.4 m/z [M+H]+. $^1$H NMR (400 MHz, DMSO-D6) δ 7.303-7.359 (m, 3H), 7.223 (d, *J* = 6.8Hz,2H), 5.303(s,2H) , 4.145 (s, 2H), 4.036 (d, *J* =6.8 Hz ,2H), 2.337(s, 2H), 2.241 (t, *J* =6.8 Hz, 2H), 1.763 (s,4H), 1.507 (s, 2H), 1.288 (s, 3H), 1.157 (t, *J* =6.4 Hz, 3H), 0.798 (d, *J* =6.0 Hz, 6H).

**Step-3: Synthesis of compound 7-((4-Benzyl-5-isobutyl-4H-1,2,4-triazol-3-yl)thio)heptanoic acid (S5)**

To a solution of Ethyl 7-((4-benzyl-5-isobutyl-4H-1,2,4-triazol-3-yl)thio)heptanoate (**S4**) (0.400 g, 0.99 mmol, 1.0 eq) in THF:$H_2O$ (1:1) (16 ml), LiOH (0.25 g, 1.8 mmol, 2.0 eq) was added. The mixture was stirred at RT for 3 h. After completion of the reaction, the reaction mixture was diluted with water (50 ml), extracted with ethyl acetate (1×50 ml). The aqueous layer was acidified with 2N HCl until pH~2, It was then extracted with ethyl acetate (3×100 ml). The combined organic layer was dried over $Na_2SO_4$ and concentrated under vacuum. The crude product was directly used in the next step without further purification (0.350 g, 94%). LCMS: 88%, ESI-MS 375.5.

**Step-4: Synthesis of compound Tert-butyl4-((7-((4-benzyl-5-isobutyl-4H-1,2,4-triazol-3-yl)thio)-N-((2-(methylamino)pyrimidin-4-yl)methyl)heptanamido)methyl)piperidine-1-carboxylate (S6)**

To a solution of 7-((4-benzyl-5-isobutyl-4H-1,2,4-triazol-3-yl)thio)heptanoic acid (**S5**) (0.200 g, 0.53 mmol, 1.0 eq) in DMF was added HATU (0.300g, 0.798 mmol, 1.5 eq), DIPEA (0.51ml, 0.16 mmol, 3.0 eq) followed by addition of tert-butyl 4-((((2-(methylamino)pyrimidin-4-yl)methyl)amino)methyl)piperidine-1-carboxylate (**Int.-S7 in Scheme S2**) (0.214 g, 0.639 mmol, 1.2 eq). The reaction mixture was stirred at RT for 16 h. After completion of reaction, the reaction mixture was poured into water, extracted with ethyl acetate (3×50 ml). The combined organic layer was dried over $Na_2SO_4$ and concentrated under vacuum. The resulting crude product was directly used in the next step (0.190g, 48%). LCMS: 100%, ESI-MS 694.0 m/z [M+H]+.

**Step-5: Synthesis of compound 7-((4-benzyl-5-isobutyl-4H-1,2,4-triazol-3-yl)thio)-N-((2-(methylamino)pyrimidin-4-yl)methyl)-N-(piperidin-4-ylmethyl)heptanamide (7)**

To a solution of tert-butyl 4-((7-((4-benzyl-5-isobutyl-4H-1,2,4-triazol-3-yl)thio)-N-((2-(methylamino)pyrimidin-4-yl)methyl)heptanamido)methyl)piperidine-1-carboxylate (**S6**) (0.190g, 0.14 mmol, 1.0 eq) in DCM (10 ml) was added 4N HCl in dioxane (2.5 ml) drop wise at RT. The reaction was stirred at RT for 2 h. After completion of the reaction, the reaction mixture was evaporated under vacuum and it was then purified by preparative HPLC using 5 mM ABC + 0.1% $NH_3$ in water:Acetonitrile as eluent. The desired product was isolated as white solid (10 mg, 22%). LCMS: 100% ESI-MS 593.2 m/z $[M+H]^+$. $^1$H NMR (400 MHz, DMSO-d6, high temperature 350.5 K) δ 8.22-8.13(d,1H), 7.41 – 7.27 (m, 3H), 7.07 (d, J = 7.4 Hz, 2H), 6.77 (s, 1H), 6.37 (d, J = 19.5 Hz, 1H), 5.18 (s, 2H), 4.39 (s, 2H), 3.22 (d, J = 7.0 Hz, 2H), 3.08-3.04 (s, 4H), 2.93 (s, 2H), 2.82 (d, J = 4.8 Hz, 3H),

2.41 (s, 2H), 2.38 (s, 2H), 2.28 (s, 1H), 1.99 (dt, J = 13.4, 6.7 Hz, 1H), 1.57 (d, 8H), 1.51 (s,4H) 1.05 (s, 2H), 0.90 (d, J = 6.6 Hz, 6H).

**Synthesis of 7-((4-benzyl-5-isobutyl-4H-1,2,4-triazol-3-yl)thio)-N-ethyl-N-((2-(methylamino)pyrimidin-4-yl)methyl)heptanamide (8) described in Scheme S9**

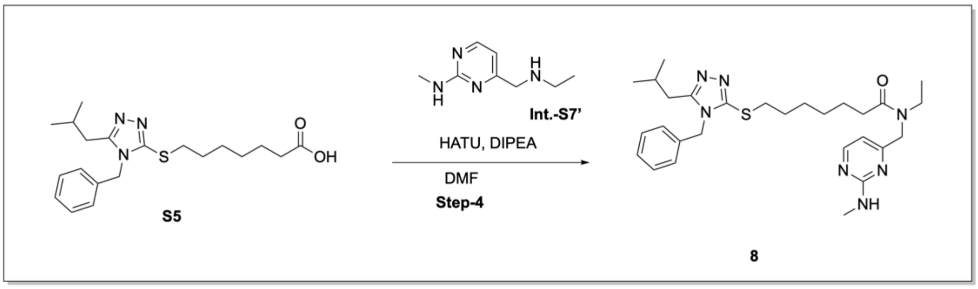

**Scheme S9**

**Step-5: Synthesis of compound 7-((4-benzyl-5-isobutyl-4H-1,2,4-triazol-3-yl)thio)-N-ethyl-N-((2-(methylamino)pyrimidin-4-yl)methyl)heptanamide (8)**

To a solution of 7-((4-benzyl-5-isobutyl-4H-1,2,4-triazol-3-yl)thio)heptanoic acid (**S5**) (0.100 g, 0.266 mmol, 1.0 eq) in DMF (5 ml) was added HATU (0.151 g, 0.399 mmol, 1.5 eq), DIPEA (0.14 ml, 0.8 mmol, 3.0 eq) followed by addition of 4-((ethylamino)methyl)-N-methylpyrimidin-2-amine (**Int.-S7' in Scheme S3**) (0.053 g, 0.319 mmol, 1.2 eq). The reaction mixture was stirred at RT for 16 h. After completion of the reaction, the reaction mixture was poured into water, extracted with ethyl acetate (3×50 ml). The combined organic layer was dried over $Na_2SO_4$ and concentrated under vacuum. The resulting crude product was purified by preparative HPLC using 5 mM ABC + 0.1% $NH_3$ in water:Acetonitrile as eluent. The desired product was isolated as white solid (10 mg, 8%). LCMS: 100%, ESI-MS 524.0 m/z [M+H]+. $^1$H NMR (400 MHz, DMSO-d6, high temperature 350.5 K) 8.19 (d, 1H), 7.43 – 7.26 (m, 3H), 7.07 (d, J = 7.4 Hz, 2H), 6.76-6.69(d, 1H), 6.36 (d,1H), 5.18 (s, 2H), 4.37 (s, 2H), 3.39 (s, 2H), 2.83 (d, J = 4.9 Hz, 3H), 2.34 (s,2H), 2.28(m, 1H), 2.02-1.97(m, 1H) 1.58 (d, J = 34.5 Hz, 4H), 1.28 (s, 6H), 1.12 (s, 4H), 0.90 (d, J = 6.7 Hz, 6H).

#### Synthesis of (9) described in Scheme S10

**Synthesis of Methyl 4-((N-ethyl-7-(4-fluoro-N-isopropylbenzamido)heptanamido)methyl)benzoate (9)**

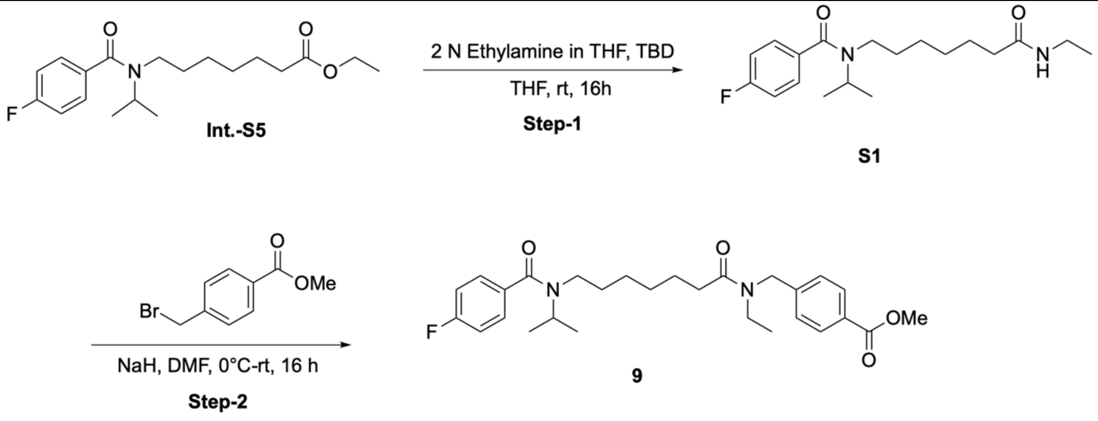

**Scheme S10**

**Synthesis of N-(7-(ethylamino)-7-oxoheptyl)-4-fluoro-N-isopropylbenzamide (S1)**

To a solution of ethyl 7-(4-fluoro-N-isopropylbenzamido)heptanoate (**Int.-S5 in Scheme S4**) (2 g, 5.93 mmol) in THF (2 ml) was added Ethylamine (2M in THF, 20 mL) followed by TBD (1.64 g, 11.86 mmol) at 0°C and the reaction mixture was stirred for 16 h at RT (monitored by TLC). The reaction mixture was quenched with water (5 ml) and extracted with ethyl acetate (3x10 ml). The combined organic extracts were dried over $Na_2SO_4$, evaporated and purified by column chromatography (1:1 ethyl acetate in hexane) to get N-(7-(ethylamino)-7-oxoheptyl)-4-fluoro-N-isopropylbenzamide (**S1**) (1.8 g 90% yield). LCMS: 337.7 $[M+H]^+$.

**Synthesis of Methyl 4-((N-ethyl-7-(4-fluoro-N-isopropylbenzamido)heptanamido)methyl)benzoate (9):**

To a solution of N-(7-(ethylamino)-7-oxoheptyl)-4-fluoro-N-isopropylbenzamide (**S1**) (0.5 g, 1.49 mmol) in DMF (5 ml) was added NaH (118 mg, 2.98 mmol) at 0°C. The reaction mixture was stirred for 30 minutes at 0°C and methyl 4-(bromomethyl)benzoate (682 mg, 2.98 mmol) was added. The reaction mixture was stirred at RT for 16 h (monitored by TLC) and quenched by water (20 ml), extracted with ethyl acetate (3 x 20 ml). The combined organic extracts were dried over $Na_2SO_4$, evaporated and purified by column chromatography eluting with 50% ethyl acetate in hexane to get methyl 4-((N-ethyl-7-(4-fluoro-N-isopropylbenzamido)heptanamido)methyl)benzoate (**9**) (80 mg, 12% yield) as a colorless sticky material. LCMS: 485.4 $[M+H]^+$, HPLC: 100%, $^1$H NMR (400MHz, $CDCl_3$) δ = 8.03-8.00 (m, 2H), 7.40 – 7.30 (m, 4H), 7.14 – 7.09 (m, 2H), 4.67 – 4.61 (m, 2H), 3.96 (m, 4H), 3.50 (q, *J* = 7.2 Hz, 1H), 3.33 (m, 4H), 2.44 (bs, 1H), 2.31 (bs, 1H), 1.70 (m, 2H), 1.47 – 1.38 (m, 4H), 1.29 – 1.13 (m, 10H).

#### Synthesis of (10) described in Scheme S11

**Synthesis of Methyl 4-(4-((N-ethyl-7-(4-fluoro-N-isopropylbenzamido)heptanamido)methyl)piperidin-1-yl)benzoate (10)**

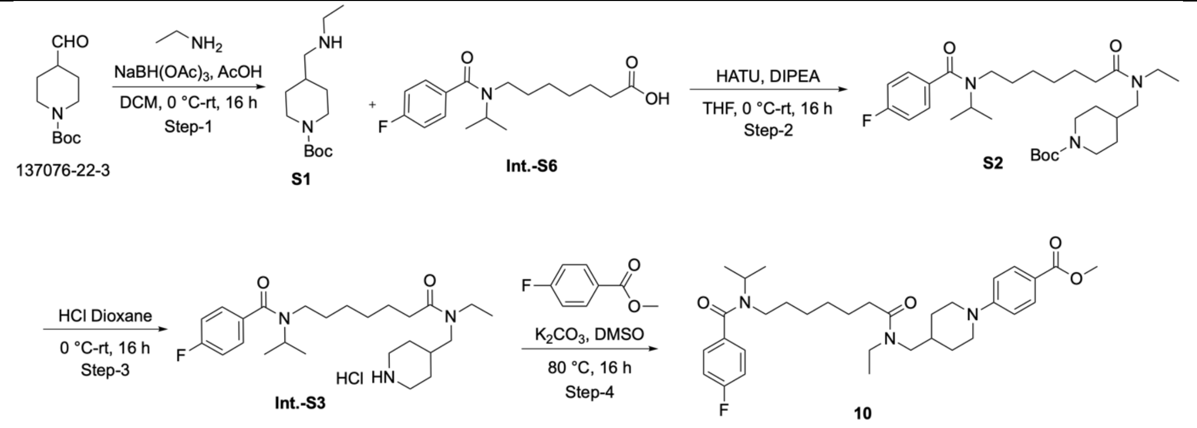

**Scheme S11**

**Synthesis of Tert-butyl 4-((ethylamino)methyl)piperidine-1-carboxylate (S1)**

To a solution of tert-butyl 4-formylpiperidine-1-carboxylate (18 g, 84.5 mmol) in THF (200 ml) was added ethylamine (42.25 ml, 84.5 mmol) at 0°C followed by drop of acetic acid (cat.) and the reaction mixture was stirred for 1 h at RT (monitored by TLC). The reaction mixture cooled to 0 °C and $NaBH(OAc)_3$ (17.91 g, 84.5 mmol) was added and stirred at RT for 16 h. The reaction mixture quenched with saturated $NaHCO_3$ and organic solvent is removed under reduced pressure. The aqueous layer was washed with ethyl acetate once to remove non-polar impurities (discarded). The aqueous layer is further extracted with 5% MeOH : DCM (3 x 200 ml). The combined layers of MeOH: DCM extracts were dried over sodium sulphate, evaporated to dryness to get pure tert-butyl 4-((ethylamino)methyl)piperidine-1-carboxylate (**S1**) (15 g, 73% yield) as a colorless oil. LCMS: 243.7 $[M+H]^+$.

**Synthesis of Tert-butyl 4-((N-ethyl-7-(4-fluoro-N-isopropylbenzamido)heptanamido)methyl)piperidine-1-carboxylate (S2):**

To a stirred solution of 7-(4-fluoro-N-isopropylbenzamido)heptanoic acid (**Int.-S6 in Scheme S4**) (3 g, 9.7 mmol) in dry THF (30 ml) was added HATU (5.5 g, 14.5 mmol) at 0 °C and DIPEA (8.5 ml, 48.0 mmol). The reaction mixture was stirred at RT for 1 h and tert-butyl 4-((ethylamino)methyl)piperidine-1-carboxylate (**S1**) (4.2 g, 17.4 mmol) was added. The reaction mixture was stirred at RT for 16 h (monitored by TLC) and quenched with water (20 ml). The aqueous layer was extracted with ethyl acetate (3 x 50 ml) and combined organic layer was evaporated to dryness to get tert-butyl 4-((N-ethyl-7-(4-fluoro-N-isopropylbenzamido)heptanamido)methyl)piperidine-1-carboxylate (**S2**) (5.1 g, quantitative yield) as colorless sticky material. LCMS: 534.6 $[M+H]^+$ .

**Synthesis of Methyl 4-(4-((N-ethyl-7-(4-fluoro-N-isopropylbenzamido)heptanamido)methyl)piperidin-1-yl)benzoate (10):**

To a stirred solution of tert-butyl 4-((N-ethyl-7-(4-fluoro-N-isopropylbenzamido)heptanamido)methyl)piperidine-1-carboxylate (**S2**) (10 g, 23.0 mmol) in dry DCM (20 ml) was added 4M HCl in dioxane (30 ml) at 0°C and reaction mixture was stirred for 4 h (monitored by TLC). The solvent was removed under reduced pressure and resulting HCl salt (**Int.-S3**) (10 g) was used for next step without further purification.

To a solution of above crude (10 g) in DMSO (100 ml) was added $K_2CO_3$ (15.8 g, 115.2 mmol) (pH~9) followed by methyl 4-fluorobenzoate (7.09 g, 46.0 mmol). The reaction mixture was heated to 80°C for 16 h (monitored by TLC), quenched with cold water and extracted with ethyl acetate (3 x 50 ml). The combined organic extracts were dried, evaporated and purified by flash column chromatography eluting with 15% MeOH: DCM to product. The obtained product was further purified using 0.1% formic acid in water: acetonitrile to get methyl 4-(4-((N-ethyl-7-(4-fluoro-N-isopropylbenzamido)heptanamido)methyl)piperidin-1-yl)benzoate (**10**) (3.3 g, 31% yield over two step) as a white solid. LCMS: 568.5 $[M+H]^+$, HPLC: 99.95%, $^1$H NMR (400MHz, DMSO-d6) δ = 7.76 (d, $J$ = 6 Hz, 2H), 7.40 (m, 2H), 7.29-7.23 (m, 2H), 6.97 (m, 2H), 3.98 - 3.89 (m, 2H), 3.76 (s, 3H), 3.73 (m, 1H), 3.16 (m, 3H), 2.82 (t, $J$ = 12 Hz, 2H), 2.33 (m, 3H), 1.85 (s, 1H), 1.67-1.58 (m, 6H), 1.42-1.31 (m, 4H), 1.23-0.98 (m, 13H).

#### Synthesis of (11), (12), (13) as described in Scheme S12

**Syntheses of Methyl 4-(4-((N-ethyl-7-(4-fluoro-N-isopropylbenzamido)heptanamido)methyl)piperidin-1-yl)-2-nitrobenzoate (11), 4-(4-((N-ethyl-7-(4-fluoro-N-isopropylbenzamido)heptanamido)methyl)piperidin-1-yl)-2-nitrobenzoic acid (12) and 2-amino-4-(4-((N-ethyl-7-(4-fluoro-N-isopropylbenzamido)heptanamido)methyl)piperidin-1-yl)benzoic acid (13)**

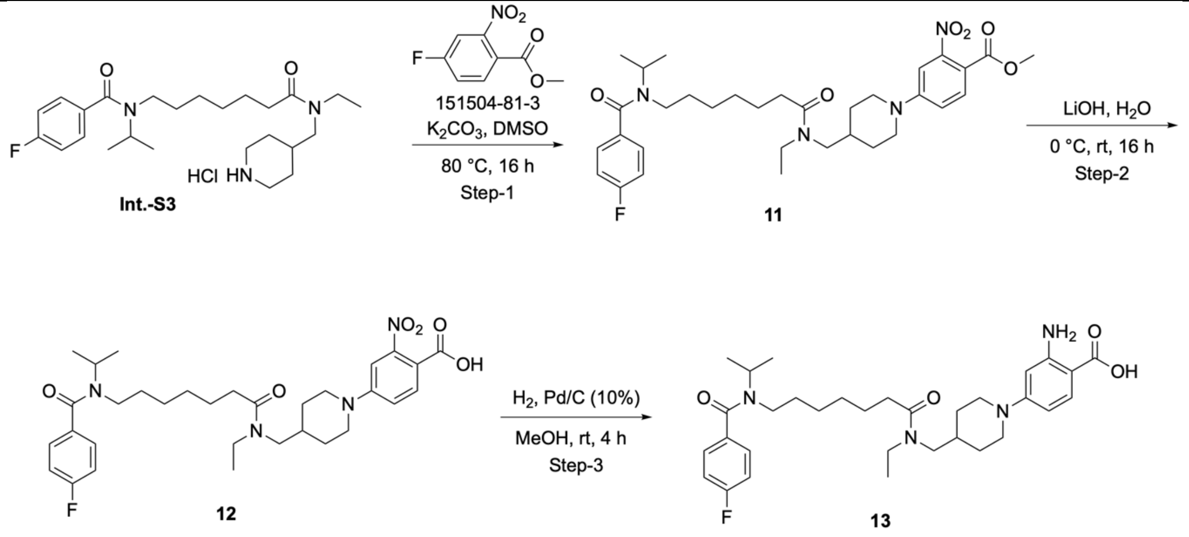

**Scheme S12**

**Synthesis of Methyl 4-(4-((N-ethyl-7-(4-fluoro-N-isopropylbenzamido)heptanamido)methyl)piperidin-1-yl)-2-nitrobenzoate (11)**

To a solution N-(7-(ethyl(piperidin-4-ylmethyl)amino)-7-oxoheptyl)-4-fluoro-N-isopropylbenzamide hydrochloride (**Int.-S3 in Scheme S11**) (300 mg, 0.69 mmol) in DMSO (3 ml) was added $K_2CO_3$ (191 mg, 1.38 mmol) (pH~9) followed by methyl 4-fluoro-2-nitrobenzoate (165 mg, 0.82 mmol). The reaction mixture was heated to 80°C for 16 h (monitored by TLC). Quenched with cold water and extracted with ethyl acetate (3 x 10 ml). The combined organic extracts were dried, evaporated and purified by flash column chromatography eluting with ethyl acetate in hexane (7:3) to get methyl 4-(4-((N-ethyl-7-(4-fluoro-N-isopropylbenzamido)heptanamido)methyl)piperidin-1-yl)-2-nitrobenzoate (**11**) (150 mg, 38% yield) as a colorless sticky material. LCMS: 613.5 $[M+H]^+$, HPLC: 100%, $^1H$ NMR (400MHz, DMSO-d6) δ = 7.73 (d, *J* = 8.8 Hz, 1H), 7.40 (m, 2H), 7.31-7.23 (m, 3H), 7.12 (m, 1H), 4.06 (m, 2H), 3.74 (m, 4H), 3.29 (m, 2H), 3.16 (m, 4H), 2.92 (t, *J* = 12 Hz, 2H), 2.33 (m, 3H), 1.89 (s, 1H), 1.66-1.58 (m, 5H), 1.31 (m, 3H), 1.23-0.97 (m, 12H).

**Synthesis of 4-(4-((N-ethyl-7-(4-fluoro-N-isopropylbenzamido)heptanamido)methyl)piperidin-1-yl)-2-nitrobenzoic acid (12)**

Methyl 4-(4-((N-ethyl-7-(4-fluoro-N-isopropylbenzamido)heptanamido)methyl)piperidin-1-yl)-2-nitrobenzoate (**11**) (280 mg, 0.4 mmol) in THF:MeOH:Water (9 ml, 1:1:1) and $LiOH{\cdot}H_2O$ (96 mg, 2.2 mmol) at 0°C was added. The reaction mixture was stirred at RT for 16 h (monitored by TLC) and the solvent was removed under reduced pressure. The residue was dissolved in water (1 ml) and acidified to pH~ 3 at 0°C. The aqueous layer is extracted with ethyl acetate (3 x 5 ml) and combined organic layer was evaporated to dryness. The obtained semisolid was further purified by RP-preparative-HPLC using 0.1% formic acid in water and acetonitrile to get pure 4-(4-((N-ethyl-7-(4-fluoro-N-isopropylbenzamido)heptanamido)methyl)piperidin-1-yl)-2-nitrobenzoic acid (**12**) (150 mg, 55% yield) as a white solid. LCMS: 599.5 $[M+H]^+$, HPLC: 100%, $^1H$ NMR (400MHz, DMSO-d6) δ = 7.74 (d, *J* = 8.8 Hz, 1H), 7.40 (m, 2H), 7.33-7.24 (m, 3H), 7.10 (m, 1H), 4.05 (m, 2H), 3.78 (s, 1H), 3.18 (m, 4H), 2.91 (t, *J* = 12.4 Hz, 2H), 2.34 (m, 2H), 1.90 (bs, 1H), 1.67-1.58 (m, 6H), 1.33 (m, 3H), 1.25-1.02 (m, 14H). (Note: acid proton not observed in $^1H$ NMR).

**Synthesis of 2-amino-4-(4-((N-ethyl-7-(4-fluoro-N-isopropylbenzamido)heptanamido)methyl)piperidin-1-yl)benzoic acid (13)**

To a stirred solution of 4-(4-((N-ethyl-7-(4-fluoro-N-isopropylbenzamido)heptanamido)methyl)piperidin-1-yl)-2-nitrobenzoic acid (**12**) (120 mg, 0.02 mmol) in MeOH (3 ml), 10% Pd/C (50 mg, 50% moisture) was added. The reaction mixture was stirred at RT for 4 h (monitored by TLC) and catalyst was filtered out by using celite bed. The filtrate was evaporated under reduced pressure and residue was purified by RP-preparative-HPLC using 0.1% formic acid in water and acetonitrile to get 2-amino-4-(4-((N-ethyl-7-(4-fluoro-N-isopropylbenzamido)heptanamido)methyl)piperidin-1-yl)benzoic acid (**13**) (60 mg, 53% yield) as a white solid. LCMS: 569.5 $[M+H]^+$, HPLC: 100%, $^1$H NMR (400MHz, DMSO-d6) δ = 7.51 (dd, $J$ = 8.8, 2.4 Hz, 1H), 7.40 (m, 2H), 7.29-7.23 (m, 2H), 6.18 (m, 2H), 4.85 (bs, 1H), 4.42 (m, 1H), 3.83 (m, 2H), 3.17 (m, 5H), 2.75 (t, $J$ = 10 Hz, 2H), 2.34 (m, 2H), 1.83 (s, 1H), 1.67-1.57 (m, 6H), 1.33 (bs, 3H), 1.25-1.01 (m, 14H). (Note: acid proton not observed in $^1$H NMR).

#### Synthesis of (14) and (15) as described in Scheme S13

**Syntheses of Methyl 4-(4-((N-ethyl-7-(4-fluoro-N-isopropylbenzamido)heptanamido)methyl)piperidin-1-yl)-2-nitrobenzoate (Int-1) and 4-(4-((N-ethyl-7-(4-fluoro-N-isopropylbenzamido)heptanamido)methyl)piperidin-1-yl)-3-nitrobenzoic acid (14) and Methyl 3-amino-4-(4-((N-ethyl-7-(4-fluoro-N-isopropylbenzamido)heptanamido)methyl)piperidin-1-yl)benzoate (15)**

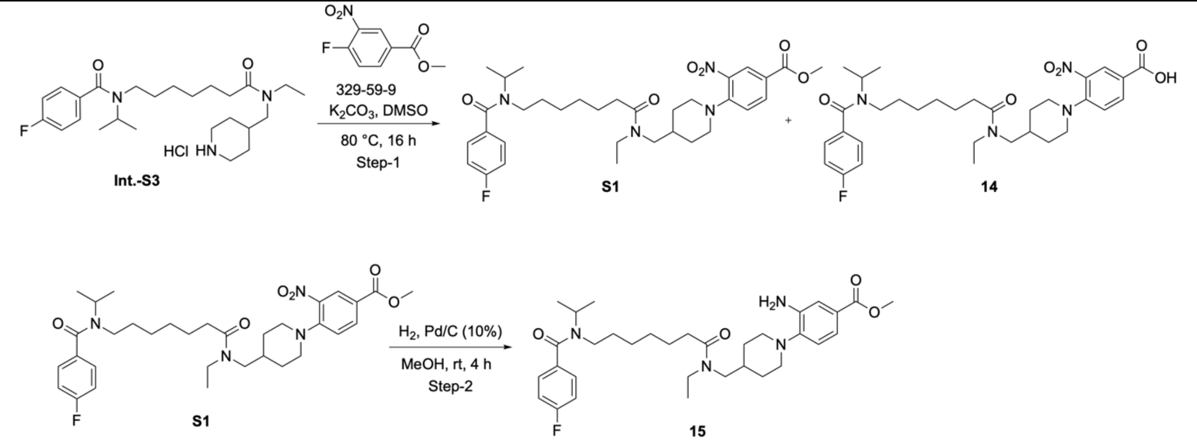

**Scheme S13**

**Synthesis of Methyl 4-(4-((N-ethyl-7-(4-fluoro-N-isopropylbenzamido)heptanamido)methyl)piperidin-1-yl)-2-nitrobenzoate (Int-1) and 4-(4-((N-ethyl-7-(4-fluoro-N-isopropylbenzamido)heptanamido)methyl)piperidin-1-yl)-3-nitrobenzoic acid (14)**

To a solution of N-(7-(ethyl(piperidin-4-ylmethyl)amino)-7-oxoheptyl)-4-fluoro-N-isopropylbenzamide hydrochloride (**Int.-S3 in Scheme S11**) (300 mg, 0.69 mmol) in DMSO (3 ml) was added $K_2CO_3$ (191 mg, 1.38 mmol) (pH~9) followed by methyl 4-fluoro-3-nitrobenzoate (165 mg, 0.82 mmol). The reaction mixture was heated to 80 °C for 16 h (Monitored by TLC). Quenched with cold water and extracted with ethyl acetate (3 x 10 ml). The combined organic extracts were dried, evaporated and purified by flash column chromatography eluting with ethyl acetate in hexane (7:3) to get methyl 4-(4-((N-ethyl-7-(4-fluoro-N-isopropylbenzamido)heptanamido)methyl)piperidin-1-yl)-3-nitrobenzoate (**S1**) (100 mg, 26.17% yield) as a colorless sticky material and its corresponding hydrolyzed product acid 4-(4-((N-ethyl-7-(4-fluoro-N-isopropylbenzamido)heptanamido)methyl)piperidin-1-yl)-3-nitrobenzoic acid (**14**) which was further purified by RP-prep-HPLC using 0.1% formic acid in water and acetonitrile to get pure 4-(4-((N-ethyl-

7-(4-fluoro-N-isopropylbenzamido)heptanamido)methyl)piperidin-1-yl)-3-nitrobenzoic acid (**14**) (10 mg, 3% yield) as a colorless sticky material. LCMS: 599.4 $[M+H]^+$, HPLC: 99.72%, $^1$H NMR (400MHz, DMSO-d6) δ = 8.28 (s, 1H), 8.00 (d, $J$ = 8 Hz, 3sH), 7.35 (d, $J$ = 8.4 Hz, 3H), 3.84 (s, 6H), 3.02 (t, $J$ = 11.2 Hz, 5H), 2.24 (d, $J$ = 5.2 Hz, 6H), 1.82 (d, $J$ = 10.4 Hz, 5H), 1.69 (s, 3H), 1.22 (m, 6H), 0.96 (s, 4H).

**Synthesis of Methyl 3-amino-4-(4-((N-ethyl-7-(4-fluoro-N-isopropylbenzamido)heptanamido)methyl)piperidin-1-yl)benzoate (15)**

To a stirred solution of methyl 4-(4-((N-ethyl-7-(4-fluoro-N-isopropylbenzamido)heptanamido)methyl)piperidin-1-yl)-3-nitrobenzoate (**S1**) (100 mg, 0.167 mmol) in MeOH (3 ml), 10% Pd/C (40 mg, 50% moisture) was added. The reaction mixture was stirred at RT for 4 h (monitored by TLC) and catalyst was filtered out by using celite bed. The filtrate was evaporated under reduced pressure and residue was purified by RP-preparative HPLC using 0.1% formic acid in water and acetonitrile to get Methyl 3-amino-4-(4-((N-ethyl-7-(4-fluoro-N-isopropylbenzamido)heptanamido)methyl)piperidin-1-yl)benzoate (**15**) (50 mg, 52% yield) as a colorless sticky material. LCMS: 583.5 $[M+H]^+$, HPLC: 97.82%, $^1$H NMR (400MHz, DMSO-d6) δ = 7.45 (m, 2H), 7.32-7.18 (m, 4H), 6.93 (m, 1H), 4.92 (m, 2H), 3.78 (s, 3H), 3.34 - 3.15 (m, 10H), 2.34 (m, 2H), 1.83 (s, 1H), 1.70-1.55 (m, 5H), 1.41 - 1.34 (m, 6H), 1.25-1.01 (m, 10H). (Note: Acid proton not observed in $^1$H NMR).

#### Synthesis of (16) as described in Scheme S14

**Synthesis of N-(7-(((1-(4-bromo-2-nitrophenyl)piperidin-4-yl)methyl)(ethyl)amino)-7-oxoheptyl)-4-fluoro-N-isopropylbenzamide (16)**

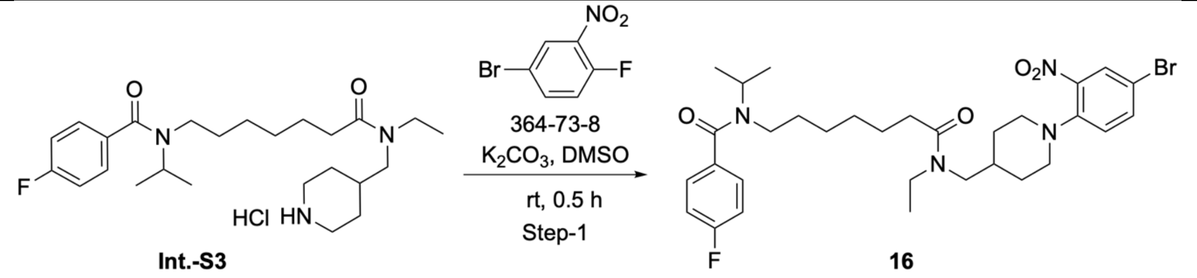

**Scheme S14**

To a solution N-(7-(ethyl(piperidin-4-ylmethyl)amino)-7-oxoheptyl)-4-fluoro-N-isopropylbenzamide hydrochloride (**Int.-S3 in Scheme S11**) (500 mg, 1.5 mmol) in DMSO (5 ml) was added $K_2CO_3$ (794 mg, 5.7 mmol) (pH~9) followed by 4-bromo-1-fluoro-2-nitrobenzene (502 mg, 2.3 mmol). The reaction mixture was stirred at RT for 0.5 h (Monitored by TLC). Quenched with cold water and extracted with ethyl acetate (3 x 10 ml). The combined organic extracts were dried, evaporated and purified by RP-prep-HPLC using 0.1% formic acid in water and acetonitrile to get N-(7-(((1-(4-bromo-2-nitrophenyl)piperidin-4-yl)methyl)(ethyl)amino)-7-oxoheptyl)-4-fluoro-N-isopropylbenzamide (**16**) (290 mg, 43% yield) as a yellow colored sticky material. LCMS: 635.2 $[M+H]^+$, HPLC: 99.16%, $^1$H NMR (400MHz, DMSO-d6) δ = 8.02 (m, 1H), 7.72 (m, 1H), 7.41 (m, 2H), 7.28 (t, $J$ = 8.4 Hz, 3H), 3.80 (m, 1H), 3.38 (m, 2H), 3.20 – 3.15 (m, 6H), 2.94 – 2.73 (m, 2H), 2.34 (m, 2H), 1.76 (s, 1H), 1.76-1.55 (m, 6H), 1.32-1.19 (m, 6H), 1.12-0.88 (m, 9H).

#### Copies of $^{1}$H-NMR spectra of compounds **1-16**

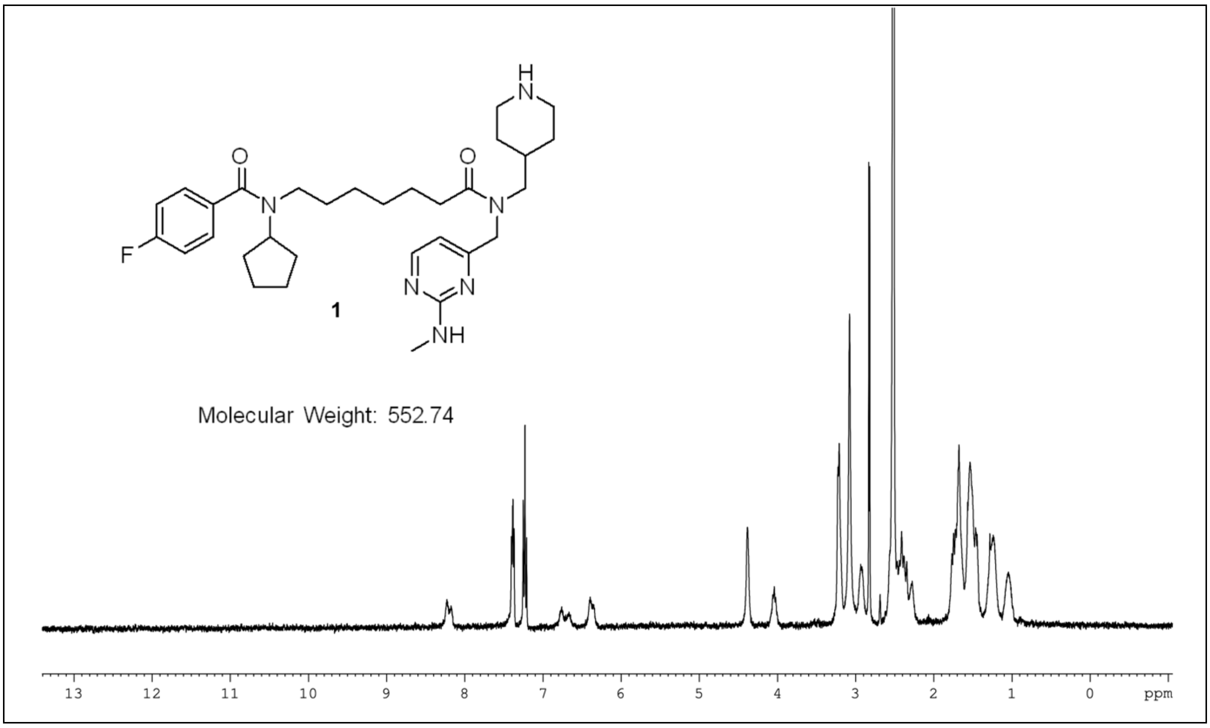

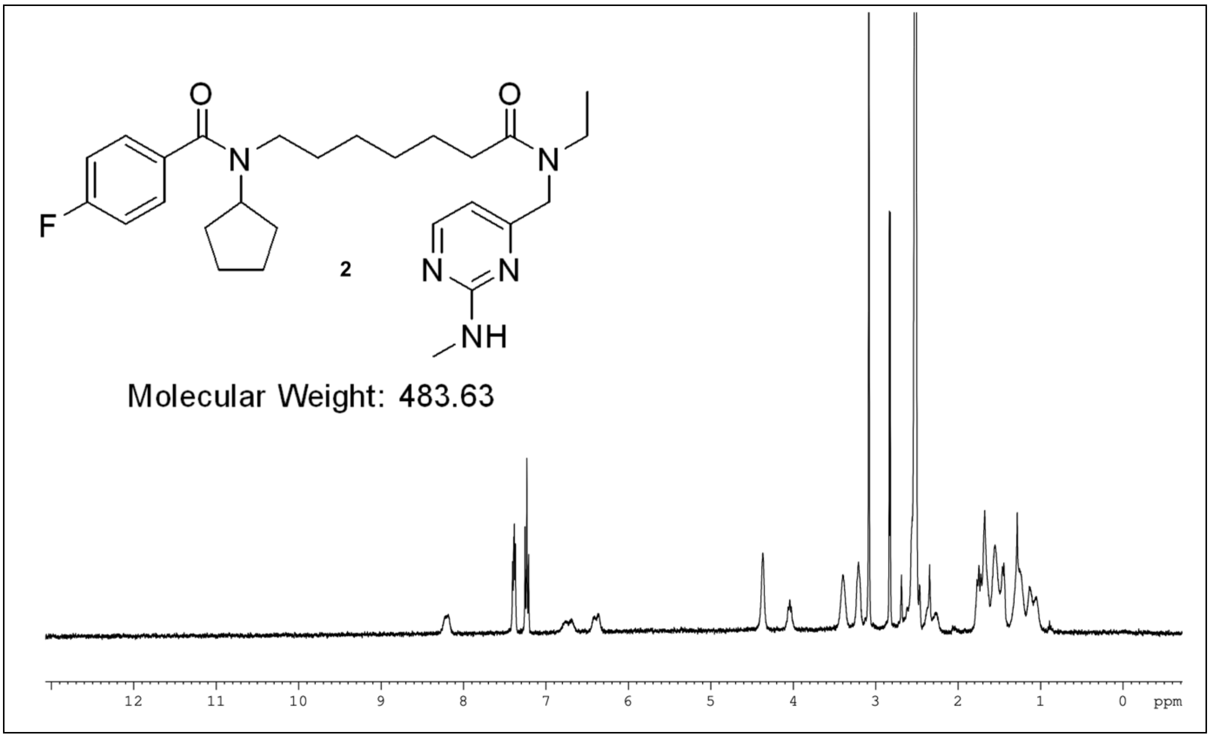

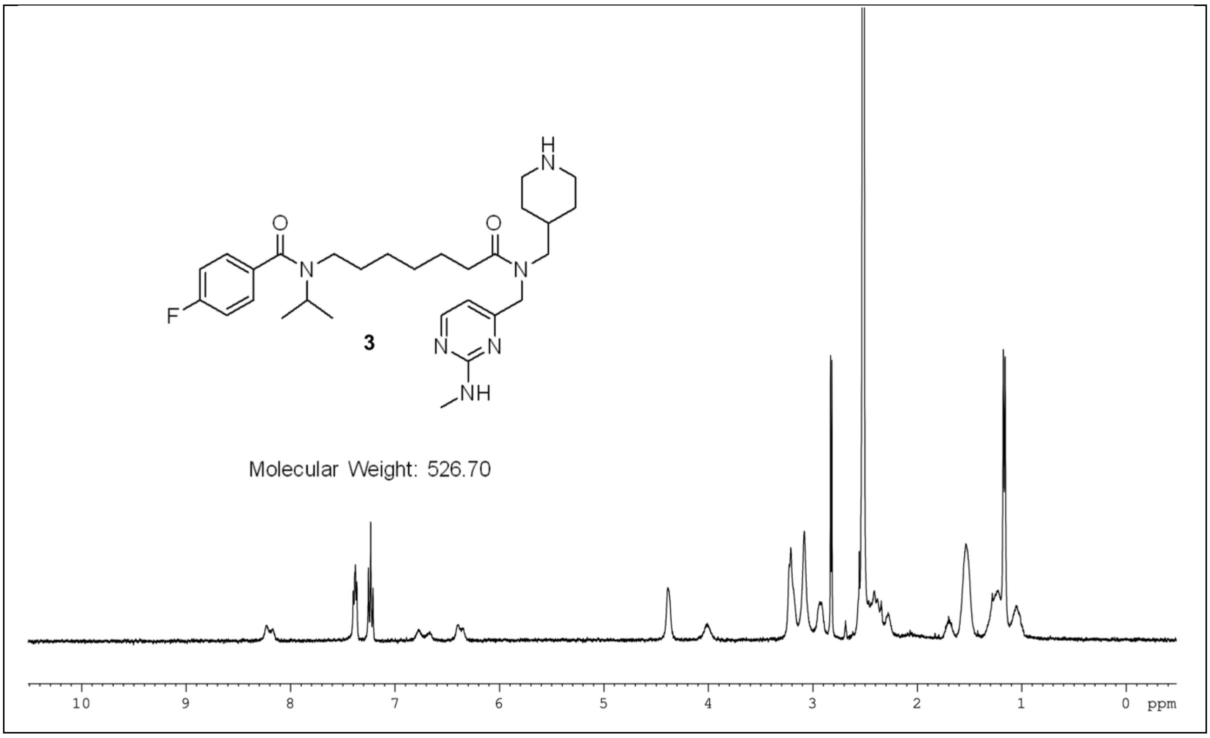
F
O
N
H
3
NH
Molecular Weight: 526.70
10 9 8 7 6 5 4 3 2 1 0 ppm

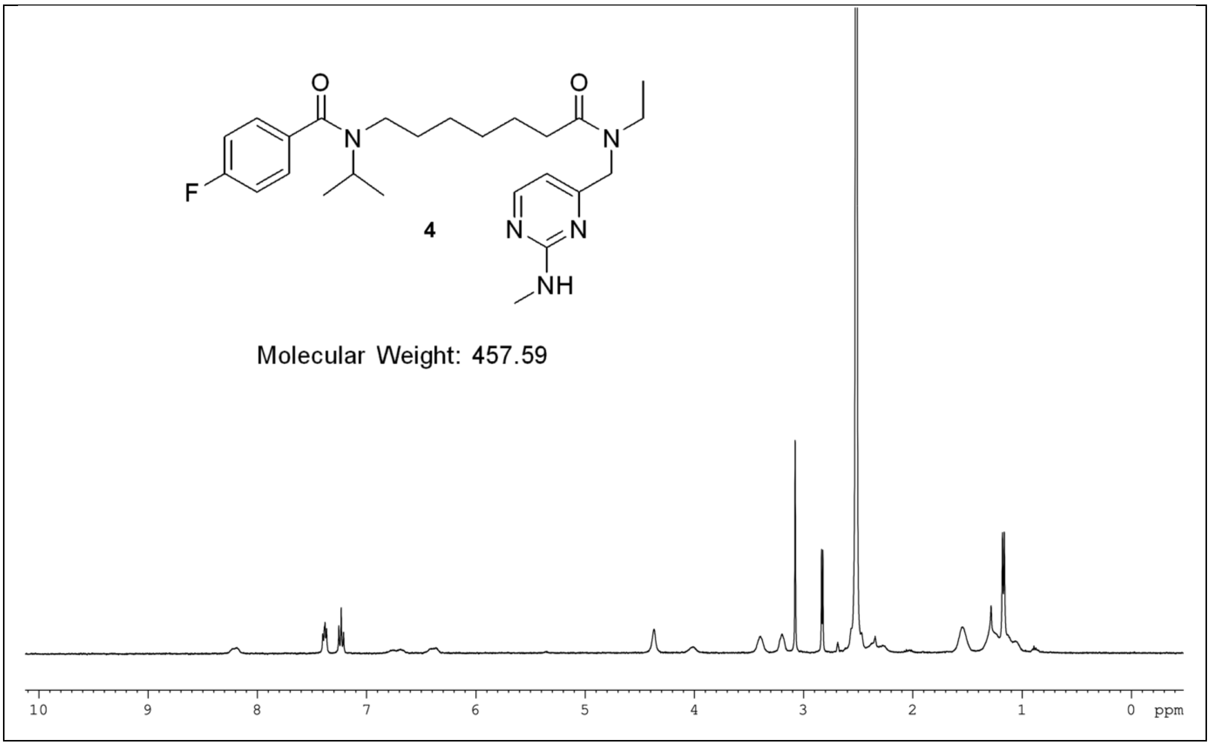
F
O
N
4
NH
Molecular Weight: 457.59
10 9 8 7 6 5 4 3 2 1 0 ppm

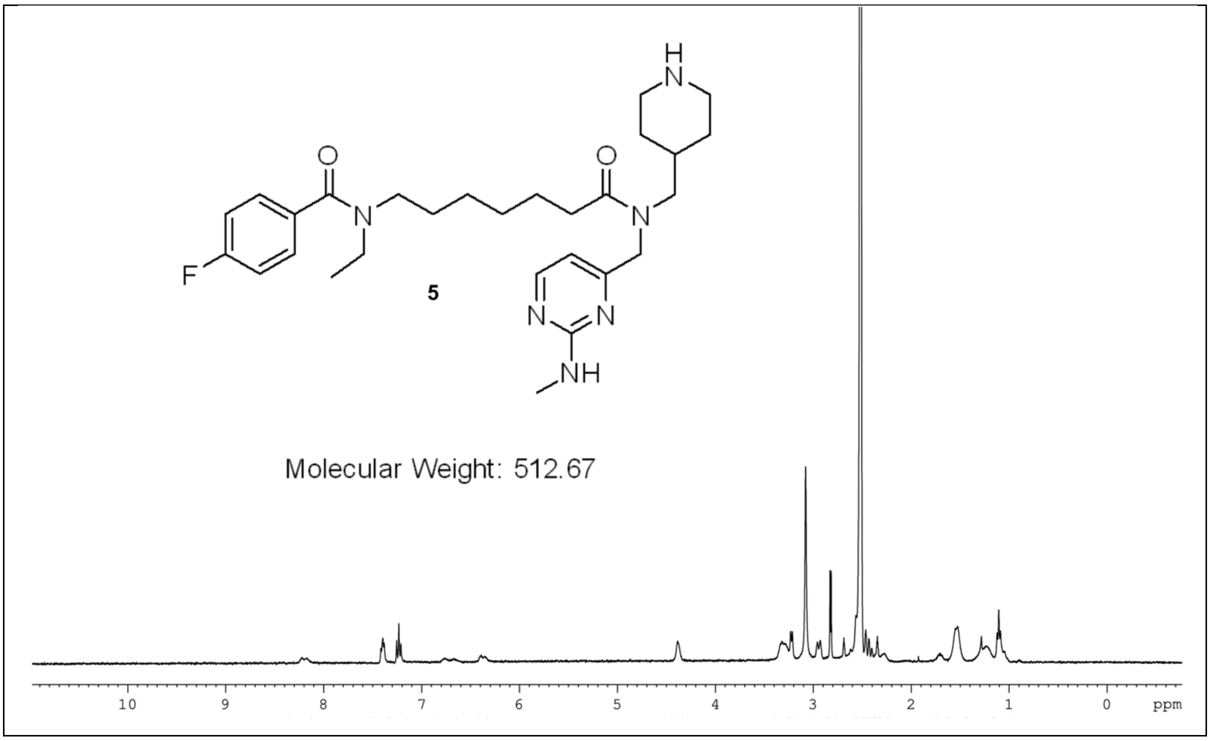
5
Molecular Weight: 512.67
10
9
8
7
6
5
4
3
2
1
0
ppm

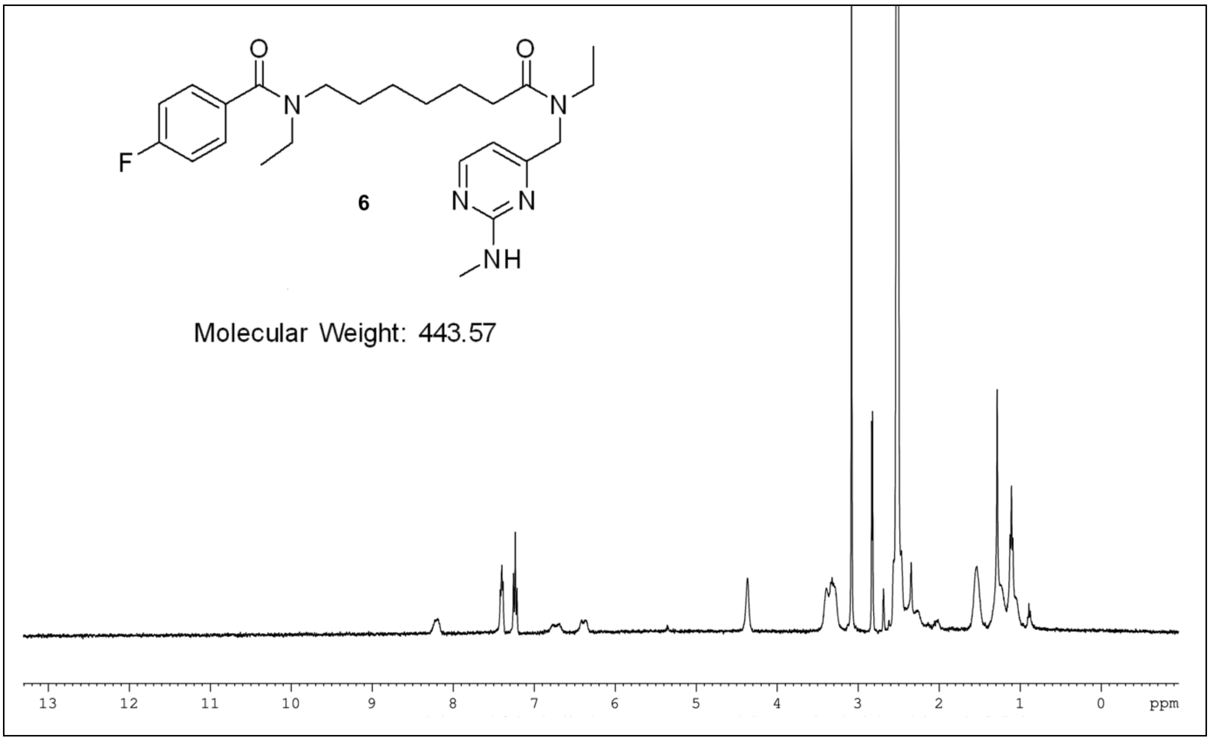
6
Molecular Weight: 443.57
13
12
11
10
9
8
7
6
5
4
3
2
1
0
ppm

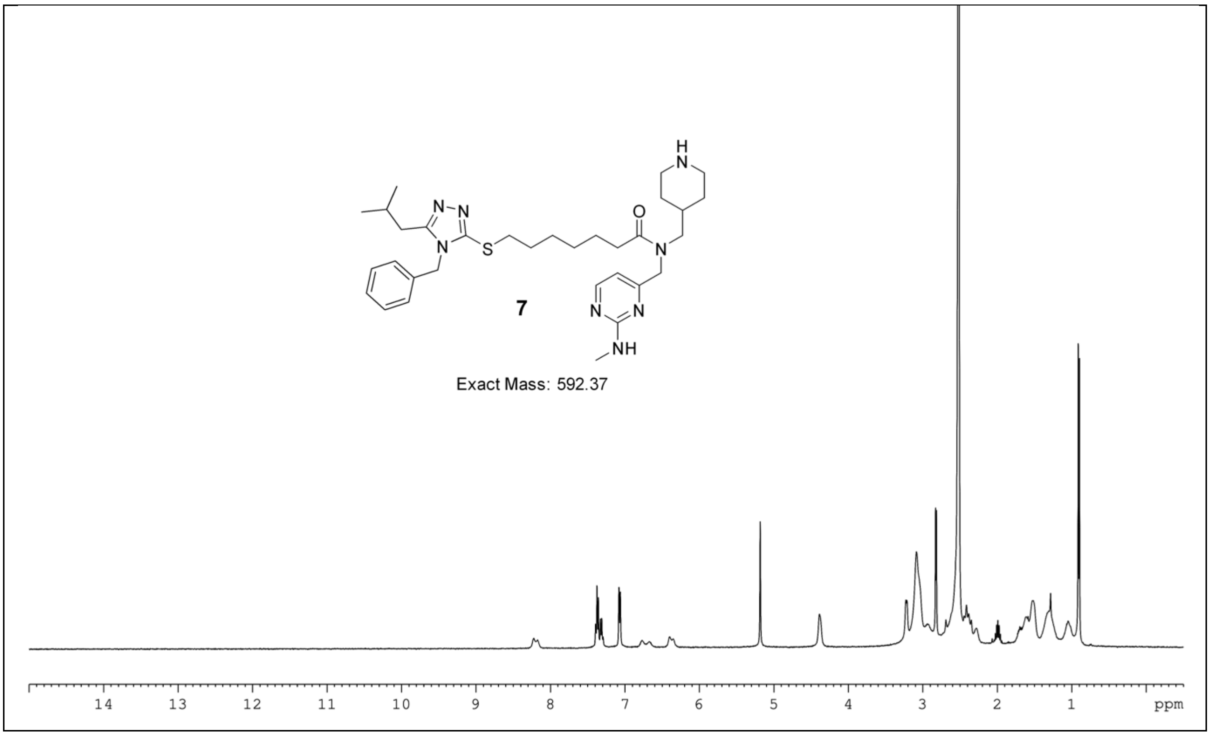

7
Exact Mass: 592.37
ppm

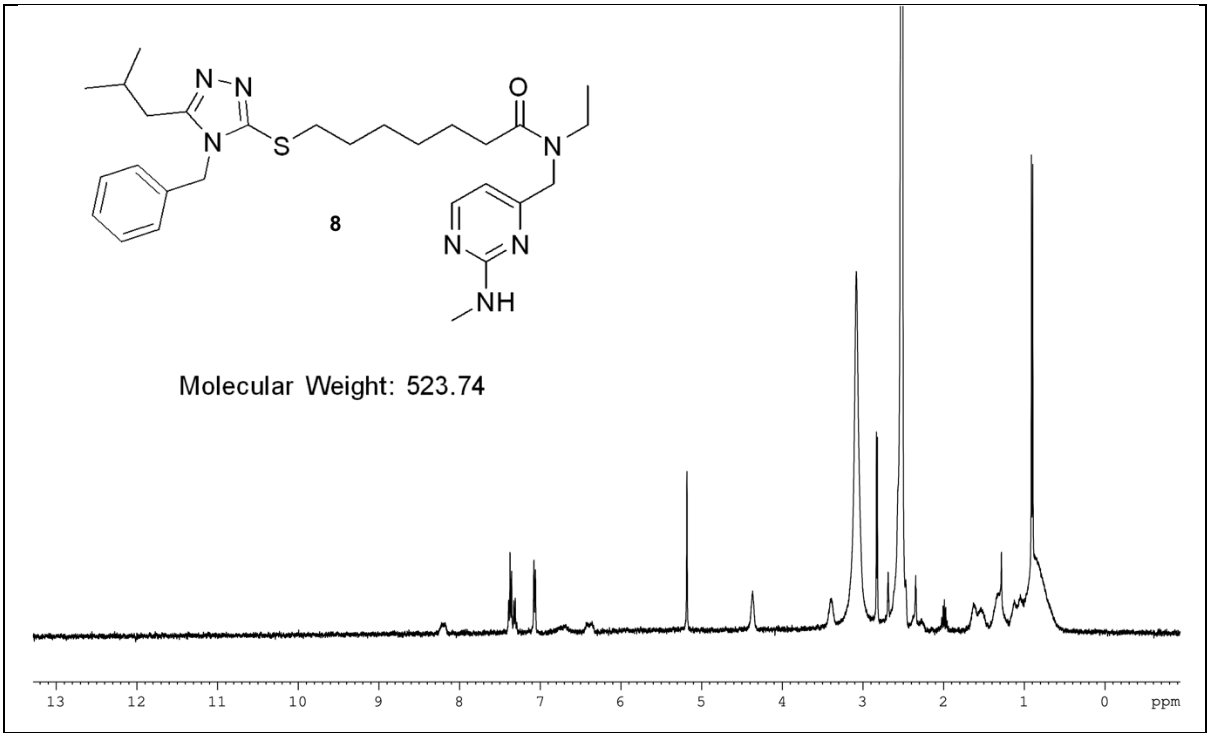

8
Molecular Weight: 523.74
ppm

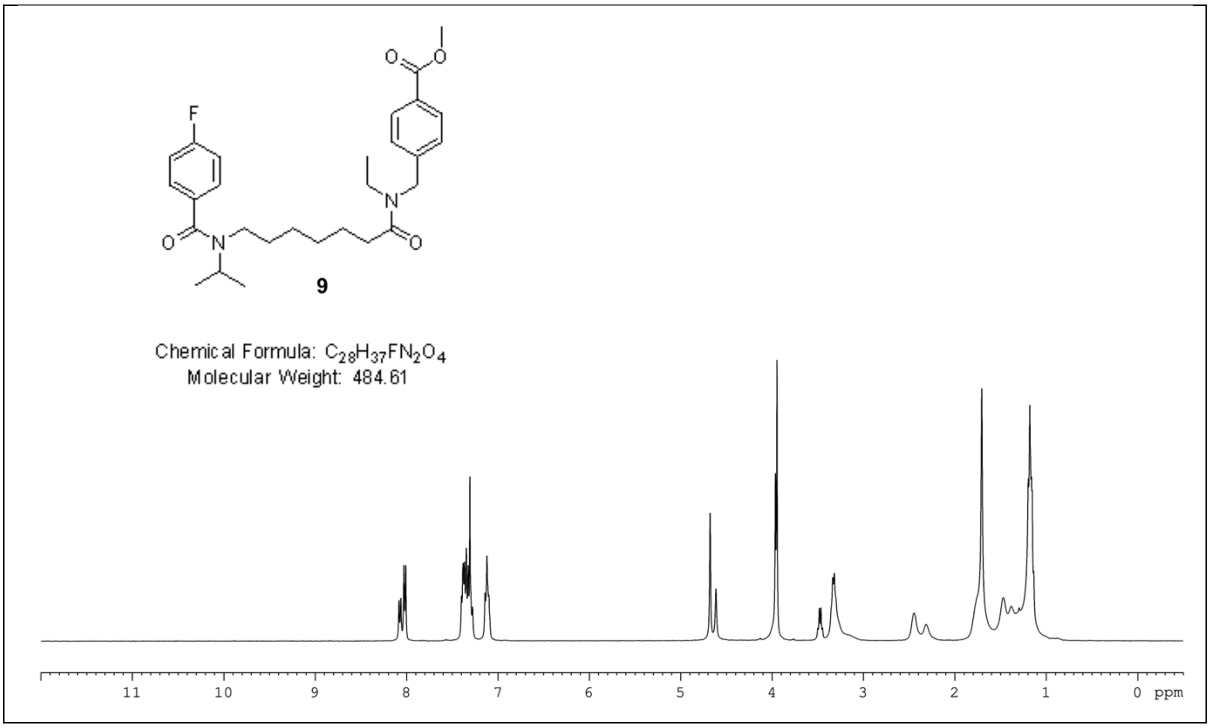
F
O
N
O
O
N
O
9
Chemical Formula: C28H37FN2O4
Molecular Weight: 484.61
11 10 9 8 7 6 5 4 3 2 1 0 ppm

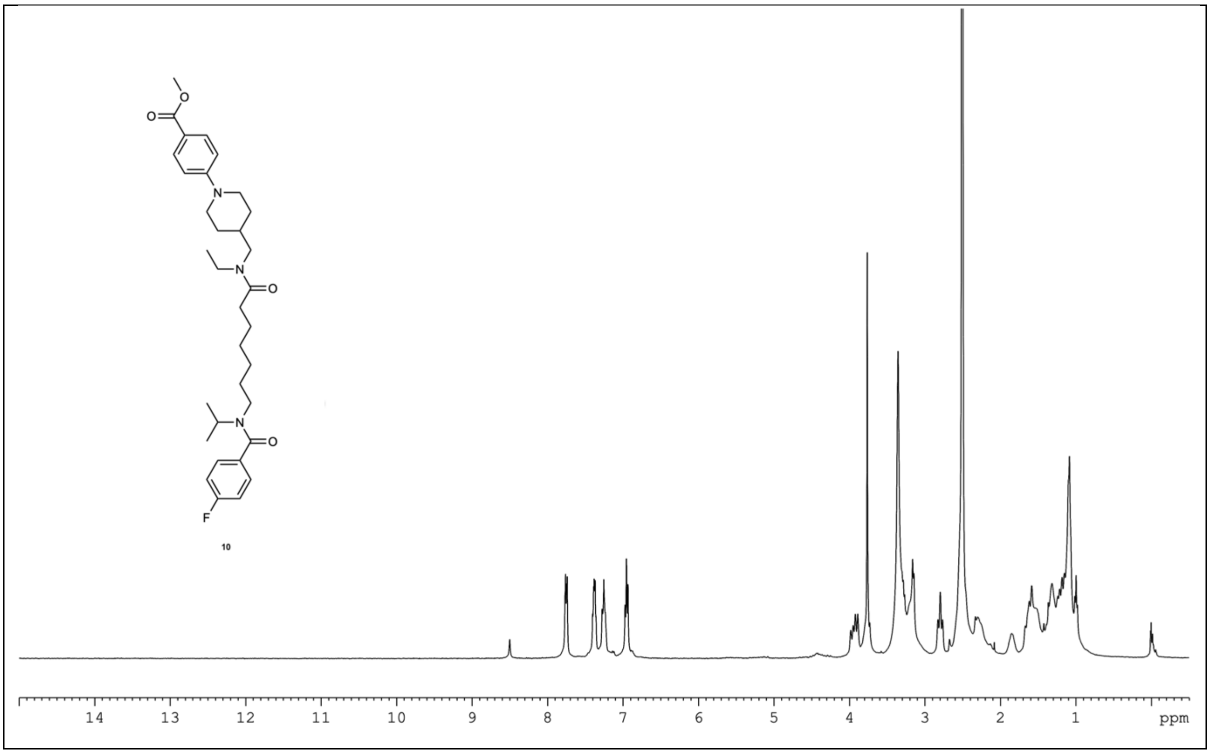
O
O
N
N
O
N
O
F
10
14 13 12 11 10 9 8 7 6 5 4 3 2 1 ppm

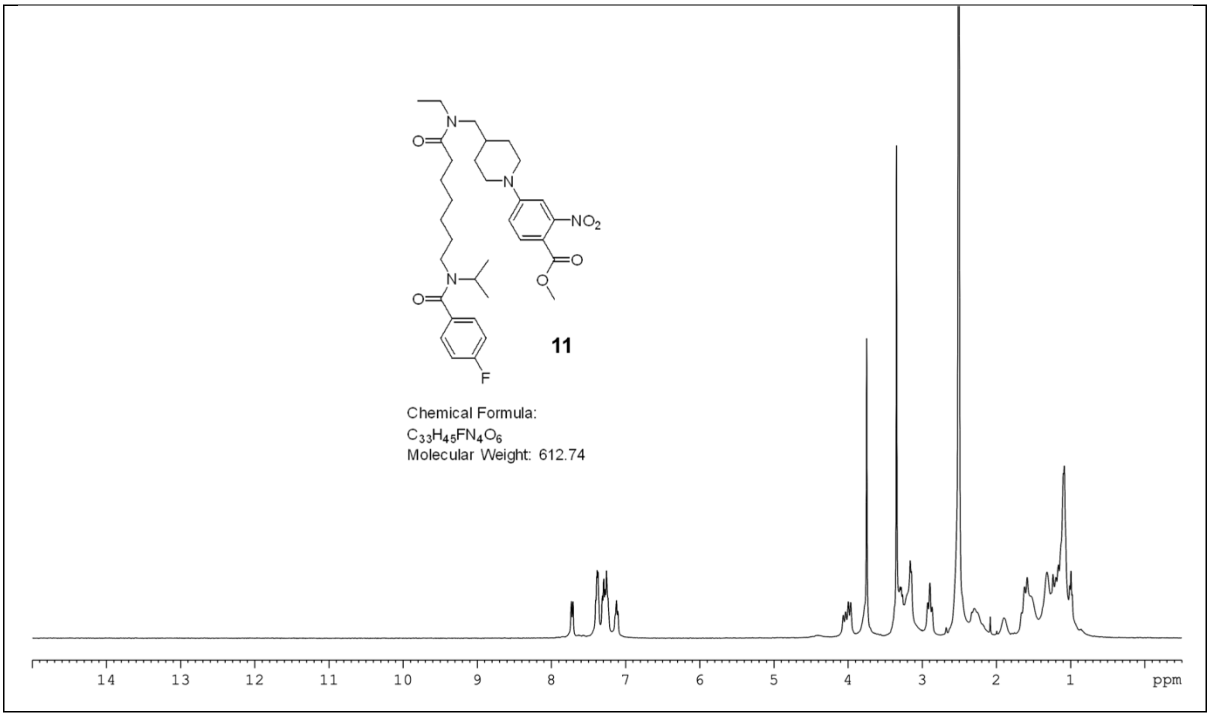
NO2
11
Chemical Formula:
C33H45FN4O6
Molecular Weight: 612.74
14 13 12 11 10 9 8 7 6 5 4 3 2 1 ppm

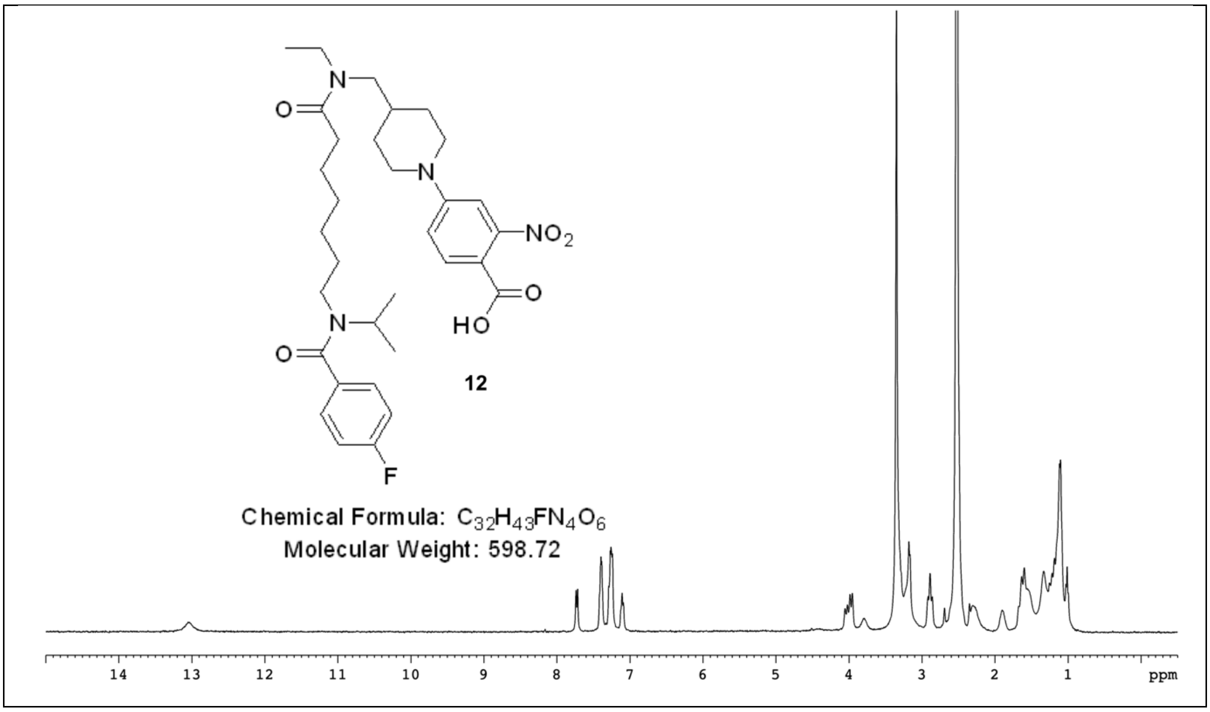
NO2
HO
12
Chemical Formula: C32H43FN4O6
Molecular Weight: 598.72
14 13 12 11 10 9 8 7 6 5 4 3 2 1 ppm

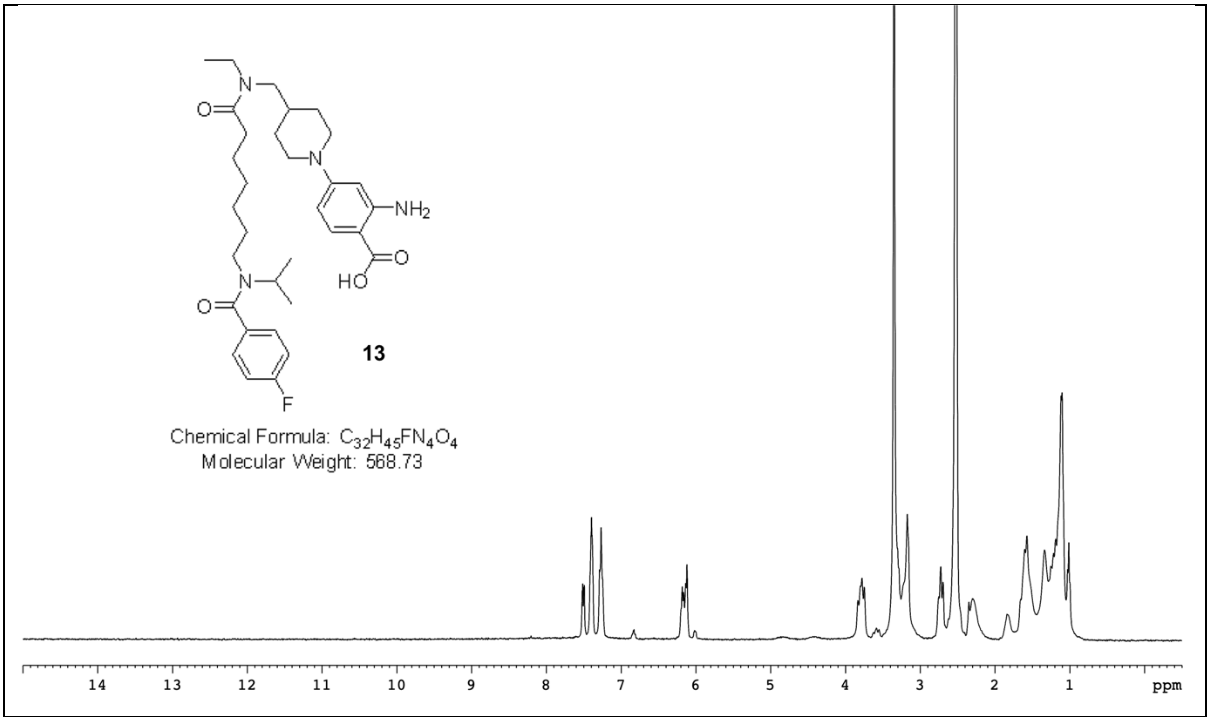
NH2
HO
O
N
F
13
Chemical Formula: C32H45FN4O4
Molecular Weight: 568.73
14 13 12 11 10 9 8 7 6 5 4 3 2 1 ppm

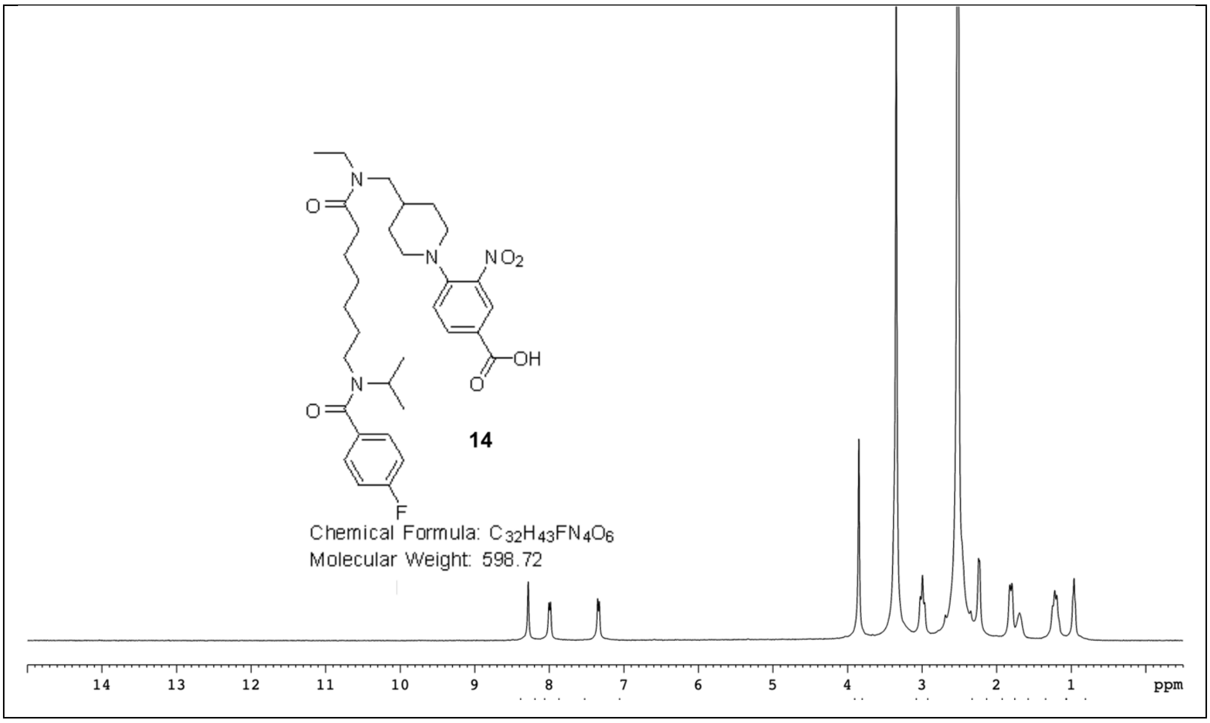
NO2
OH
O
N
F
14
Chemical Formula: C32H43FN4O6
Molecular Weight: 598.72
14 13 12 11 10 9 8 7 6 5 4 3 2 1 ppm

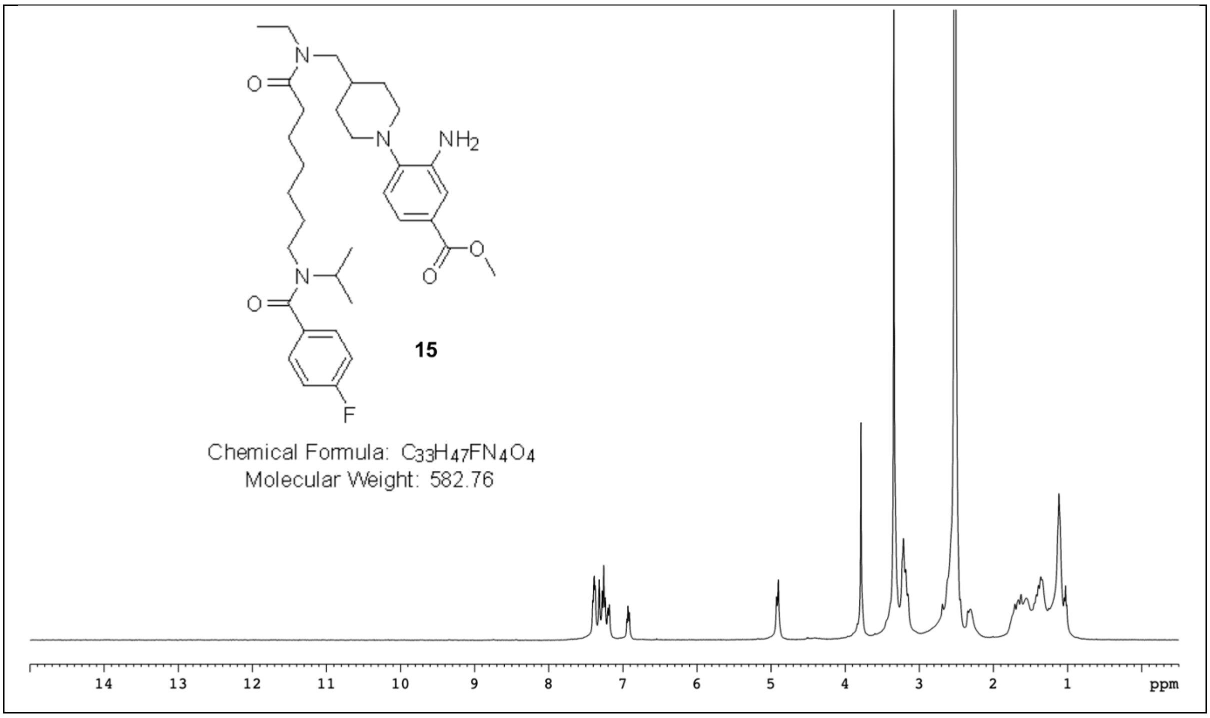

NH2
F
15
Chemical Formula: C33H47FN4O4
Molecular Weight: 582.76
14 13 12 11 10 9 8 7 6 5 4 3 2 1 ppm

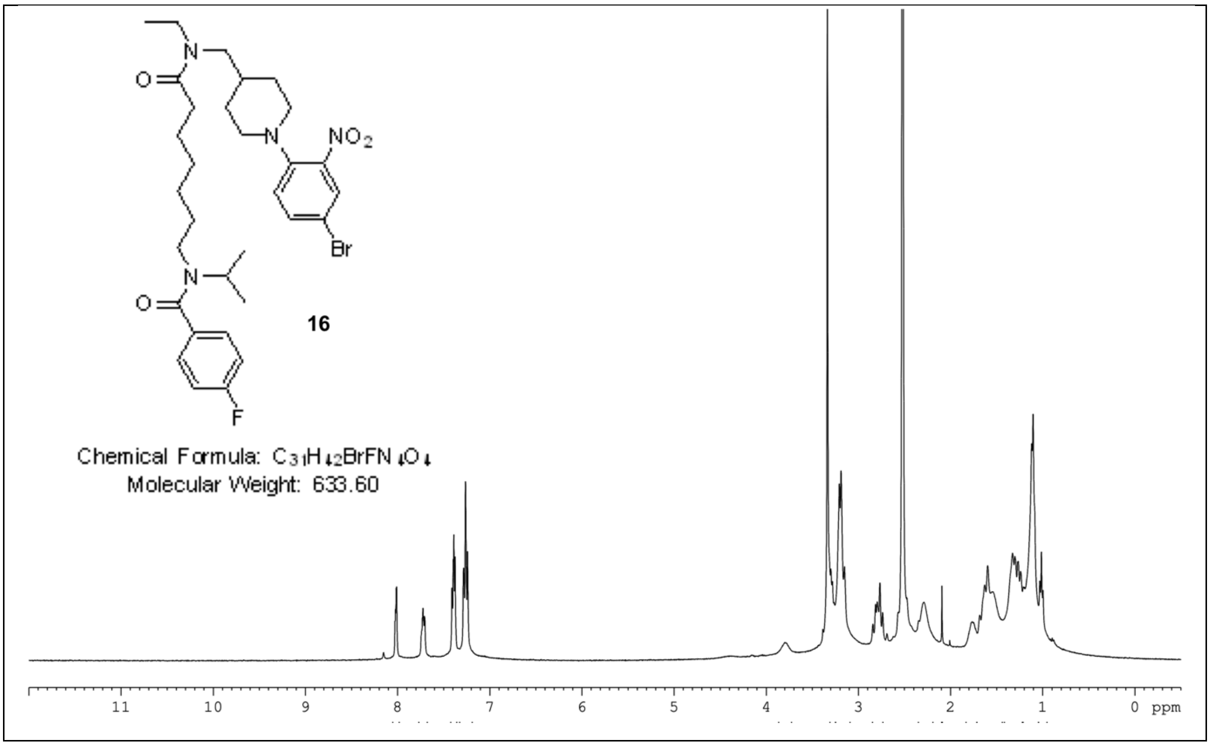

NO2
Br
F
16
Chemical Formula: C31H42BrFN4O4
Molecular Weight: 633.60
11 10 9 8 7 6 5 4 3 2 1 0 ppm
