## Supplementary material for "An improved PDE6D inhibitor combines with Sildenafil to synergistically inhibit KRAS mutant cancer cell growth": FigureS1

### Supplementary Information Figure S1

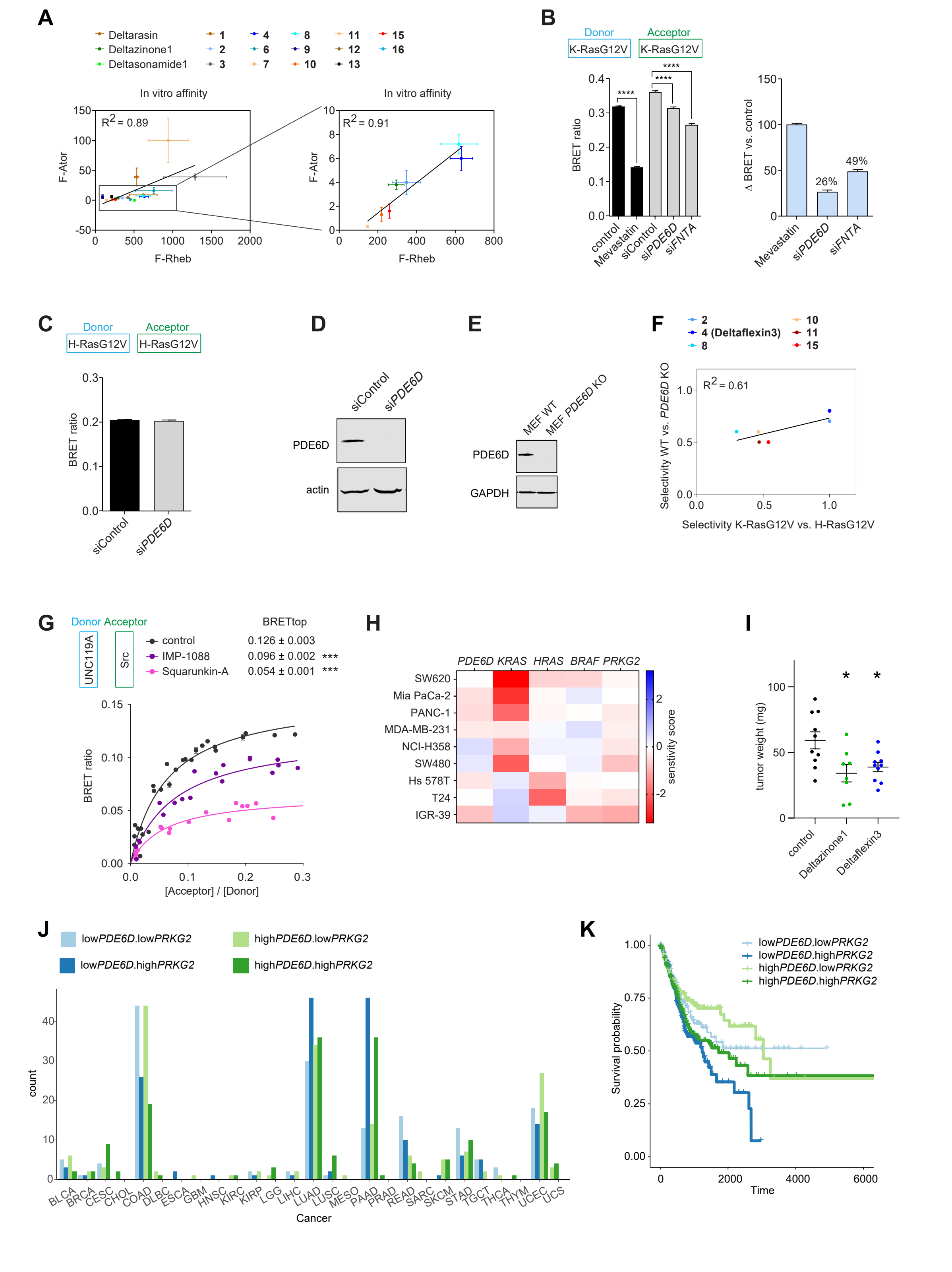


**Figure S 1. Supplementary data for the manuscript.**

(**A**) Correlation plots of inhibitor/ PDE6D-Kd values acquired using either F-Rheb or F-Ator as probes (**Data S2**). Data of compound **14** were excluded, due to its much lower affinity.

(**B**) K-RasG12V-membrane anchorage BRET is reduced by 5 µM Mevastatin treatment and knockdown of *PDE6D* or *FNTA* to different extents; n ≥ 3 (left). Derived from these data we prepared a plot of the loss of BRET ratio as compared to respective controls, by setting the loss of the Mevastatin treatment to 100% (right).
(**C**) H-RasG12V-membrane anchorage BRET is not affected by knockdown of *PDE6D*; n ≥ 3.
(**D**) Representative immunoblot data showing *PDE6D* knockdown efficiency in HEK293 EBNA cells; n ≥ 3. Immunoblot data showing *FNTA* knockdown efficiency in HEK293 EBNA cells were previously reported by us (Manoharan et al., 2023).
(**E**) Representative immunoblot data verifying *PDE6D* knockout in *PDE6D*-KO MEF cells.
(**F**) Correlation plot of K-RasG12V-BRET selectivity on the x-axis (**Fig. 2B**) vs. WT/KO-viability data derived PDE6D-selectivity (**Fig. 3A**).
(**G**) BRET-titration curves of the UNC119A/ Src complex after treatment with 5 µM of the N-myristoyl-transferase inhibitor IMP-1088 or 5 µM the UNC119-inhibitor Squarunkin A; n ≥ 3. Statistical comparisons of BRETtop values to controls were done using two-tailed Student's t-test.
(**H**) Heatmap of the ATARiS-sensitivity scores of selected genes for all cancer cell lines used in this study. Negative values indicate a decrease of proliferation upon gene knockdown and therefore a higher dependency of the cell line on that gene.
(**I**) Weights of MDA-MB-231 derived microtumors (≥ 8 per condition from n = 3) from CAM assays after treatment with 10 µM of indicated compounds.
(**J**) Number of *KRAS*-mutant patient tumor samples with indicated high or low gene expression level combinations of *PDE6D* and *PRKG2* (gene for PKG2) by cancer type (TCGA study abbreviations).
(**K**) Overall survival of *KRAS*-mutant patient tumor samples from TCGA with indicated high or low gene expression level combinations of *PDE6D* and *PRKG2* (gene for PKG2). Number of patients per group: 157 (low*PDE6D*.low*PRKG2*), 165 (low*PDE6D*.high*PRKG2*), 166 (high*PDE6D*.low*PRKG2*), 159 (high*PDE6D*.high*PRKG2*). Kaplan-Meyer test for the difference between low*PDE6D*.high*PRKG2* and high*PDE6D*.low*PRKG2* groups: p = 0.0006.
